# Population size rescaling significantly biases outcomes of forward-in-time population genetic simulations

**DOI:** 10.1101/2024.04.07.588318

**Authors:** Amjad Dabi, Daniel R. Schrider

**Affiliations:** Department of Genetics, University of North Carolina, Chapel Hill, North Carolina, USA Address: 120 Mason Farm Rd, Chapel Hill, NC 27599

**Keywords:** forward simulation, rescaling, approximation error

## Abstract

Simulations are an essential tool in all areas of population genetic research, used in tasks such as the validation of theoretical analysis and the study of complex evolutionary models. Forward-in-time simulations are especially flexible, allowing for various types of natural selection, complex genetic architectures, and non-Wright-Fisher dynamics. However, their intense computational requirements can be prohibitive to simulating large populations and genomes. A popular method to alleviate this burden is to scale down the population size by some scaling factor while scaling up the mutation rate, selection coefficients, and recombination rate by the same factor. However, this rescaling approach may in some cases bias simulation results. To investigate the manner and degree to which rescaling impacts simulation outcomes, we carried out simulations with different demographic histories and distributions of fitness effects using several values of the rescaling factor, *Q*, and compared the deviation of key outcomes (fixation times, allele frequencies, linkage disequilibrium, and the fraction of mutations that fix during the simulation) between the scaled and unscaled simulations. Our results indicate that scaling introduces substantial biases to each of these measured outcomes, even at small values of *Q*. Moreover, the nature of these effects depends on the evolutionary model and scaling factor being examined. While increasing the scaling factor tends to increase the observed biases, this relationship is not always straightforward, thus it may be difficult to know the impact of scaling on simulation outcomes *a priori*. However, it appears that for most models, only a small number of replicates was needed to accurately quantify the bias produced by rescaling for a given *Q*. In summary, while rescaling forward-in-time simulations may be necessary in many cases, researchers should be aware of the rescaling procedure’s impact on simulation outcomes and consider investigating its magnitude in smaller scale simulations of the desired model(s) before selecting an appropriate value of *Q*.

**ARTICLE SUMMARY:** Computer simulations are an indispensable tool for researchers studying the dynamics of population genetic models, but often require large computing resources and long run times. A widely used approximation alleviates these computational requirements by simulating a smaller population and increasing the values of other model parameters to compensate. However, this results in significant biases for important simulation outcomes. Fortunately, the direction and magnitude of these biases can be measured beforehand by carrying out a relatively small number of simulations.

## INTRODUCTION

Stochastic simulations are widely used in population genetics (Hoban et al. 2012; Peng et al. 2015; Adrion et al. 2020). Their uses include the study of the behavior of a wide variety of population genetic models (either when used on their own in simulation studies (Hudson 1983a; Charlesworth et al. 1993; Hermisson and Pennings 2005; Orr and Unckless 2008; Peischl et al. 2013; Zeng 2013) or when used to validate theoretical predictions (Morgan and Strobeck 1979; Braverman et al. 1995; Thornton 2019; Battey et al. 2020; Schrider 2020; Torres et al. 2020)), the comparison of simulations to data to infer model parameters or choose between competing models (Tavare et al. 1997; Pritchard et al. 1999; Beaumont et al. 2002; Beaumont 2010; Csilléry et al. 2010), assessments of the accuracy of inference methods (Pavlidis et al. 2013; Alachiotis and Pavlidis 2018; Galimberti et al. 2020), and the generation of training datasets for machine learning methods (Pavlidis et al. 2010; Ronen et al. 2013; Gao et al. 2016; Sheehan and Song 2016; Chan et al. 2018; Schrider and Kern 2018; Flagel et al. 2019; Mughal and DeGiorgio 2019; Caravagna et al. 2020; Sanchez et al. 2021; Smith et al. 2023). Given their wide range of research applications, considerable effort has been made to improve the accessibility, computational speed, and flexibility of population genetic simulations (Hudson 2002; Ewing and Hermisson 2010; Messer 2013; Kern and Schrider 2016; Kelleher et al. 2018; Haller and Messer 2023).

Two main modes of population genetic simulation are available to researchers: coalescent simulations, which trace the genealogy of a sample of individuals back in time until all segments of the chromosome have reached the sample’s MRCA, and forward-in-time simulations, which explicitly simulate an entire population of chromosomes evolving under the desired model. Coalescent simulations have been a mainstay of population genetics for the last few decades because of their computational efficiency (Liang et al. 2007; Carvajal-Rodriguez 2008; Kelleher et al. 2016), as they are often used to simulate only a relatively small sample of genomes rather than the entire population (Hudson 1983b), and jump backward from one event (e.g., coalescence and recombination) to the next rather than simulating each generation of the evolutionary scenario.

While computationally efficient, coalescent simulations are relatively limited in the evolutionary scenarios that they can model, especially with respect to natural selection, and are therefore more commonly used for modeling demographic histories (see Ewing and Hermisson (2010); Kern and Schrider (2016); Baumdicker et al. (2022); Teshima and Innan (2009) for notable exceptions). Forward-in-time simulations, on the other hand, provide much greater flexibility (Peng and Kimmel 2005; Hernandez 2008; Neuenschwander et al. 2008; Thornton 2014; Haller and Messer 2017), giving users the ability to simulate more complex models involving not only demographic changes, but several types of natural selection, and scenarios that fall outside the bounds of the Wright-Fisher model (Haller and Messer 2019). Unfortunately, forward-in-time simulations can require a large amount of computational power and memory because the entire population is simulated for each generation of the desired scenario, rendering simulations of large populations and/or very long histories infeasible. One popular approach to alleviate this computational burden is to rescale forward-in-time simulations such that a smaller population is used to capture the desired evolutionary dynamics that would be expected for the larger population (Uricchio and Hernandez 2014). Typically, this is done by scaling down the population size by some factor (*Q*) while scaling up the mutation and recombination rates by the same factor (Kim and Wiehe 2009). Theoretically, this scaling approximation would preserve important population-level measures and allow easy recovery of the times at which events occurred during the simulation. For instance, the population-scaled mutation rate, θ, for a population of size *N* and with a mutation rate of μ, is equal to 4*N*μ. After rescaling, our new population size is *N*/*Q* and our new mutation rate is μ × *Q*, and thus θ is unchanged:

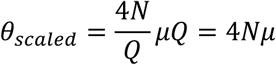

Meanwhile, under the standard neutral model, the expected time to the most recent common ancestor (TMRCA) between any two individuals in the unscaled simulation is 2*N* generations. Because the population size is decreased by a factor of *Q* in the scaled simulations, the expected TMRCA becomes 2*N*/*Q*. Consequently, to produce genealogies of the desired shape and size, the evolutionary model being simulated must be modified such that the time intervals between events specified in the simulated model (e.g., population splits, migration events, etc.) are divided by *Q*. This modification includes any applicable burn-in times and the total run time of the simulation itself. Therefore, scaling results in simulating smaller populations for shorter periods of time and thus substantially reducing computation time—in principle a quadratic speedup (Kim and Wiehe 2009). Following the simulation, if one wishes to investigate the timescale of any stochastic events occurring during the simulation such as fixations, coalescent events between particular pairs of individuals, etc., the times of these events can simply be multiplied by *Q* to produce results that approximate those expected under the original unscaled model.

This population-size “rescaling trick” has been used in forward simulations for decades, with its first use dating all the way back to Hill and Robertson (1966). However, the consequences of rescaling have rarely been evaluated, and with the exception of Hill and Robertson’s initial comparison of the effects of changing *N* from 16 to 8, those few studies that have examined the impact of rescaling have noticed a significant effect. For example, although the approximations underlying rescaling appear to be sound at least for simulations lacking natural selection, Adrion et al. (2020) found that in some cases, rescaling by modest values of *Q* can bias the outcomes of purely neutral simulations, resulting in biased estimates of summaries of diversity (π (Nei and Li 1979), and Tajima’s *D* (Tajima 1989)), and linkage disequilibrium (LD) as measured by *r*^2^. Moreover, when natural selection is included, rescaling may introduce further biases. The selection coefficient is often rescaled by multiplying it by *Q*, such that *Ns*, a commonly used measure of the efficacy of selection, is unchanged (Uricchio and Hernandez 2014; Johri et al. 2020; Schrider 2020). However, the time to fixation of a selected mutation does not scale linearly with *N*. Indeed, Uricchio and Hernandez (2014) showed that, under a recurrent hitchhiking model, rescaling blunts the impact of selection on diversity levels and the site frequency spectrum. Furthermore, Comeron and Kreitman (2002) showed that rescaling alters the dynamics of interference selection for weakly selected mutations, affecting important measures such as Tajima’s *D*. Although these studies have examined a very limited set of scenarios, their findings suggest that undesirable effects of rescaling could be commonplace. This possibility, combined with the increasing importance and use of forward simulations, underscores the need to investigate the impact of rescaling across a larger array of simulated models. We still know little about the manner in which simulations are affected (i.e., which simulation outcomes are upwardly or downwardly biased by rescaling), how widespread these effects are across simulated models, and how the severity of these effects varies with *Q*.

In this study, we carry out both scaled and unscaled simulations under a variety of demographic and selective scenarios and investigate the impact of rescaling on important simulation outcomes such as fixation times, site frequency spectra, and linkage disequilibrium. Our simulations explore various selection models containing different combinations of neutral, deleterious, and beneficial mutations, including simulations with a distribution of fitness effects estimated from human population genomic data. We also examine scaling effects in constant, expanding, and contracting populations. To quantify the effect of scaling, we train classification models that use a set of simulation outcomes as features (i.e., input data to machine learning models) to determine whether the results of scaled and unscaled simulations are distinguishable across increasing values of *Q*. Furthermore, for each feature we characterize the direction and magnitude of the deviation between scaled and unscaled simulations to provide a detailed picture of the effects of scaling across our scenarios. Our results show that rescaling simulations can result in a substantial deviation of numerous simulation outcomes, and that the magnitude of these effects usually increases with the value of *Q*. We also find that the choice of demographic model can have an impact on the severity of the rescaling effect. Overall, the results indicate the need to exercise caution in making inferences from rescaled simulations, especially when large values of *Q* are used.

## METHODS

### Simulation and rescaling strategy

We used `SLiM 4.0.1` (Haller and Messer 2023) to carry out scaled and unscaled forward-in-time population genetic simulations of a variety of demographic histories, genome architectures, and strengths/types of selection (Table 1). All our simulations conform to Wright-Fisher assumptions: generations are non-overlapping, mating is random, and all individuals are hermaphroditic, and the population size is solely determined by the specified demographic model and is not affected by the population’s fitness. For our simulations of *Drosophila* populations (bottom two rows of Table 1) we used `stdpopsim 0.2.0` (Adrion et al. 2020; Lauterbur et al. 2023), which contains a catalog for multiple species and produces `SLiM` scrips with the correct parameters, to generate a template SLiM script which we then modified. This strategy streamlined the process of faithfully producing simulation scripts with realistic *Drosophila* parameters. For our main simulations, the mutation and recombination rates are both *Q* × 1.2 × 10(N/Q) per base pair, the genome size is 25 Mb, and the scaled population size was set to *N*/*Q*, with *N* = 10,000. For mutations with a constant population size, a burn-in was performed for a duration of 10 × (N/Q) generations, after which statistics were recorded during and after a simulation run of 4 × (N/Q) generations. We also performed simulations of the full model in the presence of a single-event population expansion or contraction. In the population expansion simulations, the unscaled population size expands from *N_anc_* = 10,000/*Q* individuals to *N*_0_= 20,000/*Q* individuals after the burn-in, while in the population contraction simulations the population size decreases from *N_anc_* = 10,000/*Q* individuals to *N*_0_ = 5,000/*Q* individuals after the burn-in. In both demographic scenarios, the burn-in runs for 10 × *N_anc_* generations, followed by *N*_0_ generations after the population size change. For example, an unscaled simulation under the population expansion model would run for 100,000 generations of burn-in followed by 20,000 generations, during and after which statistics summarizing the simulation results are recorded. For each simulated scenario, we generated 1000 replicates for each scaling factor (*Q*) except for the no-beneficials larger population model described below, where the computational demand necessitated a smaller sample of 100 replicates.

**Table 1:** Summary of Models Simulated.

| Model Name | Model Description |
| --- | --- |
| Full model | Based on (Boyko et al. 2008) distribution of fitness effects. Contains neutral, beneficial, and deleterious mutations. |
| No-beneficials model | Same as the full model but contains no beneficial mutations. |
| No-beneficials sparse model | Same as the no-beneficials model, but with a lower fraction of deleterious mutations. |
| No-beneficials larger population model | Same as the no-beneficials mode but with double the population size. |
| Strictly neutral model | Same as the full model but with neutral mutations only. |
| Population expansion model | Same as the full model but with a doubling of the population size after burn-in. |
| Population contraction model | Same as the full model but with a halving of the population size after burn-in. |
| Sweep model | Simulated a sweep of a single beneficial mutation ($s=0.05$). Mutation and recombination rates are same as full mode. |
| *Drosophila* model | Based on the `stdpopsim` catalog of the species *Drosophila Melanogaster* and includes neutral, beneficial, and deleterious mutations. |
| *Drosophila* stronger-selection model | Same as the *Drosophila* model but with a higher selection coefficient for beneficial mutations. |

The fitness effects of mutations in our simulations were as specified in Table 2, and used the gamma distribution of deleterious fitness effects estimated from African American genomes by Boyko et al. 2008. We note that for mutations with fitness effects drawn from a Gamma distribution, the scaling is done by multiplying the mean of the distribution by *Q* and keeping the shape parameter fixed as this has the same effect as multiplying every value drawn from the unscaled distribution by *Q*. The gamma distribution specified in Table 2 is of the form ∼*Gamma*(*mean*, *shape*). Using this general structure, we performed scaled and unscaled simulations of several different scenarios each with different mixes of neutral, deleterious, and beneficial mutations (Table 3). Note that the parameter values listed in the second column of Table 3 specify the probability that any new mutation will be of a given mutation type as follows: *f*_n,_ is the probability that a new mutation will be neutral, *f*_a._ is the probability it will be beneficial, and *f*_n_ is the probability it will be deleterious. All individuals in the simulations are diploid, and the dominance coefficient for all mutations is 0.5.

**Table 2:** Mutation parameters for simulations.

| Mutation Type | Fitness Effect (s) |
| --- | --- |
| Neutral | 0 |
| Beneficial | $0.05 \times Q$ |
| Deleterious | $\sim \text{Gamma}(-0.0294 \times Q, 0.184)$ |

**Table 3:**
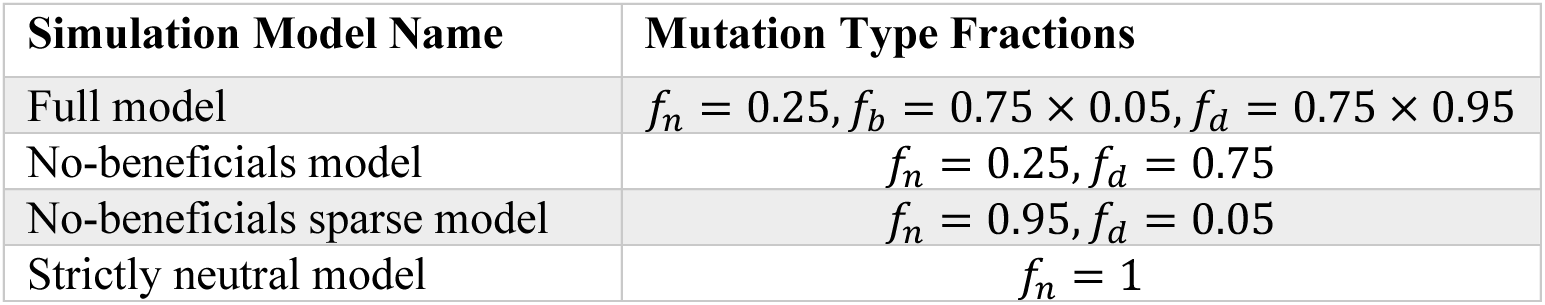
Proportions of neutral, deleterious, and beneficial mutations in each simulation model.

| Simulation Model Name | Mutation Type Fractions |
| --- | --- |
| Full model | $f_n = 0.25, f_b = 0.75 \times 0.05, f_d = 0.75 \times 0.95$ |
| No-beneficials model | $f_n = 0.25, f_d = 0.75$ |
| No-beneficials sparse model | $f_n = 0.95, f_d = 0.05$ |
| Strictly neutral model | $f_n = 1$ |

**Table 4.**
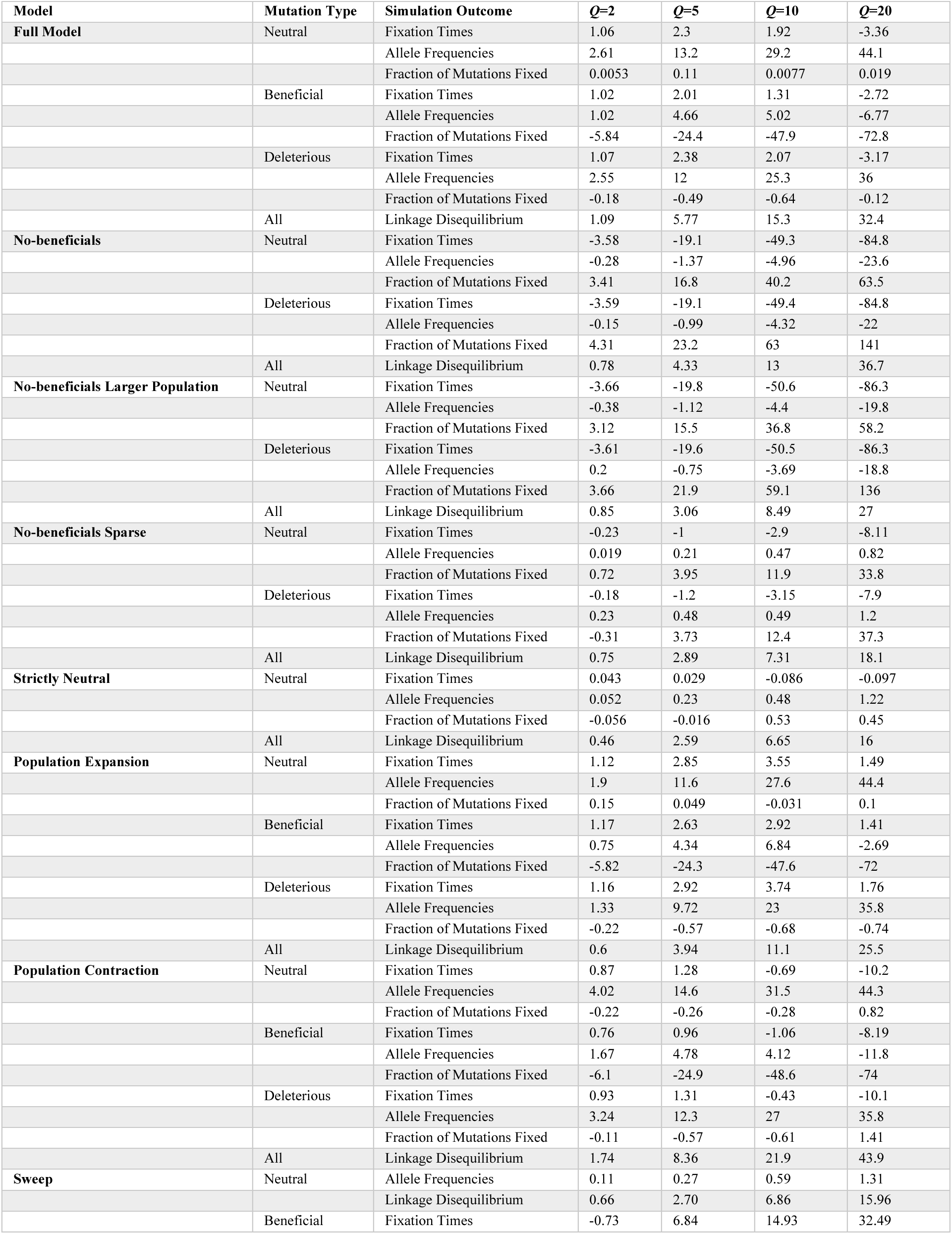
Summary of MPE Results for non-*Drosophila* Models.

**Table 5.**
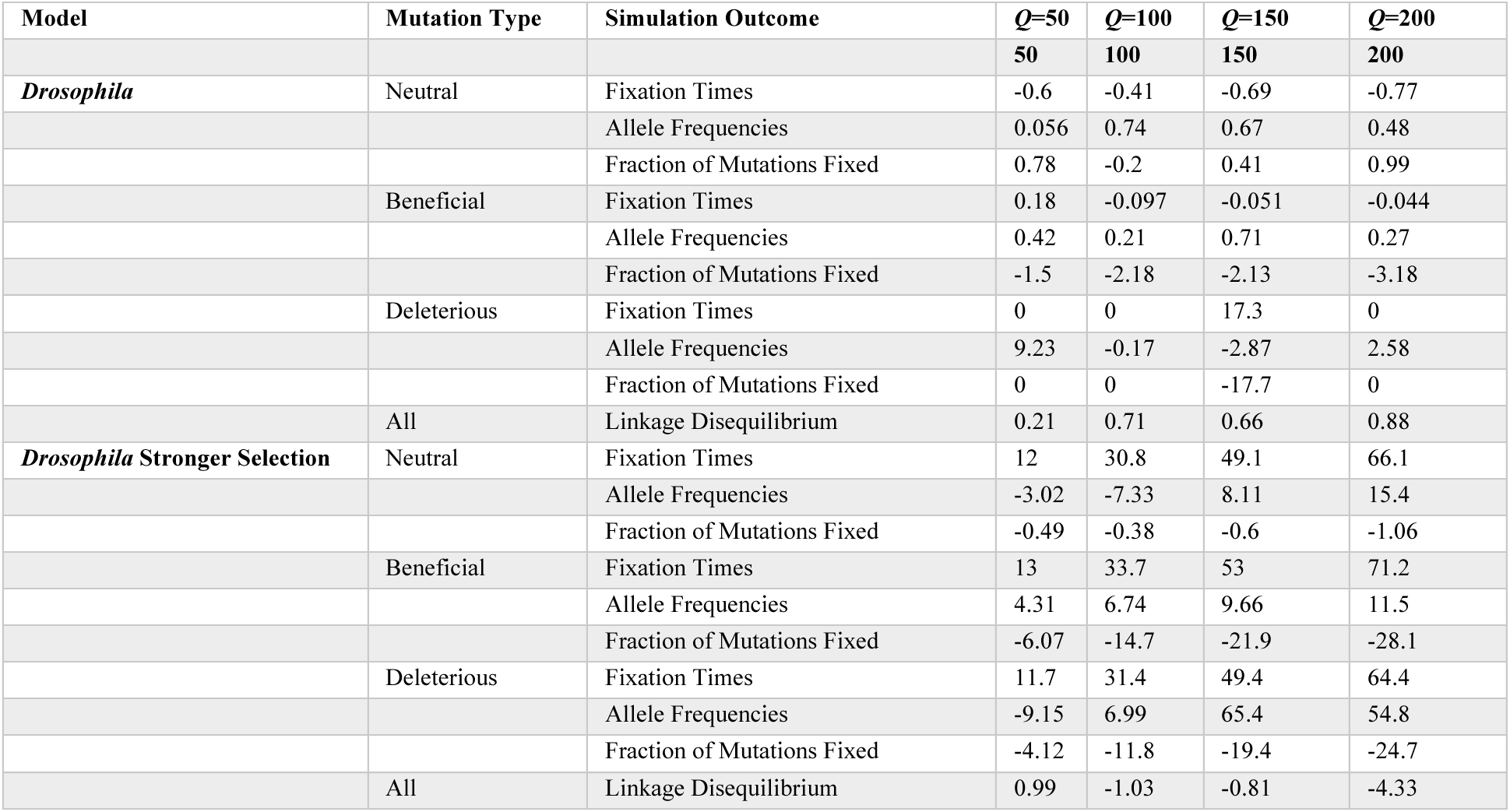
Summary of MPE Results for *Drosophila* Models.

We also tested the effect of scaling in several additional models. To test the effect of starting with a larger unscaled population size, for example, we simulated the no-beneficials model with a population size *N* = 20,000/*Q*. The no-beneficials model was selected since it exhibited the most dramatic effects due to scaling of the models described above. Furthermore, we tested a single sweep model with a population size of 10,000/*Q*, where a single beneficial mutation (*s*=0.05 × *Q*) was introduced in the middle of the genome and then swept to fixation, whereupon the simulation was terminated. If the sweeping mutation was lost, the simulation was reset and run again until fixation occurred. For all models mentioned thus far, we used *Q* values of 2, 5, 10, and 20. We note that the higher values of *Q* here are similar in magnitude to those that have been used to re-scale simulations of human-like populations in previous studies (e.g., *Q*=10 in Schrider 2020 and Hoggart et al. 2007). While scaling to higher values is theoretically possible, we ran into practical limitations with higher values of *Q* causing replicates to fail with `SLiM` emitting an error message indicating that the population had reached infinite fitness. Finally, we tested two *Drosophila melanogaster* constant population size models based on the `stdpopsim` catalog parameters for the species (Adrion et al. 2020) which specifies a population size of 1,720,600/*Q* (Li and Stephan 2006), a mutation rate of 5.49 × 10^-9^ × *Q* per base pair (Schrider et al. 2013), and a recombination rate of 1.98 × 10^-8^ × *Q* per base pair, which is `stdpopsims`’ average recombination rate for all autosomal chromosomes (based on the estimate from Comeron et al. (2012)). For these simulations, we chose a 10 kb genome size and only conducted simulations at *Q* values of 20, 50, 100, 150, 200 due to computational limitations which prevented us from carrying out unscaled simulations of these models without further reducing our genome or population size. Therefore, our baseline for these simulations, against which all other *Q* values are compared, is those simulations with *Q*=20. We note that this range of *Q* values encompasses those used in recent studies conducting forward-in-time simulations of *Drosophila* populations (Kim and Wiehe 2009; Johri et al. 2020; Caldas et al. 2022). For the first model, we used the distribution of fitness effects from Ragsdale et al. (2016). Since this DFE is expressed in units of 2*N*_e_, `stdpopsim` converts the selection coefficients to units of *s* using a population size estimate of 2.8 × 10^6^(Huber et al. 2017). This yielded a lognormal distribution with a μ parameter of 9.679 × 10^-7^ + log(*Q*) and a σ parameter of 3.36 for the selection coefficient of deleterious mutations, and a fixed value of 7.125 × 10^-6^ × *Q* for the selection coefficient of beneficial mutations. Adding log (*Q*) to the μ parameter of a lognormal distribution has the same effect as multiplying all resulting draws by *Q*. The second *Drosophila* model was identical to the first except that the selection coefficient for beneficial mutations was increased to 0.01 × *Q*. All code for performing the simulations and downstream analyses in this study are available at https://github.com/SchriderLab/simscale-snakemake.

### Statistics summarizing simulation outcomes

For each simulation replicate, we recorded the following statistics summarizing the outcome:

- The site frequency spectrum (SFS) for each mutation type, each estimated from the same sample of 100 randomly selected individuals taken at the end of the simulation.
- Linkage disequilibrium (LD), as measured by the *r*’ statistic calculated from the same sample of 100 individuals between all possible pairs of mutations. If the number of segregating mutations is larger than 5000, we first randomly sampled 5000 mutations before calculating *r*’ for all possible pairs of sampled mutations. These *r*’ values were then binned by distance to produce 50 equal distance bins across the genome and the average *r*’ value in each bin was recorded.
- The fraction of mutations that fixed for each mutation type, recorded only for mutations occurring after the burn-in. These values were estimated by taking the fraction of all mutations of a given type that reached fixation and rescaled by dividing this fraction by *Q*. To illustrate the importance of diving by *Q*, consider the case of neutral mutations where the expected fixation probability is 1/2*N*. When the population is scaled by *Q* this estimate becomes *Q*/2*N* and so dividing by *Q* recovers the unscaled fixation probability. However, we also note that our estimate of fraction of mutations fixed does not take into account the number of mutations still segregating at the end of the simulation and is also not necessarily representative of the true probability of fixation since mutations that arose near the end of the simulation may not have had time to fix.
- The distribution of fixation times, recorded separately for each mutation type and only for mutations occurring after the burn-in. These times were all rescaled by multiplying by *Q* and binned to produce around 20-30 bins for each simulation model. The varying number of bins was due to our goal of keeping the width of the fixation time bins constant across all replicates, such that they map correctly to each other. Therefore, a random unscaled replicate was chosen to decide the width of the fixation time bin (in generations) to produce 20 bins, and then propagated to all replicates across all scaling factors. For instance, if the maximum fixation time for the randomly chosen unscaled replicated was 10000 generations, then the bin width would correspond to 500 generations. Since some replicates will have a different maximum fixation time, the number of bins would vary across mutation types and simulation scenarios, but each bin would correspond to the same fixation time range in all replicates and for all scaling factors.

### Assessing the deviation between scaled and unscaled simulations

To assess whether scaled and unscaled simulations could be distinguished from one another for a given simulation model and a given value of *Q*, we trained classifiers to label a given simulation as unscaled or scaled, with the possible input features being the SFS, the binned data for fixation times and LD, or the fraction of fixed mutations as described above. 80% of replicates for a given model and value of *Q* were used to train the classifier, and the remaining 20% of the replicates were set aside for testing. For each dataset, we trained both a logistic regression model and a random forest model. Model parameters were the default values of the Python package scikit-learn, version 1.1.3 (Pedregosa et al. 2011). The input features were grouped into the following four separate feature sets to separately test their ability to distinguish between scaled and unscaled simulations: 1) the SFS across all mutation types (199 features per mutation type) represented as a 1-dimensional vector per replicate, 2) the average *r*’ values at different (binned) inter-mutation distances (50 features, all mutation types), 3) the fraction of mutations fixed (one feature per mutation type), and 4) the histograms of fixation times (20-30 features per mutation type) concatenated along the time axis to produce a 1-dimensional vector per replicate. After training, classification accuracy (fraction of correct predictions on the test data) was recorded. Classifier accuracy in this context provides a reasonable estimate of the extent to which the distributions of the distribution of outcomes across all replicates overlap (with greater classification accuracy implying less overlap).

Comparing classifier accuracies across *Q* values also provides a general high-level picture of the extent to which the distribution of simulation outcomes shifts as *Q* increases. We note that we balanced both our training and test sets, with the latter meaning that 50% accuracy represents the baseline accuracy of an uninformative classifier.

To further characterize the differences between scaled and unscaled simulations, we calculated the mean percent error (MPE) between the scaled and unscaled fixation times, mutation frequencies, LD *r*’ values, and fraction of mutations fixed by first obtaining the mean of each outcome for each replicate, averaging the means across all replicates, and calculating the MPEs as:

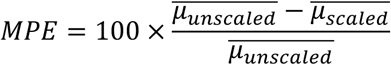

Where μ is the mean of the outcome for each replicate. In cases where the unscaled value of a simulation outcome was zero, MPE is undefined. When plotting such cases we simply omitted the MPE bar, and we note that this causes undefined MPE values to be visually indistinguishable in our bar plots from cases where MPE was very low.

Furthermore, the Kullback-Leibler (KL) divergence between the scaled and unscaled fixation times and site frequency spectra were calculated. KL divergence indicates the amount of statistical distance between probability distributions, and therefore captures information that is complementary to that provided by MPE about the rescaling effect. For instance, a metric may have a smaller MPE at *Q*=10 compared to *Q*=5, but a larger KL divergence, indicating that while the mean of the distribution at *Q*=10 has shifted closer to its unscaled counterpart, the overall shape of the distribution has actually diverged even further (e.g., perhaps due to a large change in the variance, skewness, or kurtosis of the distribution). For two discrete distributions made of up *n* bins with values of *x_n_*, at each bin, the KL divergence between these two distributions is calculated as:

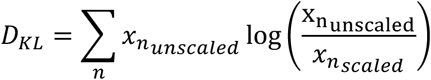

For these calculations, the distributions for each outcome were first averaged for each bin across all replicates for each scaling factor such that:

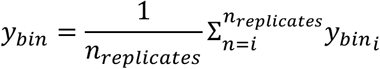

The distributions were then standardized to a probability distribution by dividing every bin value by the sum of values at all bins. KL divergence of the scaled distribution for each outcome was then calculated against the corresponding unscaled distribution. For LD, we opted to use the RMSE instead of KL divergence since we summarized LD by the average *r*’ values within genomic distance bins rather than probability distributions of *r*’ values. This was obtained by averaging the squared error values for each bin in the LD distribution for each scaling factor such that:

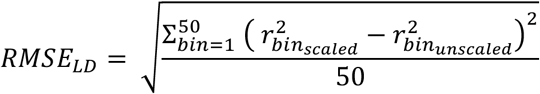

For each of those error measures (MPE, KL divergence, and average RMSE), we generated distributions via bootstrapping. To do this, we sampled the entirety of both the scaled and unscaled replicates 1000 times with replacement, calculating our error measures each time as outlined above, and finally reported the mean and standard deviation.

### Assessing the impact of sample size on estimates of deviation between scaled and unscaled simulations

To investigate the impact of the number of replicates on the estimation of deviation due to scaling, we recalculated our MPEs, KLD, and LD average RMSE values for our simulation outcomes between scaled and unscaled simulations using smaller sample sizes of 100 and 10 for the full model, the no-beneficials model, and the *Drosophila* stronger-selection model. For each of those deviation measures, we sampled the appropriate number of replicates randomly for each scaling factor and recalculated our estimates as described above, and repeated this process 1000 times to acquire a distribution of these estimates under each sampling scheme.

## RESULTS

### Rescaling alters the SFS, LD, and fixation times/probabilities in a simulated population with a gene-dense genome

To characterize the impact of population size rescaling on simulations, we simulated a variety of scenarios and examined how rescaling alters allele frequencies, LD, fraction of mutations fixed, and time to fixation (Methods). We began by examining simulations of a constant-sized population with a genome densely packed with selected mutations (both deleterious and beneficial, with 5% of selected mutations belonging to the latter category); because this is our richest model in terms the amount of natural selection, we refer to it as the “full model” (Methods).

When examining the distribution of mutation fixation times for neutral mutations under the full model (Figure 1A), the differences between the unscaled and scaled distributions were subtle and non-monotonic. For instance, fixation times at *Q*=2 were slightly shifted towards longer sojourns, while those at *Q*=20 were shifted in the opposite direction. The distributions of fixation times for beneficial and deleterious mutations (Figure S1) also showed this non-monotonic trend, with longer sojourns at *Q* of 2, 5 and 10 and shorter sojourns at *Q*=20. The site frequency spectrum for neutral mutations (Figure 1B), again showed a non-monotonic trend across values of *Q*. The fraction of mutations that are singletons, for which differences are most pronounced, decreased with increasing values of *Q* and slightly increased at *Q*=20 compared to *Q*=10. The SFS for deleterious mutations followed a very similar pattern to neutral mutations, while for beneficial mutations the fraction of singletons increased more dramatically at *Q*=20 (Figure S1). The amount of LD across the genome markedly changed with *Q* (Figure 1C) as it appears that rescaling increased LD between mutations regardless of the distance between them. Furthermore, the fractions of mutations fixed for all mutation types also changed with *Q* (Figure 1D), with the largest shift occurring for beneficial mutations; we note that this comparison was done after rescaling all fraction of mutations fixed estimates themselves to account for the impact of rescaling on the fraction of neutral mutations that fixed (Methods). For example, for beneficial mutations, the fraction of mutations fixed was around 0.20% for unscaled simulations and gradually decreased to around 0.05% for *Q*=20. On the other hand, the fraction of neutral and deleterious mutations fixed remain unchanged at ∼ 0.005% and 0.003%, respectively, for all values of *Q*.

**Figure 1.**
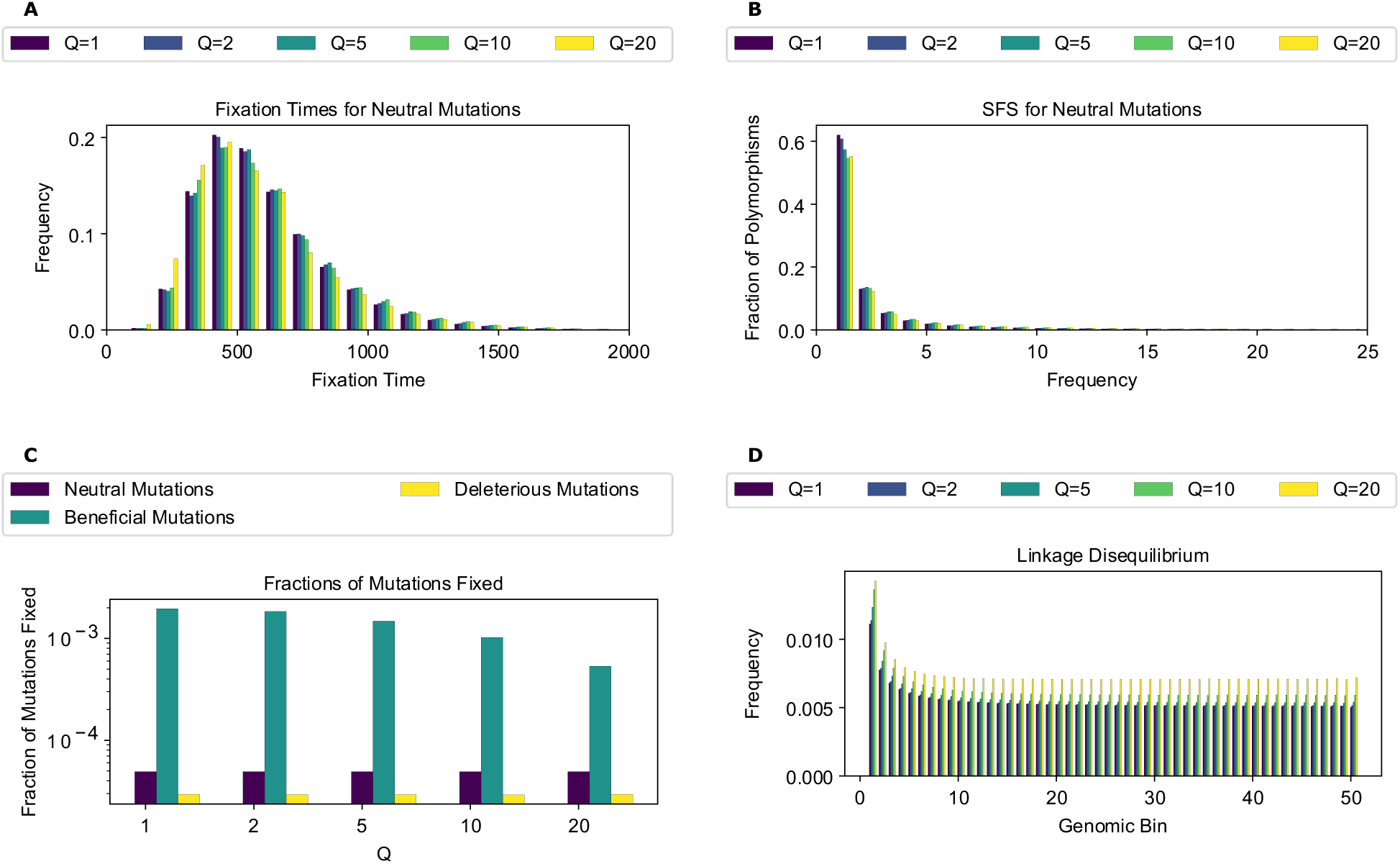
Select simulation results for the full model across scaling factors. **(A)** Fixation times for neutral mutations. **(B)** The site frequency spectra for neutral mutations. **(C)** Fractions of mutations fixed for all mutation types. **(D)** Linkage disequilibrium across 50 genomic bins for all mutation types. As higher values of *Q* are used, all statistics showed larger deviations from values observed in unscaled simulations. The fraction of beneficial mutations that reached fixation was impacted markedly more than for other mutation types.

Next, we trained classifiers to determine whether scaled and unscaled simulations were distinguishable from one another based on these simulation outcomes. To assess this, we trained a set of binary classifiers, each using a different set of input features (fixation times, the SFS, fractions of mutations fixed, and LD) to discriminate between unscaled simulations at a given value of *Q*. As *Q* increases, classification accuracy increases for the SFS- and LD-based classifiers, starting at 72% for the SFS at *Q*=2 and increasing to 100% at *Q*=10 for the logistic regression classifier. For LD, the accuracy of logistic regression increased from 60% at *Q*=2 to 99.75% at *Q*=10. For fixation times and fraction of mutations fixed, accuracy was already at 100% when *Q*=2 (Figure 2A), consistent with a larger deviation between scaled and unscaled simulation outcomes at higher values of *Q*. Similar results were obtained by the random forest classifiers (Figure 2B). Most importantly, we found that all feature sets could be used to distinguish between scaled and unscaled simulation replicates, even at relatively modest values of *Q*. For example, the minimum accuracy observed at *Q*=5 was ∼90% for both classifiers (both examining LD).

**Figure 2.**
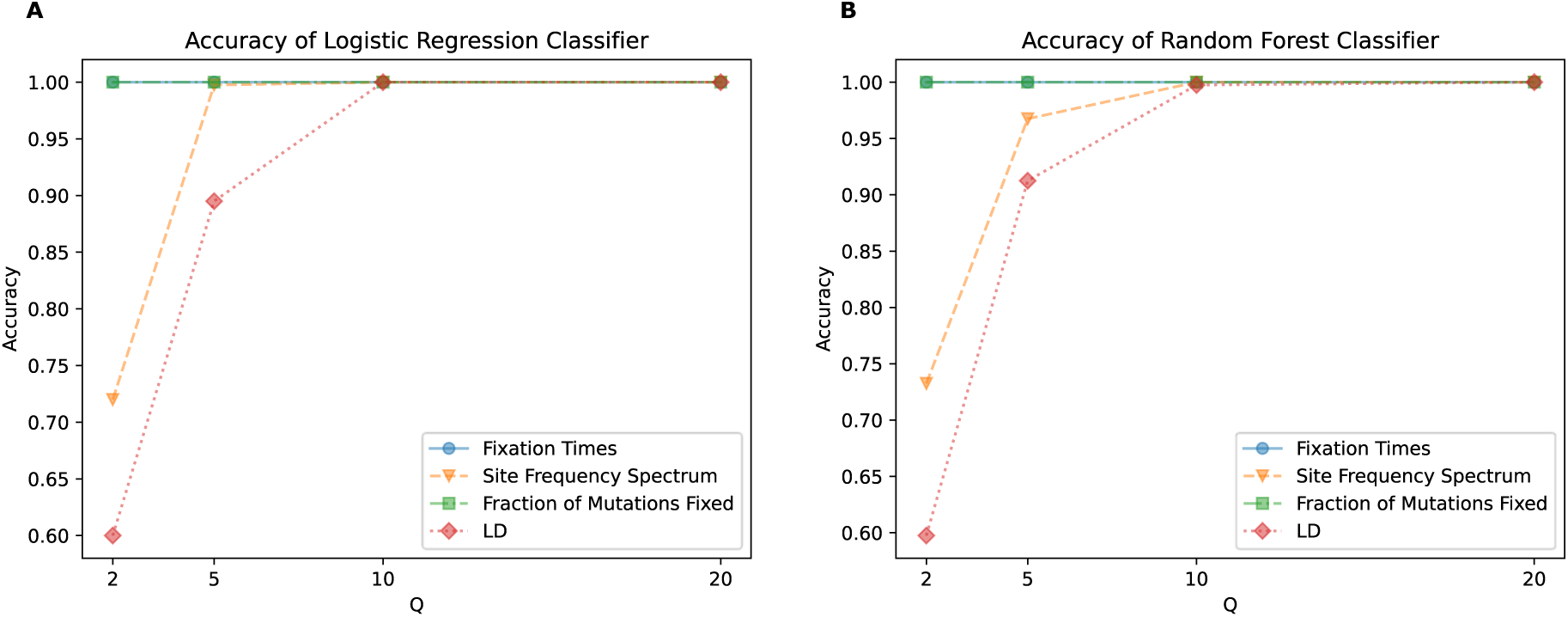
Classifier accuracy for models trained to distinguish between scaled and unscaled replicates of the full model. **(A)** Accuracy of the logistic regression classifier trained to classify a given simulation replicate as scaled or unscaled. **(B)** Accuracy of the random forest classifier trained to classify a given simulation replicate as scaled or unscaled. A separate classifier was trained for each *Q* value and each set of features (those summarizing the SFS, the fraction of mutations that fixed, LD, and the distribution of fixation times). Accuracy represents fraction of simulation replicates in the test dataset correctly identified as scaled or unscaled.

The high accuracy obtained by classifiers discriminating between unscaled and rescaled simulations demonstrates that the rescaling effects summarized in (Figure 1) are significant and systematic.

Having established that rescaling significantly influences simulation outcomes, we sought to quantify the magnitude of this effect. We first examined the mean percent deviation between scaled and unscaled simulations (hereafter referred to as mean percent error, or MPE) for each feature examined above (Figure 3). MPE for fixation times of all mutations and allele frequencies of neutral mutations showed a non-monotonic rescaling effect across values of *Q*. For fixation times, MPE for all mutation types increased between *Q* values of 2 and 5, but decreased with *Q*=10, and then flipped sign at *Q*=20 where it exhibited the largest magnitude across scaling factors. The decrease in error between *Q*=5 and *Q*=10 occurs due to an initial increase in the mean fixation time as *Q* increases. For example, in the case of neutral mutations the mean fixation time increased from 622 generations at *Q*=1 to 628 and 636 generations at *Q* of 2 and 5, respectively. This is followed by a decrease in mean fixation time to 634 and 601 generations at *Q* of 10 and 20, respectively. This caused the mean fixation time to be closer to that of the unscaled simulation at *Q*=10 compared to *Q*=5, before it decreased below the unscaled simulation mean at *Q*=20. When comparing the different mutation types within each *Q*, fixation time MPEs appeared to be the lowest for beneficial mutations across all values of *Q* while neutral and deleterious mutations both exhibited slightly larger MPEs that are similar to one another. For allele frequencies, we observed a considerably higher MPE, which increased monotonically in magnitude from *Q*=2 to *Q*=20 for all mutation types. Here, neutral mutations had the highest MPE across *Q* values, reaching ∼44% for *Q*=20, while beneficial mutations had the lowest error at −6.77% for *Q*=20 and were the only type of mutation to experience a sign change in MPE (occurring at *Q*=20). MPE was also large for LD and for the fraction of beneficial mutations fixed, and again increased in magnitude with *Q*. However, deleterious and neutral mutations exhibited miniscule MPEs for their fraction of mutations fixed.

**Figure 3.**
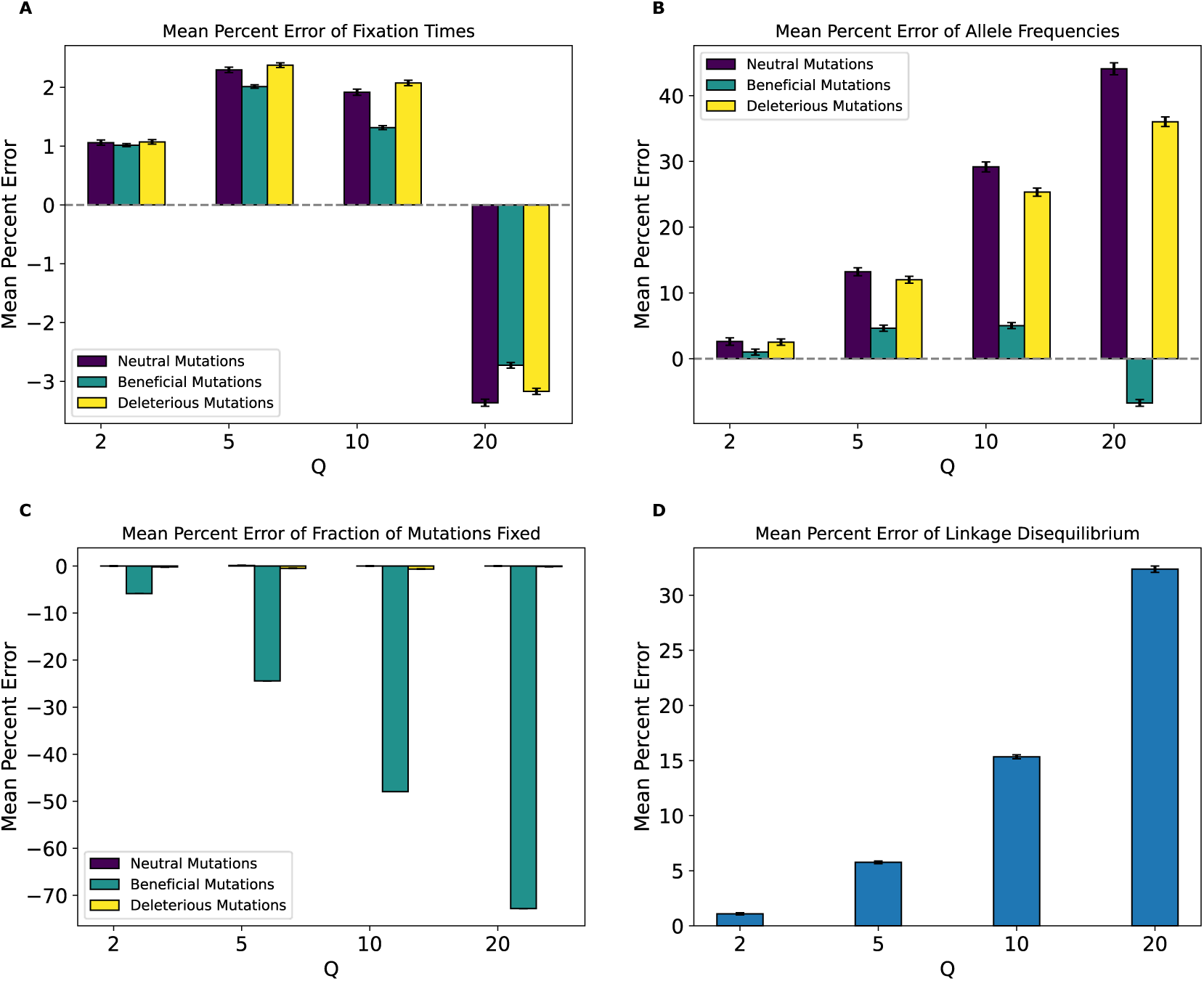
Mean percent error of various simulation outcomes for the full model by values of *Q*. **(A)** Mean percent error for average mutation fixation times. **(B)** Mean percent error for average allele frequencies. **(C)** Mean percent error for average fractions of mutations fixed. **(D)** Mean percent error for average linkage disequilibrium as measured by the values of *r*^2^. For **(A)**, **(B)**, and **(D)** the mean value was calculated for each replicate, followed by averaging the means across all replicates in each value of *Q*.

For **(C)**, we averaged across all replicates only, since each replicate has only one value for the fraction of mutations fixed for each mutation type. The mean percent error was calculated between the resulting value from each *Q* and the corresponding value at *Q*=1. This process was performed for 1000 bootstrap samples and the values and error bars shown in all panels represent the mean and standard deviation across the bootstrapped distributions.

As an alternative way of quantifying the degree of deviation between scaled and unscaled simulations, we examined KL divergence for fixation times and SFS distributions and RMSE for LD (Figure S2). For the distribution of fixation times, all mutation types showed similar KL divergence within each scaling factor, with KL divergence increasing steadily for each mutation type as *Q* increases. This trend changed at *Q*=20, where beneficial mutations displayed substantially higher KL divergence than other mutation types. Similarly, when examining the SFS, KL divergence for each mutation type increased with *Q*, although in this case, and while beneficial mutations again had the highest KL divergence at *Q*=20, neutral mutations had markedly higher KL divergence at *Q*=10. For LD decay, the root mean squared error (RMSE) between scaled and unscaled simulations showed a similar trend as MPE, increasing with *Q* from around 0.00007 at *Q*=2 to around 0.002 at *Q*=20.

### Rescaling effects persist in simulations with no beneficial mutations

Our full model serves as a useful stress test of the robustness of forward simulations to rescaling but contains a larger number of beneficial mutations than will be realistic in most cases (5% of non-neutral mutations). To see how rescaled simulations would behave without such a pervasive impact of positive selection, we altered this model by removing all beneficial mutations (the “no-beneficials” model). Under this model, we again observed a significant scaling effect, as our classifiers were able to discriminate between scaled and unscaled simulations with higher accuracy than those obtained in the full model except for fixation times at *Q*=2, where classifiers were less accurate for this model compared to the full model. (Figure S4).

We next examined the magnitude of the rescaling effect for the no-beneficials model as measured by MPE. We observed much larger MPEs in the no-beneficials model (Figure 4) than for the full model for fixation times, LD, and fraction of fixed mutations. However, we observed smaller MPEs for allele frequencies than under the full model (MPE is approximately 5% at *Q*=10 compared to ∼29% for neutral mutations in the full model and increases to over 23% at *Q*=20 compared to 44% for neutral mutations in the full model), with both mutation types having similar errors across values of *Q*. Of note here is that unlike our full model, the magnitude of error increased monotonically and in the same direction for all measured statistics, with fixation times and allele frequencies exhibiting negative MPE. This indicates that rescaling in this model lowered average fixation times and allele frequencies, rather than increasing them as was the case for most values of *Q* under the full model. LD and fraction of mutations fixed, on the other hand, exhibited positive MPE, with the MPE for LD increasing with *Q* in a similar manner to the full model, with slightly smaller values compared to the full model at all values of *Q* except 20, where the error is around 37% compared to the full model’s 32%. The MPE for fraction of mutations fixed was quite high, especially for deleterious mutations, with values peaking at ∼141% for deleterious mutations and ∼64% for neutral mutations at *Q*=20. Interestingly, under the full model, the corresponding MPEs for these mutation types were negligible, as that model instead showed large MPEs for beneficial mutations, suggesting a substantial change in the dynamics of rescaling effects depending on the presence or absence of large numbers of beneficial mutations.

**Figure 4.**
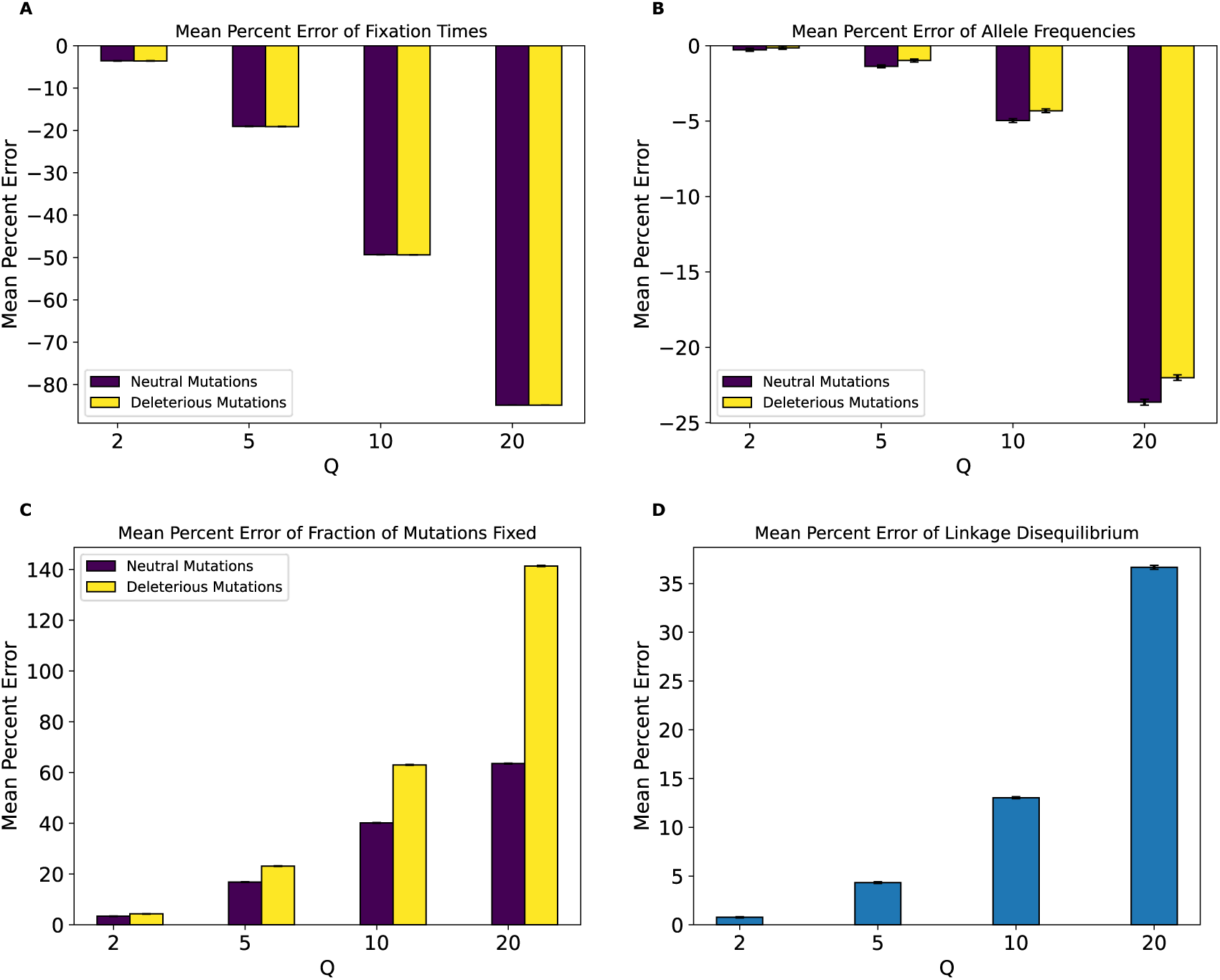
Mean percent error of various statistics for the no-beneficials model by values of *Q*. **(A)** Mean percent error for average mutation fixation times. **(B)** Mean percent error for average allele frequencies. **(C)** Mean percent error for average fractions of mutations fixed. **(D)** Mean percent error for average linkage disequilibrium as measured by the values of *r*^2^. Values and error bars represent the mean and standard deviation of 1000 bootstrapped estimates.

KL divergence for the fixation time distribution in the no-beneficials simulations (Figure S4) again increased monotonically with *Q*, with similar values for both mutation types, but showed a much greater magnitude than observed for the full model across *Q* values. For the SFS, KL divergence was lower than that of the full model for lower values of *Q*, but suddenly increased to higher values than observed under the full model at *Q*=20. Consistent with the full model, however, neutral mutations showed larger KL divergence than deleterious ones. For LD, KL divergence increased monotonically with *Q* and exhibited higher values at all values of *Q* compared to the full model. Overall, the comparison between the full model and no-beneficials model suggests that the presence of large amounts of positive selection may, at least in some cases, actually diminish the impact of rescaling on simulation outcomes.

Given the large scaling effects present in this model, we sought to test the impact of increasing the population size on these effects. To this end, we carried out simulations of this same model containing only deleterious and neutral mutations but with double the population size (N=20000). Results indicate that MPE for fixation times (Figure 5A) and fractions of mutations fixed (Figure 5C) were nearly identical to their counterparts in the no-beneficials model (compare to Figure 4). MPE for LD and allele frequencies were slightly diminished (Figure 5D and B), on the other hand, compared to their counterparts in the no-beneficials model. This change was not uniform across all values of *Q*. For instance, increasing the unscaled population size decreased the MPE for allele frequencies by around ∼3% at *Q*=20 for both neutral and deleterious mutations, but only by ∼1% at *Q*=10. For LD, increasing the population size reduced the MPE by ∼10% at *Q*=20 and by ∼5% for *Q*=10.

**Figure 5.**
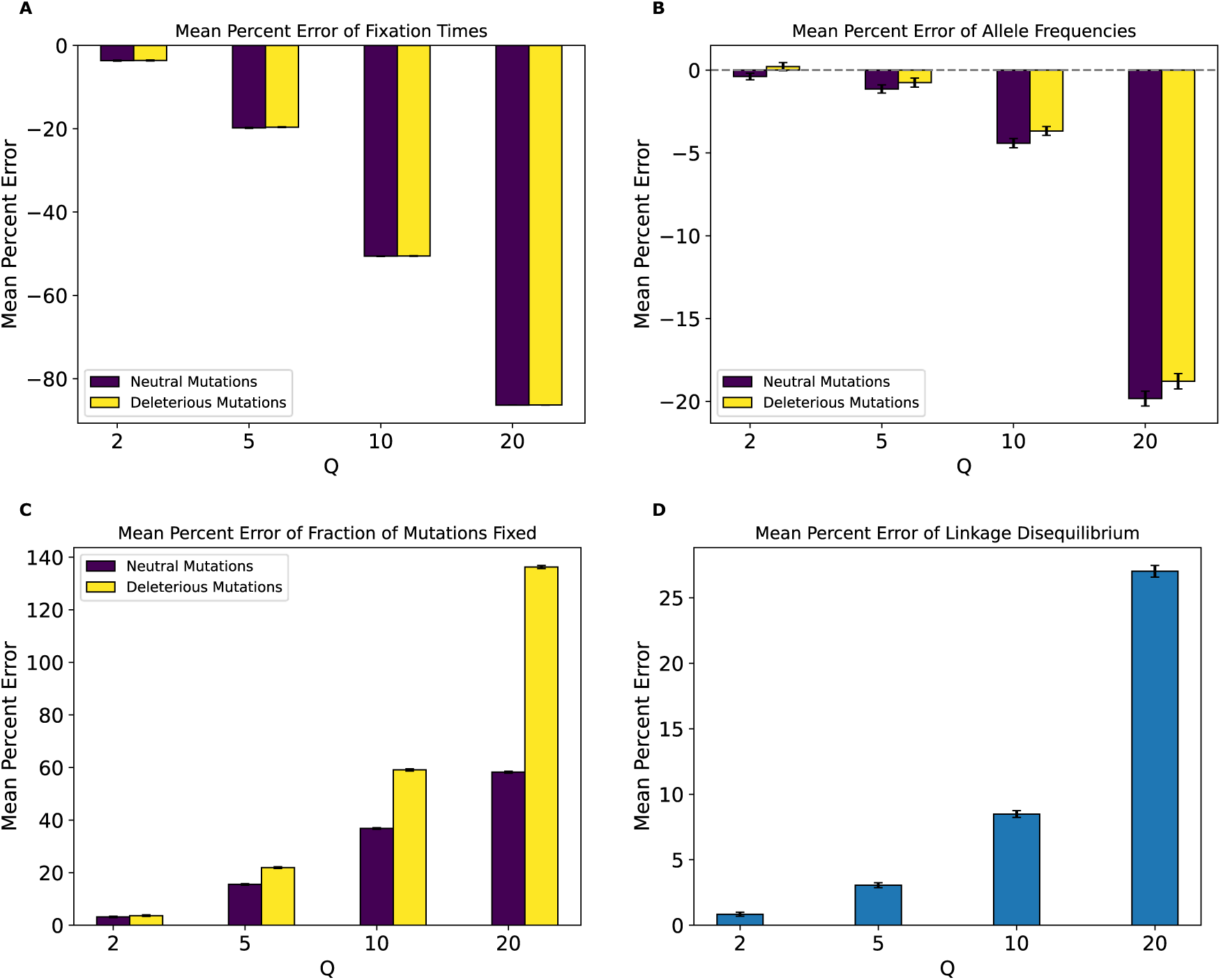
Mean percent error of various statistics for the no-beneficials larger population model by values of *Q*. **(A)** Mean percent error for average mutation fixation times. **(B)** Mean percent error for average allele frequencies. **(C)** Mean percent error for average fractions of mutations fixed. **(D)** Mean percent error for average linkage disequilibrium as measured by the values of *r*^2^. Values and error bars represent the mean and standard deviation of 1000 bootstrapped estimates.

### Rescaling effects in a sparse-genome model with no beneficial mutations

Next, we examined a model with a much smaller fraction of deleterious mutations (5% of all mutations), as might be expected in a large genome where functional elements are sparse. As in the previous section, we did not include beneficial mutations. Under this model, called the no-beneficials sparse model, our classifiers were not as effective at discriminating between unscaled and rescaled simulations at lower values of *Q* (Figure S8), especially for fixation times and fractions of mutations fixed where the random forest model accuracy is slightly higher than 50% for both at *Q*=2 and is 74% and 72% respectively at *Q*=5. However, we note that at *Q*=20 our classifiers were still quite accurate, reaching 100% on all feature sets for both classifiers. Thus, while simulation outcomes were not as impacted at lower *Q* values compared to the full model or the no-beneficials model, they still appeared to be significantly affected by rescaling under our no-beneficials sparse model, particularly at higher *Q* values.

Concordant with our decreased classifier accuracy, the MPE of fixation times for this model was lower across all statistics than that of the no-beneficials model (Figure 6). Again, the MPEs for allele frequencies, fixation times, fractions of mutations fixed, and LD increased monotonically with *Q*. However, one notable difference is that the direction of the bias caused by rescaling, as indicated by the sign of the MPE, was reversed for allele frequencies compared to the no-beneficials model—here we typically observed a slight increase in the average frequencies of neutral and deleterious mutations, as was observed under the full model rather than the no-beneficials model. However, we again stress that the magnitude of this bias was much smaller under this model than either the full or the no-beneficials model (MPE ∼1% for allele frequencies at *Q*=20 with considerable variance for deleterious mutations). The MPE was still appreciable for the other simulation outcomes at *Q*=20, however (∼-8% for fixation times, ∼18% for LD, and ∼35% for fractions of mutations fixed).

**Figure 6.**
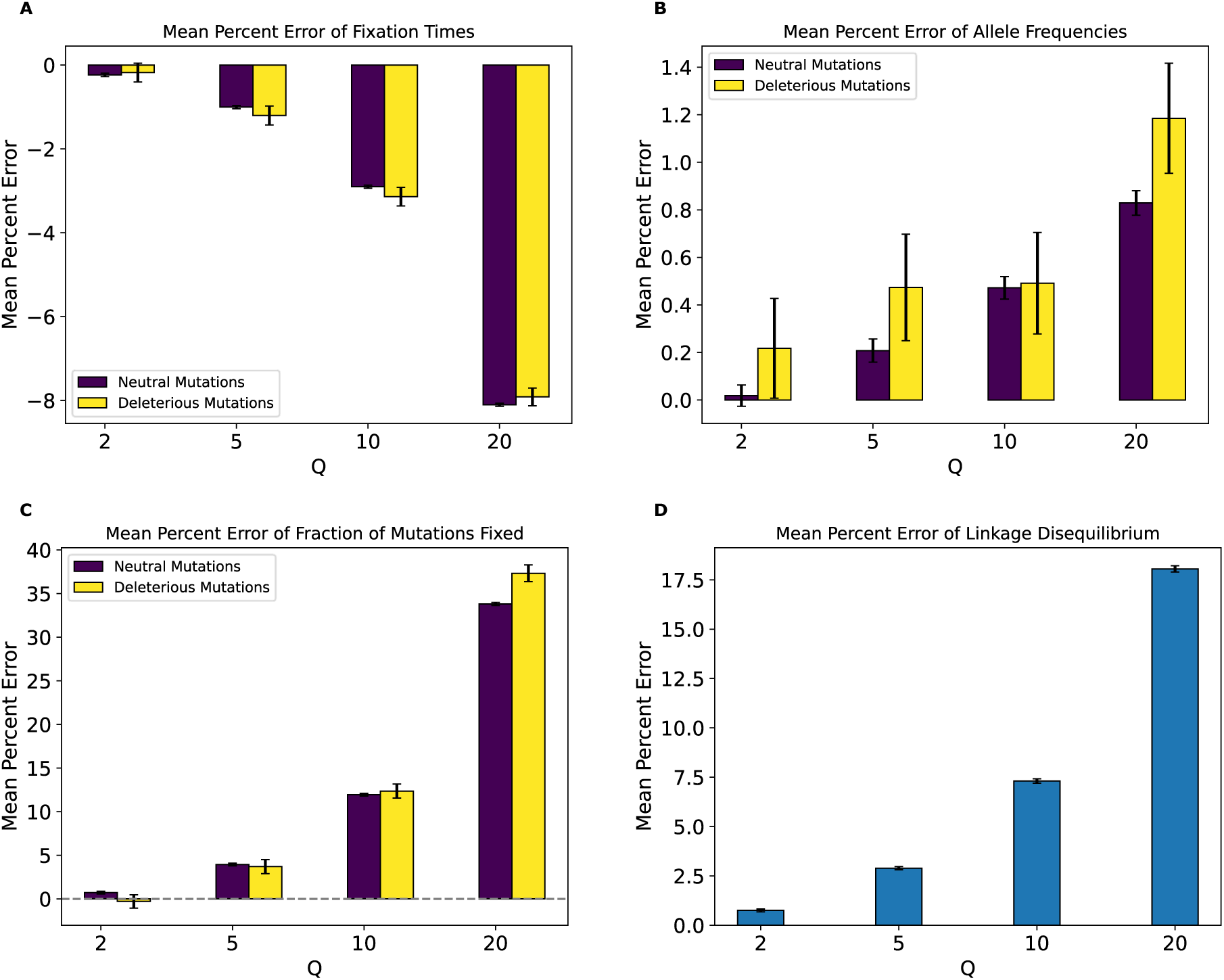
Mean percent error of various statistics for the no-beneficials sparse model by values of *Q*. **(A)** Mean percent error for average mutation fixation times. **(B)** Mean percent error for average allele frequencies. **(C)** Mean percent error for average fractions of mutations fixed **(D)** Mean percent error for average linkage disequilibrium as measured by the values of *r*^2^. Values and error bars represent the mean and standard deviation of 1000 bootstrapped estimates.

KL divergence for fixation times and the SFS (Figure S8) revealed that for simulations under the no-beneficials sparse model, the distributions of outcomes for deleterious mutations were generally more impacted by rescaling than are neutral mutations, except for fixation times at *Q*=20 where the divergence was similar for both mutation types. When compared with the no-beneficials model and, consistent with the smaller MPE values observed in this model, KL divergence for fixation times and the SFS were also smaller compared to the no-beneficials model. For instance, KL divergence for fixation times for both mutation types at *Q* =20 was ∼17 in the no-beneficials model and was ∼0.04 in the no-beneficials sparse model. SFS KL divergence for neutral and deleterious mutations was around 0.0003 and 0.0007 respectively at *Q*=20, compared to 0.048 and 0.036 for the corresponding values in the no-beneficials model. Also consistent with the MPE trends for this model is the observation that KL divergence for fixation times is similar for both mutation types and is higher for deleterious mutations in the case of the SFS. RMSE for LD is also lower compared to the no-beneficials model, measured at 0.001 at *Q* of 20 for this model compared to the corresponding no-beneficials value of 0.002.

### Strictly neutral model

Next, to investigate the degree to which natural selection (whether linked or direct) is responsible for the rescaling effects described above, we performed simulations identical to the previous models but with neutral mutations only (the strictly neutral model). In this case, classification of simulations as scaled or unscaled became more challenging for both models (Figure S10) and fixation times and fractions of mutations fixed were uninformative features for all values of *Q*. However, some impacts of scaling on the SFS and LD appeared to persist, as the random forest model achieved an accuracy of approximately 89% and 100% on those features, respectively, at *Q*=20. This indicates that rescaling can still impact allele frequencies and LD even in the absence of selection, as previously shown by Adrion et al. (2020). These results were corroborated by the MPE values for these statistics (Figure 7) which showed very small values and high variance (relative to the mean) for fixation times and fractions of mutations fixed but larger values with smaller variance (relative to the mean) for allele frequencies and LD. For LD, the MPE at *Q*=20 was around 16% under the strictly neutral model, which is comparable to that of the full model at *Q*=10, while the allele frequency MPE at *Q*=20 was around 1.22% under the strictly neutral model which is smaller than the MPE for neutral allele frequencies under the full model (the MPE for the stricly neutral model at *Q*=20 was comparable to that of the full model at *Q*=2). Fixation times and SFS KL divergence and LD RMSE values for this model (Figure S10) were also smaller than for the full model, but the LD RMSE was the closest with a value of ∼0.001 at *Q*=20 under the strictly neutral model, compared to ∼0.002 for the corresponding value in the full model.

**Figure 7.**
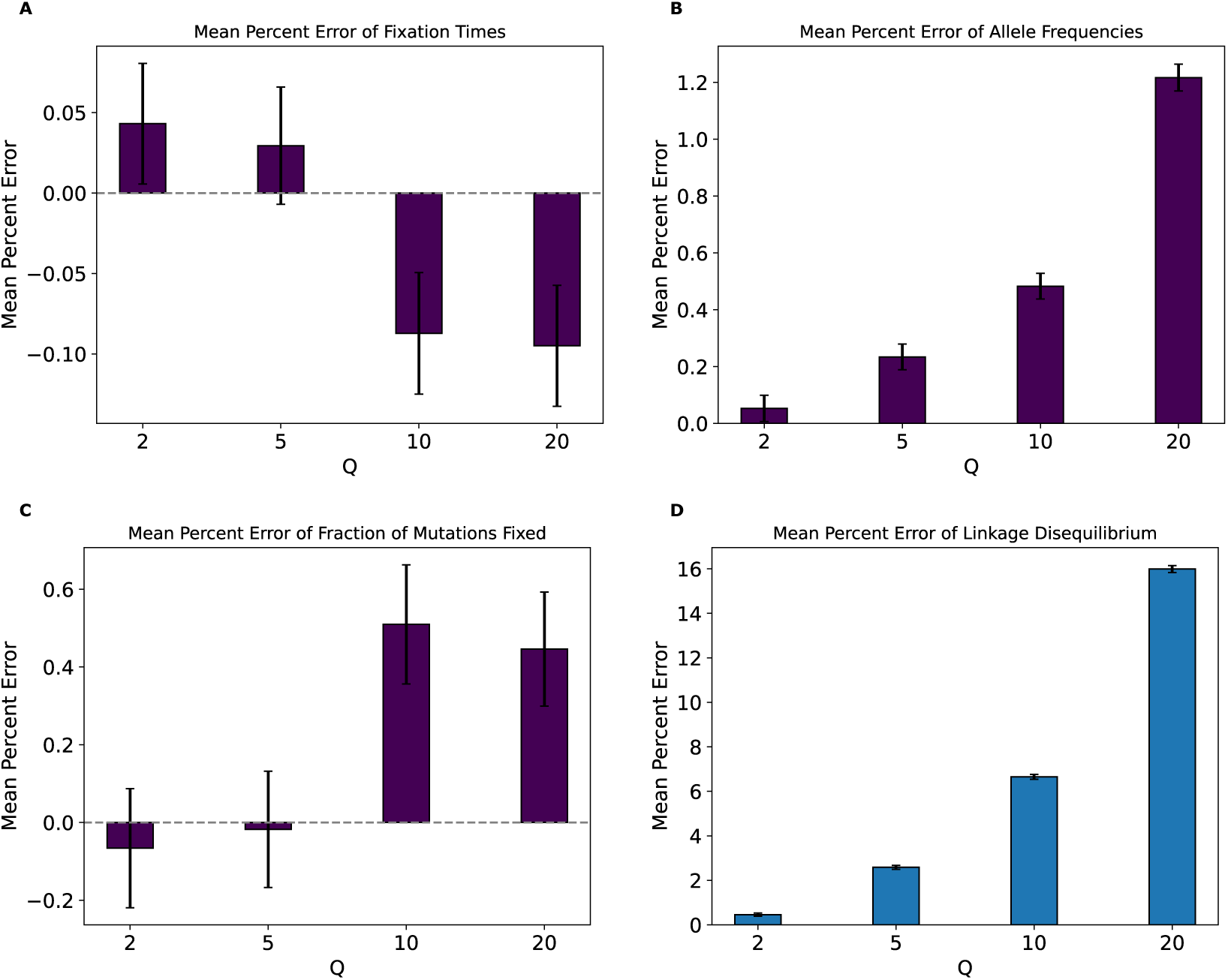
Mean percent error of various statistics for the strictly neutral model by values of *Q*. **(A)** Mean percent error for average mutation fixation times. **(B)** Mean percent error for average allele frequencies. **(C)** Mean percent error for average fractions of mutations fixed. **(D)** Mean percent error for average linkage disequilibrium as measured by the values of *r*^2^. Values and error bars represent mean and standard deviation of 1000 bootstrapped estimates.

### The rescaling effect is visible in both expanding and contracting populations

Researchers are often interested in observing evolutionary dynamics in populations experiencing demographic changes. We therefore next sought to examine the effect of population size changes on the rescaling effect. First, we simulated a variant of our full model where the population experienced a recent two-fold expansion, the population expansion model. (Methods). Our classifiers for distinguishing between rescaled and unscaled simulations performed similarly to those for the full model (Figure S12), although accuracy was generally slightly lower under the population expansion model for LD features across values of *Q* below 20. MPEs for fixation times (Figure 8A) for all mutations were higher across all values of *Q* except for 20, where the MPE became smaller in magnitude compared to the full model. For instance, the MPE for fixation times of neutral mutations at *Q* values of 2, 5, and 10 were around 1.3%, 3%, and 3.6% respectively for this model compared to around 1.1%, 2.3%, and 2% respectively for the full model. At *Q*=20 however, the error here decreased to 1.5% compared to the full model’s −3.4% (note the sign change, which did not occur under the population expansion model). The MPEs for all other statistics (Figure 8B-D) were very similar to those under the full model but were smaller for LD across all *Q* values. KL divergence values (Figure S12) for fixation times were lower in this model compared to the full model, with beneficial mutations having the lowest KL divergence at *Q*=10 and *Q*=20 in contrast to the full model (where beneficial mutations exhibited the largest KL divergence at these *Q* values). For the SFS, KL divergence was very similar to the full model, while the LD RMSE values were lower across all values of *Q*. Overall, simulations of a population experiencing a recent expansion seem to produce a rescaling effect that is fairly similar to that of a simulated constant-size population, with the biggest observed differences being in the magnitude of KL divergence for LD for all *Q*s and the magnitude and direction of KL divergence for fixation times at *Q*=20.

**Figure 8.**
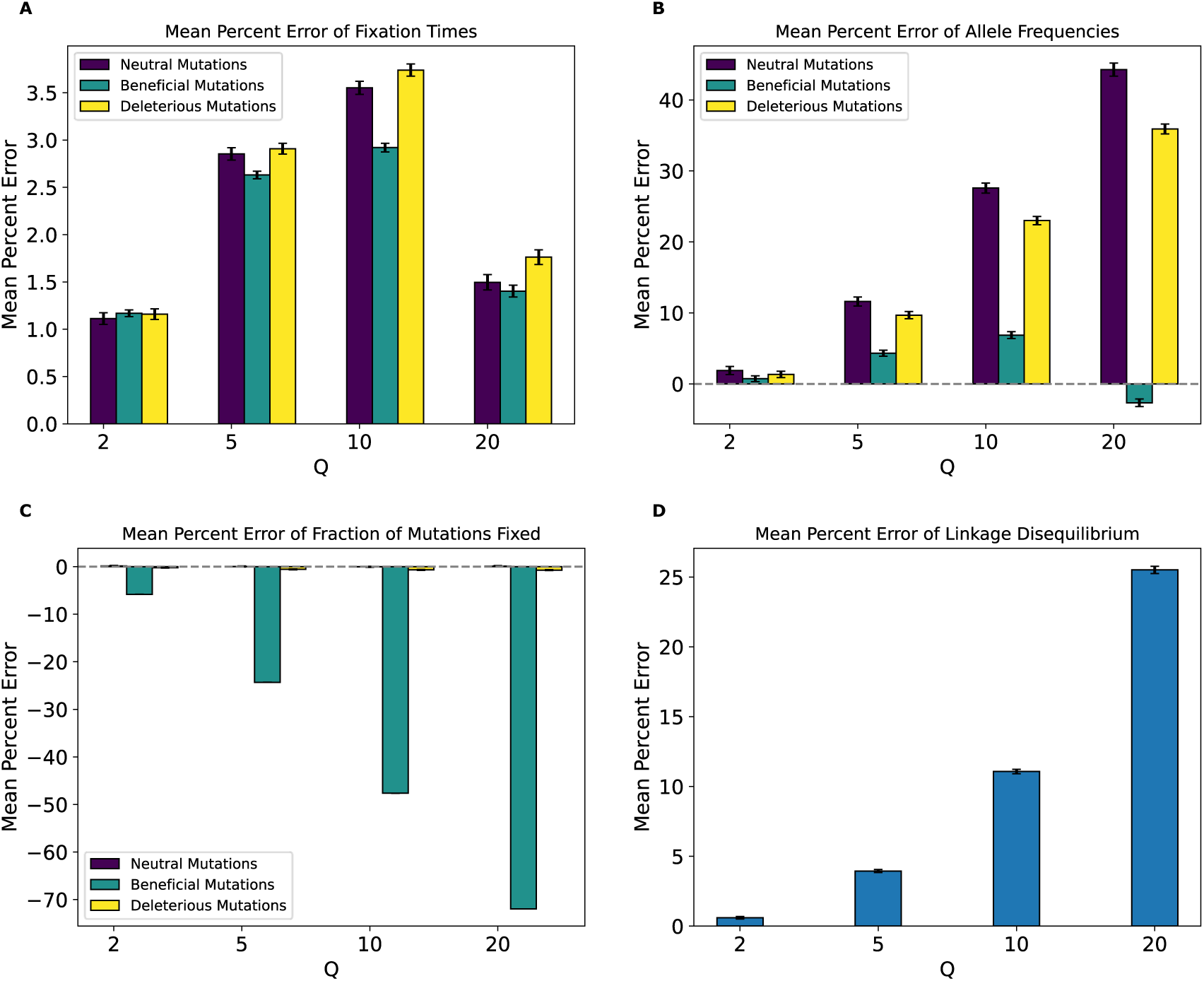
Mean percent error of various statistics for the population expansion model by values of *Q*. **(A)** Mean percent error for average mutation fixation times. **(B)** Mean percent error for average allele frequencies. **(C)** Mean percent error for average fractions of mutations fixed. **(D)** Mean percent error for average linkage disequilibrium as measured by the values of *r*^2^. Values and error bars represent mean and standard deviation of 1000 bootstrapped estimates.

Our next exploration of the effect of population size change was the simulation of a two-fold *decrease* in population size, the population contraction model. Again, our classifiers for this model performed similarly to those for the constant-size model, although the classifiers examining LD were slightly more accurate (Figure S14). Comparing MPE between the population contraction and full models, we observed fairly similar results for allele frequencies, fractions of mutations fixed, and LD (Figure 9). However, for fixation times, we observed smaller MPE values for *Q*=2 through *Q*=10, and a sign change from positive to negative MPE occurred at *Q*=10 whereas this was not observed until *Q*=20 under the full model. More notably, the magnitude of MPE for fixation times suddenly increased at *Q*=20 to values much larger than those observed under the full model (∼8–10%, depending on the mutation type, for the population contraction model vs. ∼3% for the full model). Concordantly, fixation times showed similar KL divergence under the population contraction model (Figure S14) compared to the full model except for *Q*=20 where KL divergence increased markedly compared to the full model, with values of 0.058, 0.051, and 0.065 for neutral, beneficial, and deleterious mutations at *Q*=20 compared to the full model’s values of 0.018, 0.025, and 0.017 respectively. Beneficial mutations were also no longer showing the highest KL divergence for fixation time across any *Q*. The KL divergence for the SFS was very similar to the full model, with slightly higher KL divergence for beneficial. For LD, RMSE values were all slightly higher than the full model, with a value of 0.0027 at *Q*=20 compared to the full model’s 0.002.

**Figure 9.**
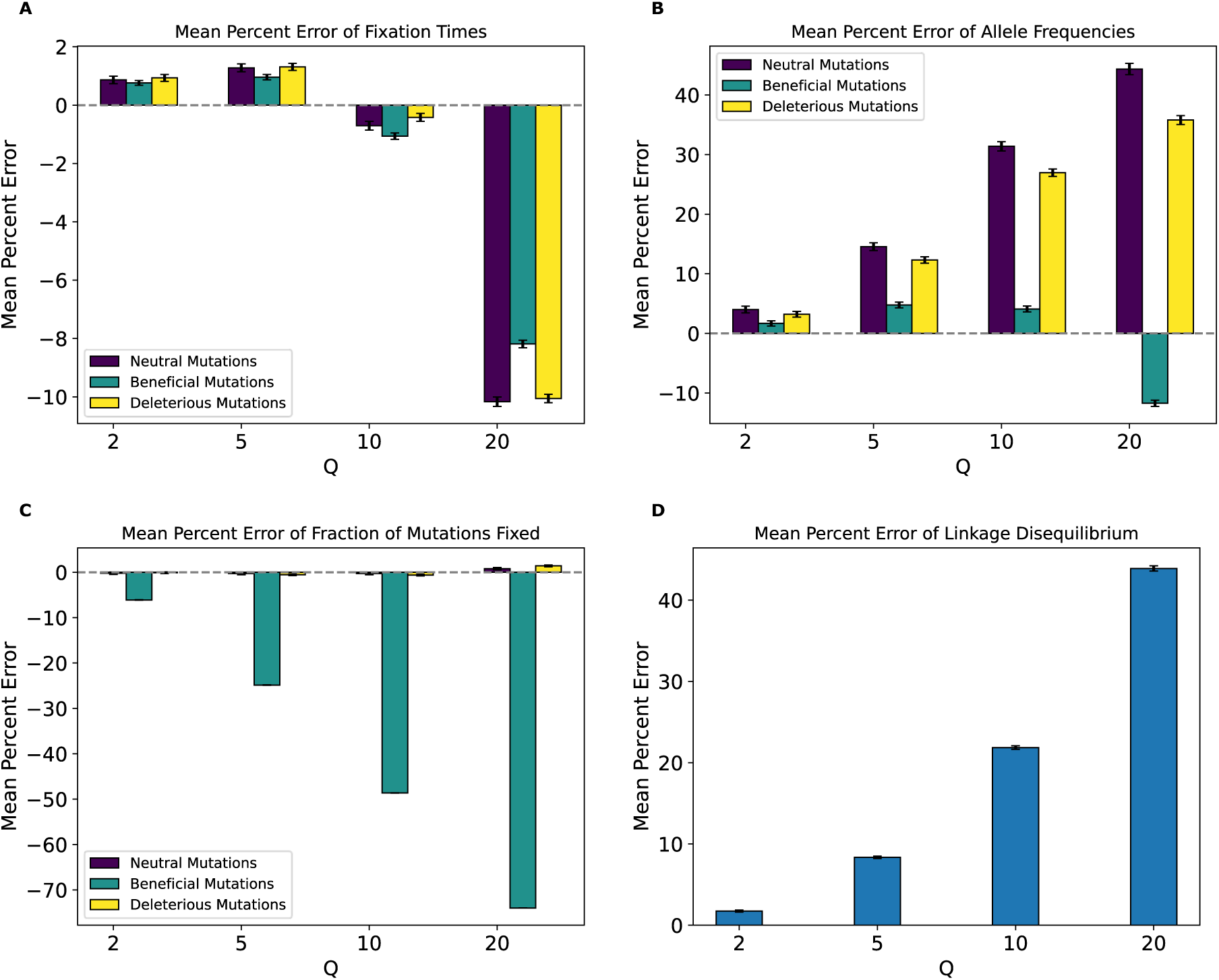
Mean percent error of various statistics for the population contraction model by values of *Q*. **(A)** Mean percent error for average mutation fixation times. **(B)** Mean percent error for average allele frequencies. **(C)** Mean percent error for average fractions of mutations fixed. **(D)** Mean percent error for average linkage disequilibrium as measured by the values of *r*^2^. Values and error bars represent mean and standard deviation of 1000 bootstrapped estimates.

### Scaling effects are greatly diminished in a larger simulated population modeled after *Drosophila* populations

Next, to investigate the impact of rescaling in a more empirically grounded model with a larger population, we performed simulations of a *Drosophila melanogaster* population (Methods). The *Q* values used for the *Drosophila* and the *Drosophila* stronger-selection models were 20, 50, 100, 150, and 200, with simulation outcomes at every *Q* compared to those at *Q*=20, which served here as the baseline instead of *Q*=1 in the previous models. When applying the classification procedure to the *Drosophila* model, we observed lower performance across all values of *Q* than seen for any of the other simulation models examined above (Figure S17). While accuracy hovered around 50% (i.e., random guessing) for all features across almost all *Q* values, the random forest classifier using the fractions of mutations fixed achieved 72% accuracy at *Q*=20, suggesting that the fraction of mutations fixed was the simulation outcome most affected by rescaling in this model. The SFS also appeared to be affected more than other outcomes with a random forest accuracy of around 66% at *Q*=20. Note that the lower accuracy here may be in part due to the smaller chromosomes we simulated for this model because of its greater computational burden (Methods). Thus, there were fewer mutations within each replicate, providing less information to the classifiers. However, this would not affect KL divergence and MPE as much since these measures aggregate information across replicates (although those metrics could still be inflated by noise in sparse datasets; see below).

When examining MPE (Figure S16) and KL divergence (Figure S17) we observed substantial bias in allele frequencies, fixation times and fractions of mutations fixed for deleterious mutations, but observed comparatively subtle effects for neutral and beneficial mutations. For instance, the average fixation time MPE was ∼17% at *Q*=150 for deleterious mutations, but only around −0.7% and −0.05% for neutral and beneficial mutations, respectively. The MPE for fraction of mutations fixed was also high for deleterious mutations at *Q*=150 (∼-18%) and low for beneficial and neutral mutations (∼-2.13% and ∼0.41% respectively). The MPE trend for both the fractions of fixed mutations and average allele frequencies were not monotonic for any of the mutation types. However, it is important to note here that the probability of the fixation of deleterious mutations was extremely small, ∼3.3 × 10^(C’^for *Q*=20 and no such fixations were observed for *Q* values of 50, 100, and 200 (yielding an estimated fractions of fixed mutations of zero). This also means that the distribution of fixation times for deleterious mutations was based on very sparse information and in some cases could not be estimated at all. This is confirmed by examining the MPE standard deviation from the bootstrapping procedure, which is very large for deleterious mutations for both fixation times and the fraction of mutations fixed. This was not a concern for neutral and beneficial mutations, however, and we observed that KL divergence for fixation times of these mutations (Figure S16) showed a weak and non-monotonic trend with increasing *Q*. KL divergence for the SFS and RMSE for the LD were also small and non-monotonic. Overall, the degree of the rescaling effects under this model was fairly small for beneficial and neutral mutations. Although our error measures were much higher for deleterious mutations, these probably were very noisy estimates that reflect the sparsity of our data rather than a genuinely strong rescaling effect.

### Scaling effects are substantial in a population modeled after *Drosophila* but with stronger selection on beneficial mutations

In the previous section we examined the behavior of a *D. melanogaster* population model with weak selection acting on beneficial mutations (*s*=7.125×10^-6^ (Ragsdale et al. 2016; Lauterbur et al. 2023)). We therefore sought to examine whether the scaling effects remain diminished in our *Drosophila* modeled population when a higher selection coefficient for beneficial mutations is used (*s*=0.01); we refer to this model as the “*Drosophila* model with stronger selection”. Classifier results (Figure S19) indicate that scaling effects were much more drastic when strongly beneficial mutations are present. Accuracy for the random forest model for fixation times and fraction of mutations fixed were 100% for all values of *Q*. Accuracy for LD and SFS features were close to 50% for *Q* of 50 but increased until around 76% and 78% respectively at *Q*=200.

MPE of fixation times (Figure 10A) showed a monotonic increase in error values for all mutation types with increasing *Q*. While deleterious mutations had errors that are smaller and in the opposite direction than the *Drosophila* model with across all values of *Q*, errors for beneficial and neutral mutations were much larger. For instance, neutral mutations exhibited an MPE for average fixation times of ∼12% at *Q*=50 which increased to 66% for *Q*=200. In contrast the fixation time MPEs for neutral mutations under the *Drosophila* model at those same values of *Q* were around −0.6% and −0.8%. MPE for beneficial mutations’ average fixation time in the *Drosophila* stronger-selection model was ∼13% at *Q*=50 and increased to around 71% at *Q*=200, compared to the corresponding values of 0.18% and −0.04% in the *Drosophila* model. Deleterious mutations exhibited errors of ∼12% at *Q*=50 and ∼64% at *Q*=200. We note that unlike the *Drosophila* model, deleterious mutations did occasionally fix at all values of *Q* under the *Drosophila* stronger-selection model likely due to hitchhiking with beneficial mutations, and that the standard deviation from the bootstrapping estimate was much smaller for fixation MPEs, especially for deleterious mutations, compared to the *Drosophila* model.

**Figure 10.**
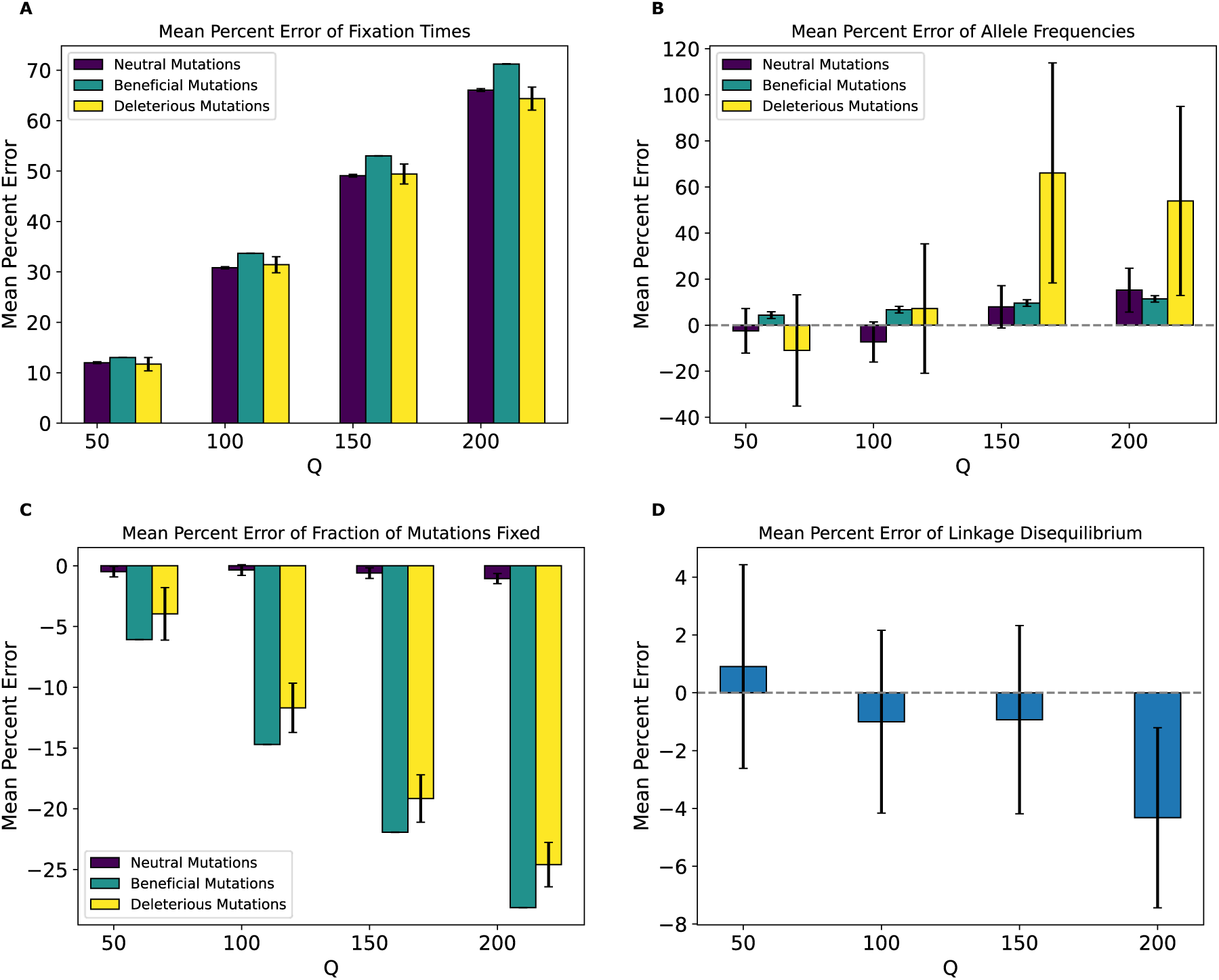
Mean percent error of various statistics for the *Drosophila* stronger-selection model by values of *Q*. **(A)** Mean percent error for average mutation fixation times. **(B)** Mean percent error for average allele frequencies. **(C)** Mean percent error for average fractions of mutations fixed. **(D)** Mean percent error for average linkage disequilibrium as measured by the values of *r*^2^. Values and error bars represent mean and standard deviation of 1000 bootstrapped estimates.

The trend for allele frequency MPE (Figure 10B) was more complex and non-monotonic. For neutral mutations, MPE alternates its sign at each successive *Q*. Deleterious mutations exhibited the highest error at *Q*=150 (around 65%) and a general non-monotonic trend. In contrast, the allele frequency MPE for beneficial increased monotonically from ∼4% at *Q*=50 to ∼11% at *Q*=200. Of note here is that all allele frequency MPEs for this model were higher compared to their corresponding mutation types in the *Drosophila* model, even for deleterious mutations which had a maximum error of around 9% for the *Drosophila* model. For the fraction of mutations fixed (Figure 10C), MPE was lowest for neutral mutations, where the highest magnitude MPE was at *Q*=200 (around −1%). Beneficial and deleterious mutations, on the other hand, had much higher allele frequency MPEs, which were negative and increased in magnitude monotonically with *Q* to ∼-26% for both mutation types at *Q*=200. This was starkly different than the corresponding values for the *Drosophila* model. MPE values for LD (Figure 10D) were also non-monotonic with *Q* as was the case for the *Drosophila* model, but were much higher in magnitude, with the highest value being around −4.3% at *Q*=200, compared to the highest value of around 0.88% at *Q*=200 for the *Drosophila* model.

KL divergence for fixation times (Figure S19) increased monotonically with *Q* to around 1.07, 3.02, and 1.48 for neutral, deleterious, and beneficial mutations respectively, at *Q*=200. On the other hand, KL divergence values for the SFS exhibited different trends for each mutation type; for beneficial mutations, KL divergence increased monotonically from around 0.03 at *Q*=50 to around 0.08 at *Q*=200. For neutral mutations, the trend was somewhat of a monotonic decrease from a value of around 0.5 at *Q*=50 to around 0.31 at *Q*= 200. RMSE for LD increased slightly with *Q* until ∼0.005 at *Q*=200.

### Scaling effects are present in a single selective sweep model

Finally, we sought to examine the effects of scaling in a simple hitchhiking model with only neutral mutations and a single beneficial mutation sweeping through the population. We focus here on the distribution of fixation times for the beneficial mutation across all replicates, and on the SFS and LD for neutral mutations. Classifier accuracy results (Figure S21) showed that scaling also affects these simulations with the intensity of the effects increasing with *Q*. While classifier accuracy was low at *Q*=2, with the best accuracy being ∼64% for LD, accuracy increased with *Q*, especially for LD where accuracy was at 85% for *Q*=5 and reached 100% for *Q*=20. The highest classification accuracy for fixation times and the SFS was also observed at *Q*=20 (73% and 90%, respectively). MPE for fixation times of the beneficial mutation (Figure 11A) were substantial and increased monotonically with *Q* from −0.7% at *Q*=2 to ∼32% at *Q*=20. MPEs for the neutral mutation allele frequency also increases with *Q* but were smaller, reaching only as high as ∼1.3% at *Q*=20. For LD, we also observes a monotonic increase in MPE from ∼0.66% at *Q*=2 to ∼16% at *Q*=20. KL divergence for the distribution of fixation times of the beneficial mutation (Figure S21) was non-monotonic and was ∼0.41 at *Q*=20. For neutral mutations, the SFS KL divergence and LD RMSE for both increased monotonically with *Q*, with the highest KL divergence and RMSE values being 0.32 and 0.001 respectively at *Q*=20. These results demonstrate that even with a scenario involving a single sweep, scaling can bias simulation outcomes, especially fixation times for the sweeping mutation and LD between pairs of linked neutral mutations.

**Figure 11.**
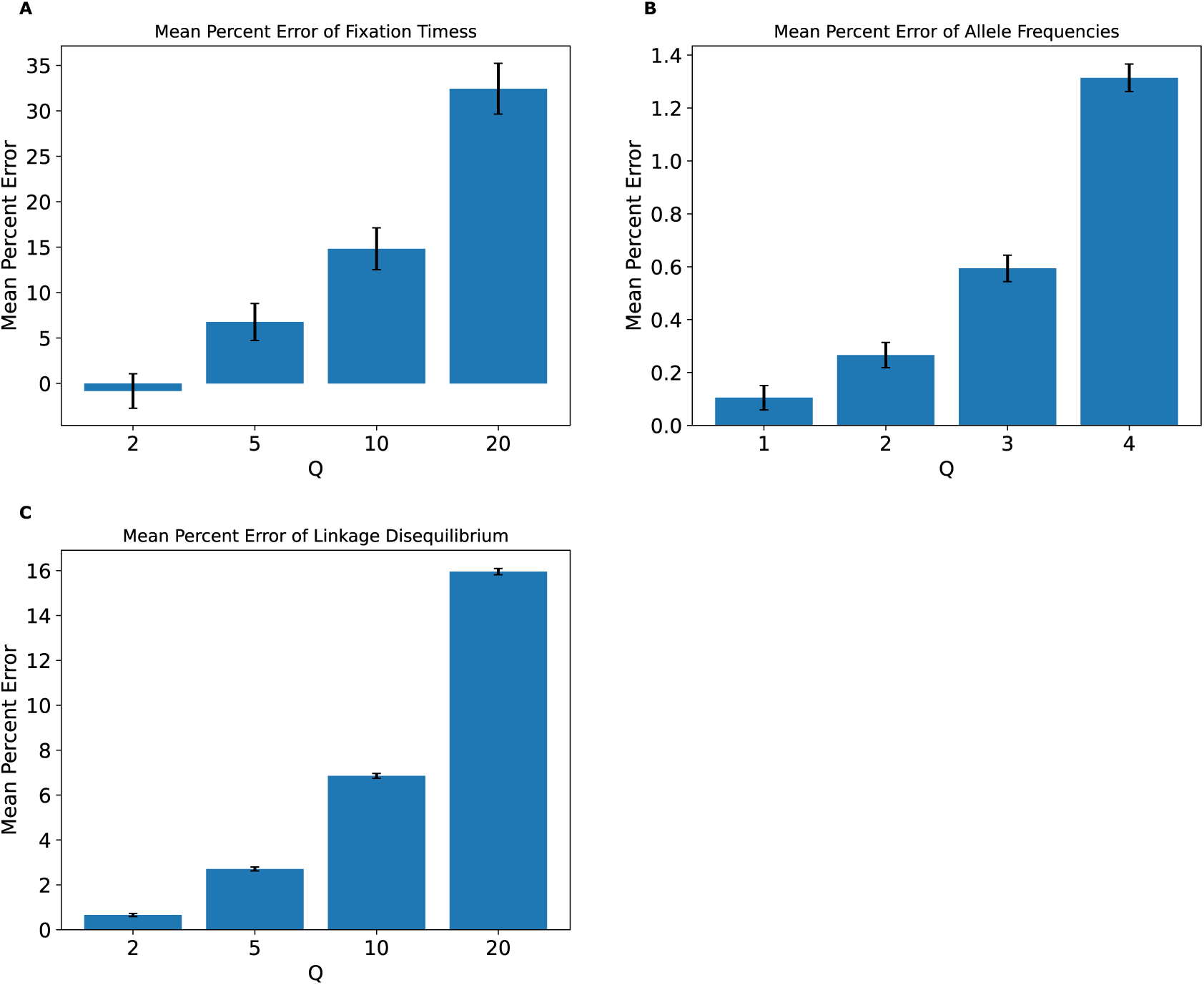
Mean percent error of various statistics for the single sweep model by values of *Q*. **(A)** Mean percent error for average beneficial mutation fixation times. **(B)** Mean percent error for average allele frequencies for neutral mutations. **(C)** Mean percent error for average linkage disequilibrium for neutral mutations as measured by the values of *r*^2^. Values and error bars represent mean and standard deviation of 1000 bootstrapped estimates.

### Scaling effects can often be recapitulated from a small number of replicates

Although some general conclusions can be drawn from our experiments above (e.g., rescaling effects are common and generally increase with *Q*), behavior differs substantially across models and thus it may be difficult for a researcher to know what the impact of rescaling would be on their simulations without conducting similar experiments themselves on the model of interest. Since it is often unfeasible to acquire many replicates of minimally-scaled (ideally unscaled) simulations under some models due to computational limitations, we sought to investigate whether statistical measures of the impact of scaling such as MPE, KL divergence, and RMSE can be reasonably approximated with a smaller number of simulation replicates for a subset of our simulations. We therefore randomly sampled smaller sets of replicates for the full model, the no-beneficials model, and the *Drosophila* model with strong beneficials for each value of *Q*. Our results indicated that when information about the outcome of interest is sufficiently dense in each replicate, measures of deviations gleaned from a smaller number of replicates were generally close to those obtained using all 1000 replicates. For example, in a sample of 100 replicates simulated under the full model (Figure S22), all measures of deviation were extremely similar to those obtained using 1000 replicates, with the exception of KL divergence for the SFS of beneficial mutations, which was slightly higher for all values of *Q* compared to the results from 1000 replicates, and had the highest variance compared to other mutation types. When using only 10 replicates (Figure S23), deviation outcomes for the full model were still informative but less well preserved and predictably showed higher variance. For instance, while the bootstrapping means for the fixation times MPEs were similar when using 10 replicates to when using the full 1000 replicates, the variance was much higher compared to when using 100 or 1000 replicates. KL divergence for fixation times and the SFS was more severely affected, especially for neutral and deleterious mutations at *Q*=20 and also had higher variance compared to using 100 replicates, whereas the average RMSE for LD was still well-preserved and had slightly less variance than other deviation measures.

For the no-beneficials model, using 100 replicates (Figure S24) preserved all measures of deviation rather well, with the exception of slightly higher KL divergence for fixation times of neutral mutations at *Q*=20 compared to that estimated from 1000 replicates. Using 10 replicates only was sufficient to recover similar estimates of MPE for average fixation times, fraction of mutations fixed, and LD, but induced a more marked change in KL divergence of fixation times and the SFS (Figure S25). The average RMSE for LD, however, remained similar when decreasing the number of replicates. For almost all simulation outcomes, variance estimates obtained by repeating the replicate subsample process with 10 replicates were higher than those obtained from 100 replicate samples.

Unfortunately, down-sampling the number of replicates for the *Drosophila* stronger-selection model did not recapitulate the measured errors for some outcomes as well. While down-sampling to 100 replicates (Figure S26) does conserve some error measurements, such as the average fixation time MPEs for neutral and beneficial mutations, the average fraction of mutations fixed MPE for beneficial mutations, and the average allele frequency MPE for beneficial mutations, we observed substantial shifts for all other measurements, including KL divergence for fixation times and the SFS for all mutation types and the average LD RMSE. Further down-sampling to 10 replicates (Figure S27) caused further loss of ability to capture all measures of error. Additionally, the estimates of variance from repeating the sampling process were much higher, even when sampling 100 replicates, compared to the full or no-beneficials models. This result was not unexpected, as we simulated much smaller regions in the *Drosophila* genome to examine smaller scaling factors (Methods), and this resulted in a much smaller number of mutations available for analysis (e.g., the estimated SFS was based on fewer than 20 mutations present in a sample for the *Drosophila* stronger-selection model on average), resulting in nosier (and possibly smaller) rescaling effects that exhibit greater variance at smaller sample sizes.

In summary, using a smaller number of replicates can be informative for characterizing deviation between scaled and unscaled simulations when replicates are sufficiently information dense (e.g., there are enough polymorphisms in the sample taken at the end of the simulation to estimate the SFS, or there were enough fixations during the simulation to accurately estimate the fraction of mutations fixed). Using a much smaller number of replicates can impair these estimates, especially for KL divergence, and most severely for simulation models whose replicates contain sparse information such as a low number of segregating or fixed polymorphisms (e.g., our simulations of 10 kb regions in the *Drosophila* genome).

## DISCUSSION

Population genetic simulation software is increasingly powerful and flexible (Kelleher et al. 2016; Baumdicker et al. 2022). This is especially so for forward-in-time simulators that allow for complex scenarios involving different types of natural selection (Thornton 2014; Haller and Messer 2017) departures from Wright-Fisher assumptions (Haller and Messer 2019), the modeling of multispecies dynamics (Haller and Messer 2023), and more. Unfortunately, because forward simulations require the entire population to be modeled, they can be too computationally costly when the population size is large. Rescaling the simulation parameters to simulate a smaller population to model a larger one is therefore necessary in many cases. Unfortunately, it has been shown that this rescaling trick can bias simulation outcomes (Uricchio and Hernandez 2014; Adrion et al. 2020). However, the rescaling effect has only been examined for a small number of simulation scenarios, and thus we still do not know the manner and magnitude with which it will affect simulation results.

Here, we sought to investigate the rescaling effect on a wider array of simulation models, including those of varying genome densities, population sizes, and strengths of positive and negative selection. We used `stdpopsim` and `SLiM` to generate these simulations under varying values of the rescaling factor *Q*, and record a set of simulation outcomes summarizing the SFS, LD, the faction of mutations fixed, and time to fixation. Ideally, the rescaling strategy would produce results that are difficult to distinguish from unscaled simulations. However, this was not the case—for most of the models that we examined, binary classifiers were able to discriminate between rescaled and unscaled simulations quite easily. For example, for our full model, classifiers examining either the estimated fraction of mutations that fixed and the distribution of fixation times could label simulation replicates as unscaled or rescaled by a factor of only *Q*=2 with 100% accuracy (Figure 2). Classifiers using the SFS or LD also performed well under this model with only a modest degree of rescaling (e.g., nearly 100% accuracy at *Q*=5). Our classifiers generally performed well for the other models examined here, but there were two notable exceptions: 1) for the strictly neutral model, only the LD-based classifier could discriminate between unscaled and scaled simulations with 100% accuracy at *Q*=20 (the highest rescaling factor examined for this model; see Figure S10); 2) for the *Drosophila* model, none of the classifiers performed well at any value of *Q* examined (although this was not the case for the *Drosophila* stronger-selection model). Thus, rescaling appears to significantly affect simulation outcomes, a result that has been corroborated by a concurrent study by Ferrari et al. (2024).

Even if rescaling alters simulations sufficiently to be detected by classifiers, the size of this rescaling effect may still in some cases be small enough to be tolerable by researchers. We therefore sought to more directly quantify the magnitude of rescaling effects by measuring the mean percent error between scaled and unscaled simulations for each of our simulation outcomes. We found that MPE between scaled and unscaled outcomes was often substantial. For example, the fraction of deleterious mutations that reached fixation under our no-beneficials model was overestimated by ∼140% when *Q*=20 (Figure 4C). However, MPE did vary considerably across simulation outcomes and models (see Tables 4,5 for a summary of all the MPE values for all models in this study, and Tables S1, S2 for all KL divergence values and classifier accuracies). Consider again the MPE for fraction of mutations fixed under the no-beneficials model.

When we added beneficial mutations into this model (yielding the “full model”; Methods), the MPE for fraction of deleterious mutations fixed was only −0.12% at *Q*=20 (Figure 3C). Furthermore, the no-beneficials sparse model experienced smaller biases due to scaling compared to the no-beneficials model, with the MPE for fraction of deleterious mutations fixed being ∼37% in this case. This suggests that simulations containing deleterious mutations without many beneficial mutations may be more prone to scaling biases, and that the magnitude of this bias scales with the total number of deleterious mutations present.

We note that classification accuracy did not always correspond with the observed trends in MPE. For instance, in the no-beneficials sparse model, the random forest classifier achieved higher accuracy on the SFS features than fixation times at *Q*=5, despite allele frequencies having an MPE that is lower in magnitude than that of fixation times. Thus, one should bear in mind that our classification results are indicative of the degree to which the scaled and unscaled simulations are separable—which could be the case for two distributions that have only a slight difference in mean value but low variance—while our estimates of MPE and KL divergence/RMSE more directly speak to the magnitude of the rescaling effect.

Our analysis of the variety of simulation models considered here does yield some conclusions that are likely to generalize to other models. For one, we might expect the rescaling effect to be more severe in smaller populations than in larger ones. For example, the magnitude of the rescaling effect observed in our no-beneficials model (Figure 4) was reduced somewhat when we re-ran these simulations with a population that was twice as large (Figure 5). We also observed a weaker rescaling effect in the simulated *Drosophila* population, although this was perhaps in large part a result of the sparsity of mutation data in these simulations which modeled smaller chromosomes (see below). However, the rescaling effect may still be appreciable in larger populations when positive selection plays a more prominent role and when *Q* is large: the rescaling effect in the *Drosophila* stronger-selection model was far greater than that under the *Drosophila* model (compare Figure 10 to Figure S16).

Intuitively, it is not unexpected that the rescaling effect would be greater in magnitude in the presence of selection. For example, in the presence of positive selection, the rationale for rescaling the selection coefficient by *Q* is not supported by theory, as neither the time to fixation nor the fixation probability are expected to be a linear function of the population-scaled selection coefficient 2*Ns* (Kimura 1962; Charlesworth 2020)—our results from simple simulations of a single selective sweep underscore this (Figure 11). Similarly, Uricchio and Hernandez (2014) showed that the theoretical prediction that the impact of a sweep on a linked variation is determined by 2*Ns* (Wiehe and Stephan 1993) is not valid when *s* is large, which will be the case when s is multiplied by *Q* during rescaling. It has also been shown that the degree to which neutral genetic diversity at a given site is reduced by both selection against linked (Nordborg et al. 1996) and genome-wide (Hudson and Kaplan 1995) deleterious mutations is dependent on *s* and not *Ns*, meaning that rescaling will produce inaccurate results for such models (as pointed out by Matheson and Masel (2024)).

Moreover, we would expect there to be a stronger rescaling effect in the presence of large amounts of natural selection because rescaling will cause each simulated genome to receive a larger number of selected mutations each generation, which may alter short-term dynamics before the increased recombination rate can break up this extra linkage. As the results for our no-beneficials model (Figure 4), the no-beneficials sparse model (Figure 6), and both of our *Drosophila* models (Figure 10 and Figure S16) show, larger numbers of deleterious mutations—even in the absence of beneficial mutations—and a higher strength of selection in general appear to compound rescaling effects. The addition of large amounts of positive selection to simulation models containing a large density of deleterious mutations may also qualitatively alters the rescaling effect, as seen by comparing the results of our full model (Figure 3) to those of the no-beneficials model (Figure 4). For example, in the no-beneficials model we observed large skews in the fixation times for neutral and deleterious mutations that increase with *Q*. This is not observed in the full model, where instead we observed a large reduction in the fraction of beneficial mutations that reach fixation as *Q* increases, among other differences.

Finally, rescaling divides the total number of generations by *Q*, thereby reducing the temporal resolution of the simulation when it comes to events that occur on short timescales. For example, all estimates of selective sweep sojourn times will necessarily be rounded up to the nearest multiple of *Q*, resulting in underestimates. This reduced temporal resolution may also impact the shape of genealogies: as *Q* increases, we expect the lengths of genealogical branches to decrease proportionally, but eventually those shortest branches in the tree will reach their minimum length (1 generation) while the longest branches continue to shrink with increasing *Q*. This could in some cases result in biased branch length distributions, potentially skewing the site frequency spectrum. A corollary of all these lines of reasoning is that the rescaling effect should be less severe under neutral simulations, perhaps especially those of large populations. However, although the effect in our strictly neutral model was more subtle, we do observe one here (Figure 7). We note that this finding is consistent with results from Adrion et al. (2020) who observed biased estimates of several summary statistics when performing purely neutral rescaled forward simulations of the demographic model from Kamm et al. (2020). Together, these results, combined with the fact that the values of *Q* examined here are similar to those commonly used in studies relying on forward simulation (Methods), suggest that some previously published results may have been impacted by some degree of rescaling effect.

One important limitation of our study is that we measured the rescaling effect only in Wright-Fisher simulations. As software programs like `SLiM` continue to add functionality an allow researchers to simulate increasingly complex non-Wright Fisher models, there will be interest in accelerating these simulations as well, perhaps in part via parameter rescaling. For instance, simulations involving spatial interactions or cross-species dynamics may need rescaling of the parameters governing such interactions. There are not yet standard approaches for rescaling the parameters of these models, and thus it is unclear how rescaling such simulations could bias their results. It is therefore important to carry out similar studies investigating the effect of any possible scaling methods for models that fall outside of the scope of the Wright-Fisher models examined in this study.

Although we find that the consequences of population-size rescaling for forward simulations are somewhat predictable, when examining the totality of our results one cannot help but notice numerous exceptions to the general trends described above. Indeed, as observed for several simulation models and outcomes, we did not always find that the magnitude of MPE increased monotonically with *Q*. Thus, it is not as of yet possible to predict *a priori* precisely, for a population genetic model of interest, which simulation outcomes will be most affected by rescaling, and whether these outcomes will be upwardly or downwardly biased for a given value of *Q*. Given the potential of bias and the difficulty in predicting its direction and magnitude when scaling, researchers may consider other alternatives for alleviating the computational burden of simulations. For instance, tree-sequence recording (Kelleher et al. 2018; Haller et al. 2019) allows for the omission of neutral mutations during simulation time, which can then be quickly added afterwards using the generated tree-sequence, significantly decreasing simulation run time. This method also allows for neutral burn-in to be performed after (via coalescent “recapitation”; (Haller et al. 2019)) rather than during simulations. However, the approach of performing a neutral burn-in could also lead to biases for simulations models with high rates of selected mutations. Alternative approaches could also in principle be used to accelerate certain phases forward simulations (e.g. simulated allele frequency trajectories using either discrete or diffusion models; Gutenkunst et al. 2009; Ewing and Hermisson 2010; Ragsdale and Gutenkunst 2017) but such methods are currently limited to modeling the dynamics of very small numbers of mutations and thus will not be helpful in most instances where researchers are interested in genetic variation along a large recombining chromosomal region.

Thankfully, for researchers still needing to utilize scaling during simulations, we have found that even a relatively small number of simulation replicates (e.g., 10, for the cases shown in Figure S23, Figure S25, and Figure S27) can be enough to accurately describe the impact of rescaling both qualitatively and quantitatively (via MPE) for a given *Q*. When using this strategy we generally obtained a similar point estimate, although the variance around this estimate was higher, as expected. Thus, a small-scale test of varying *Q* values and assessing the direction and magnitude of the resulting bias for any outcome of interest may be sufficient for a researcher wishing to determine which value of *Q* will provide the best balance between computational efficiency and accuracy of estimated outcomes—this tradeoff is discussed below. For researchers wishing to simulate very large numbers of simulation replicates each of reasonably small chromosome lengths (e.g., for ABC inference of models involving natural selection (Johri et al. 2020)), this will mean just simulating a small number of replicates for each of a range of rescaling factors. For researchers instead hoping to generate chromosome-scale simulation replicates, it may be possible to first do a small-scale test on shorter chromosomal regions to select a value of *Q* before conducting the full-scale study.

Although the approach of conducting a smaller-scale simulation study can reveal the impact of a given *Q*, the best value of *Q* for a particular study will depend on the specific goals of the researcher (tolerance for bias, size of sim dataset needed, etc). We therefore abstain from proposing a general strategy for selecting an optimal *Q* here. However, we emphasize that despite our main conclusion that rescaling effects are pervasive, our results do indicate that some degree of rescaling will likely be acceptable for most applications. For example, the smallest degree of rescaling possible, *Q*=2, typically has a very small rescaling effect for our human-inspired simulation models (MPE < 1% in many cases), but could still yield a considerable speedup (∼3-fold on average; see Table S3 for average running times for select models). Higher degrees of rescaling will yield much larger speedups, and in some cases produce results that some users may find acceptable. For example, in our full model, using *Q*=5 results in MPEs that are less than 15% for most simulation outcomes (with the fraction of beneficial mutations that reached fixation being the one exception; Figure 3), and this *Q* yields a ∼14-fold speedup. For relatively simple evolutionary scenarios, more theoretically motivated algorithmic approaches to selecting an optimal *Q* value may be possible (Uricchio and Hernandez 2014).

Further studies should also evaluate alternative approaches to maximize the benefit of rescaling while minimizing its consequences, such as changing *Q* dynamically throughout a simulation so that it is elevated during phases where rescaling effect is less likely to be a concern (e.g., the population is very large), and then decreasing it at key points where the rescaling effect could be severe (e.g., during severe population bottlenecks). Although it may also be possible in principle to “correct” the rescaling effect on simulation outcomes once the magnitude and direction of this effect has been determined, our results show that this would not be straightforward. For example, while one could try to correct simulation outcomes by shifting the distribution based on the MPE, our results show that changes in mean outcomes do not sufficiently describe the rescaling effect (e.g., for the full model in Figure 3, MPE for fixation times decreased when *Q* increased from 5 to 10, but KL divergence increased, demonstrating that the change in the distribution is more complex than a simple shift in the mean).

In summary, although population rescaling may in many cases be unavoidable for the time being, researchers may be able to mitigate the undesirable effects of rescaling by carefully considering and investigating them before performing simulations at scale. Still, additional studies are needed to characterize the manner in which population rescaling affects simulation results and the models affected. Indeed, our results, together with those of Ferrari et al. (2024) demonstrate that such work will be essential to ensure that the continuing advances in simulation power and flexibility are being used to produce accurate results.

## DATA AVAILABILITY

All code for performing simulations and analyzing/visualizing their results can be found at https://github.com/SchriderLab/simscale-snakemake.

## ACKNOWLEDGMENTS

The authors thank members of the PopSim Consortium for feedback on preliminary results from this study. We also thank Austin Daigle, Ben Haller, and two anonymous reviewers for feedback on the manuscript. This work was funded by NIH awards R01HG010774 and R35GM138286.

## Notes

### Competing Interest Statement

The authors have declared no competing interest.

### Summary of Updates

The revised version of this manuscript contains several additional analyses (including bootstrap resampling to estimate the variance of simulation outcomes, re-run simulations to ensure that the parameters--including mutation/recombination rates and chromosome lengths--exactly match those reported in the manuscript, etc) and summaries of results (including simulation run times).

