## Supplementary Material for "Population size rescaling significantly biases outcomes of forward-in-time population genetic simulations"

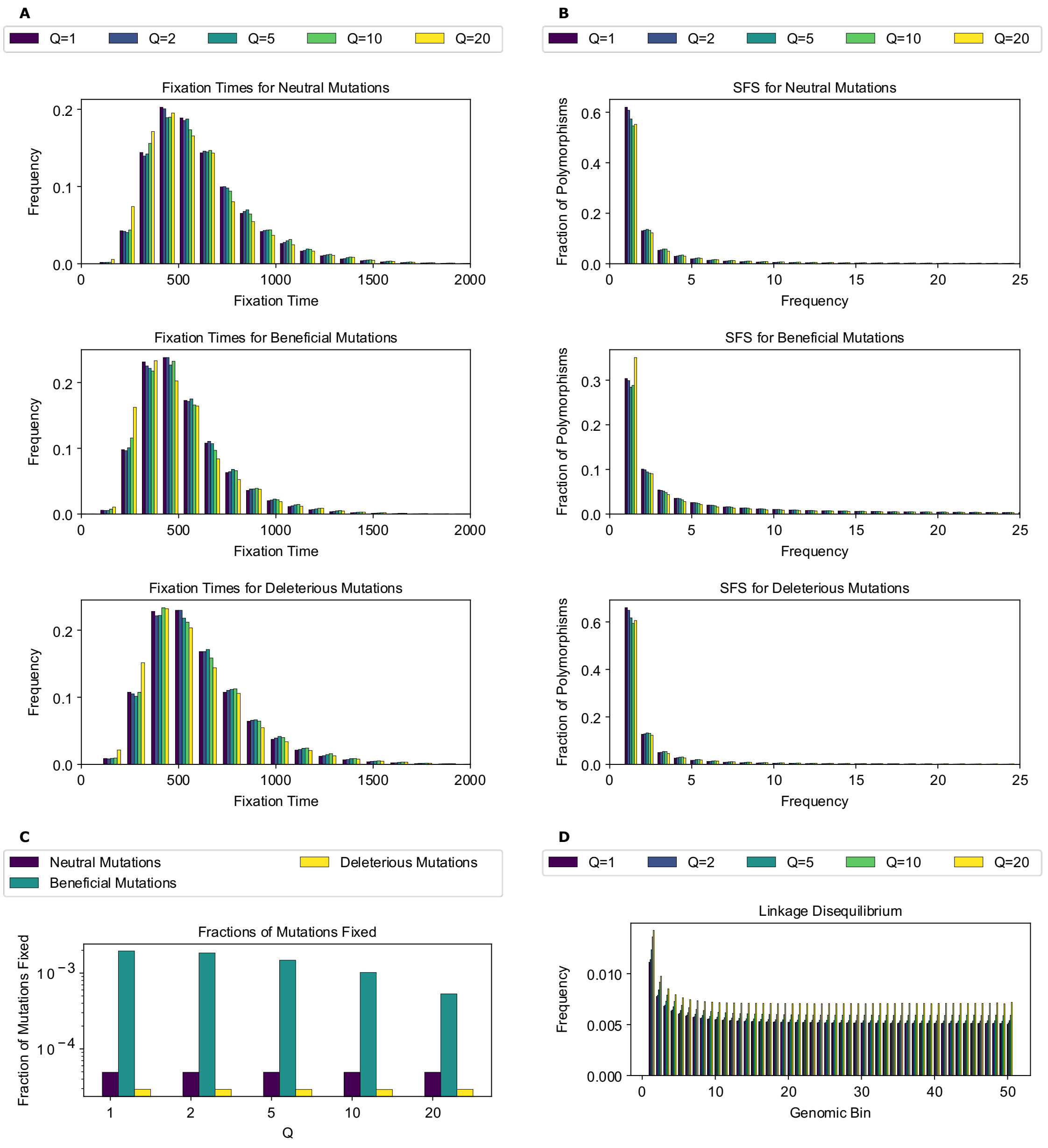


**Figure S1. Simulation outcomes for the full model.**
**(A)** Fixation time distributions for all mutation types. **(B)** Site frequency spectra for all mutation types. **(C)** Fraction of mutations fixed for all mutation types. **(D)** Linkage disequilibrium across the chromosome for 50 genomic bins, as measured by $r^2$. Each outcome was averaged across all replicates. For fixation times, SFS, and LD, this was achieved by averaging each distribution bin across the replicates.

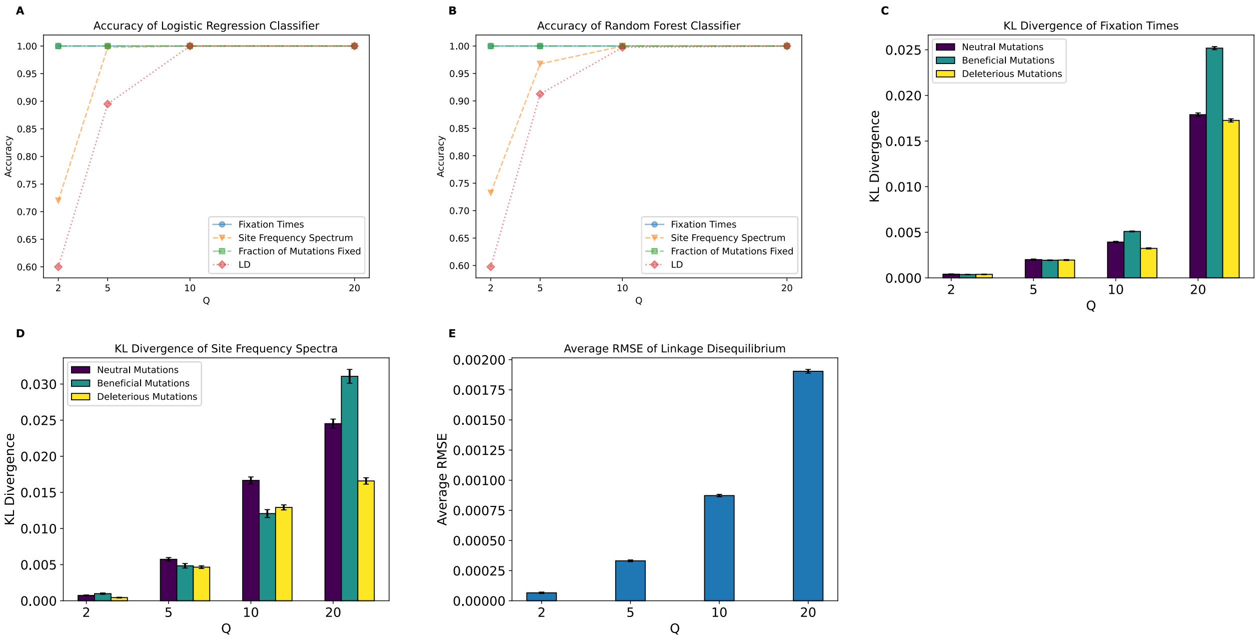


**Figure S2. Metrics of the effect of scaling for the full model.**
**(A)** Accuracy of the logistic regression classification using features subsets for each scaling factor. **(B)** Accuracy of random forest classification using feature subsets for each scaling factor. **(C)** KL divergence of fixation time distributions for all mutation types. **(D)** KL divergence for the SFS for all mutation types. (**E)** Average RMSE of LD. Classifier accuracy represents the proportion of replicates correctly identified by the model as scaled and unscaled using the feature subsets for each $Q$. Values and errors bars for panels **(C)**, **(D)**, and **(E)**, represent mean and standard deviation of 1000 bootstrapped samples.

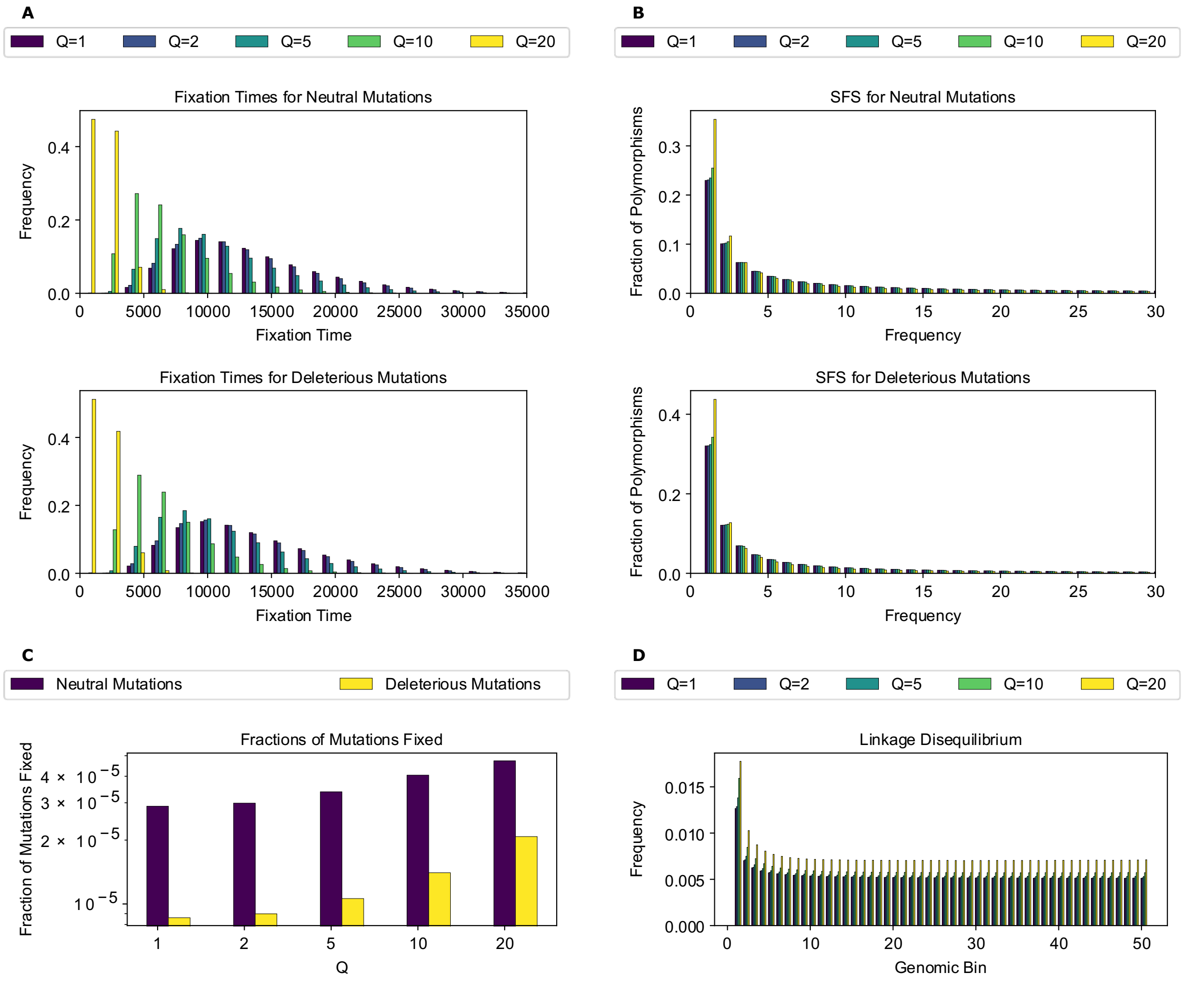


**Figure S3**. **Simulation outcomes for the no-beneficials model.**
**(A)** Fixation time distributions for all mutation types. **(B)** Site frequency spectra for all mutation types. **(C)** Fractions of mutations fixed for all mutation types. **(D)** Linkage disequilibrium across the chromosome for 50 genomic bins, as measured by $r^2$.

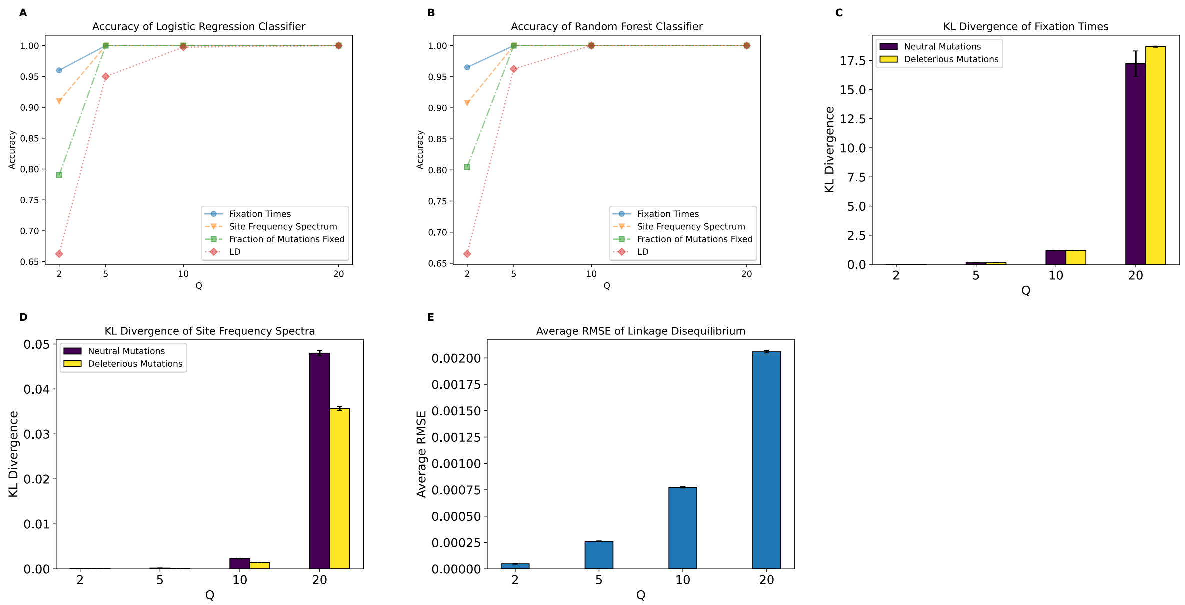


**Figure S4. Metrics of the effect of scaling for the no-beneficials model.**
**(A)** Accuracy of the logistic regression classification using features subsets for each scaling factor. **(B)** Accuracy of the random forest classification using feature subsets for each scaling factor. **(C)** KL divergence of fixation time distributions for all mutation types. **(D)** KL divergence for the SFS for all mutation types. **(E)** Average RMSE of LD. Values and errors bars for panels **(C)**, **(D)**, and **(E)**, represent mean and standard deviation of 1000 bootstrapped samples.

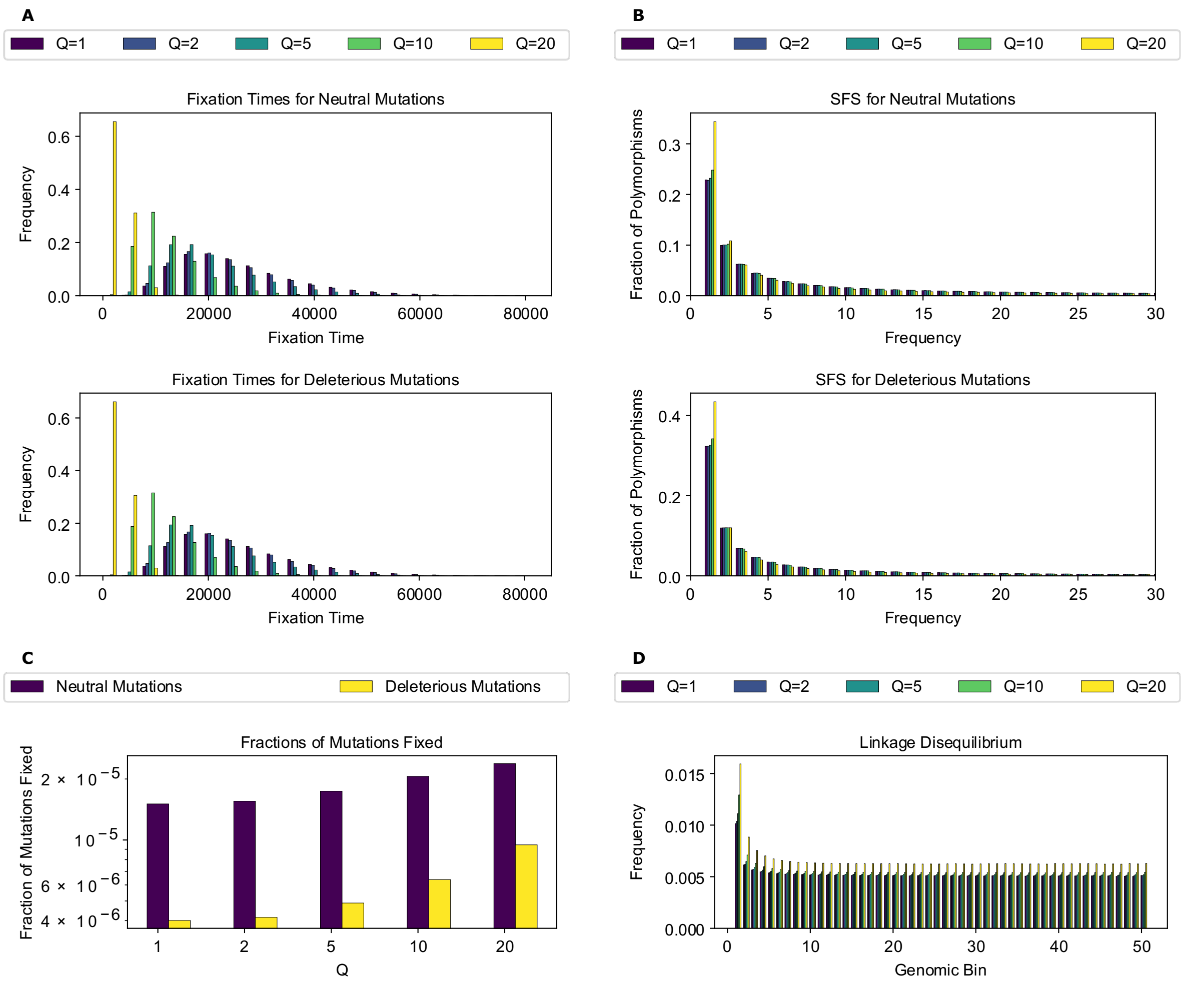


**Figure S5. Simulation outcomes for the no-beneficials larger population model.**
**(A)** Fixation time distributions for all mutation types. **(B)** Site frequency spectra for all mutation types. **(C)** Fractions of mutations fixed for all mutation types. **(D)** Linkage disequilibrium across the chromosome for 50 genomic bins, as measured by $r^2$.

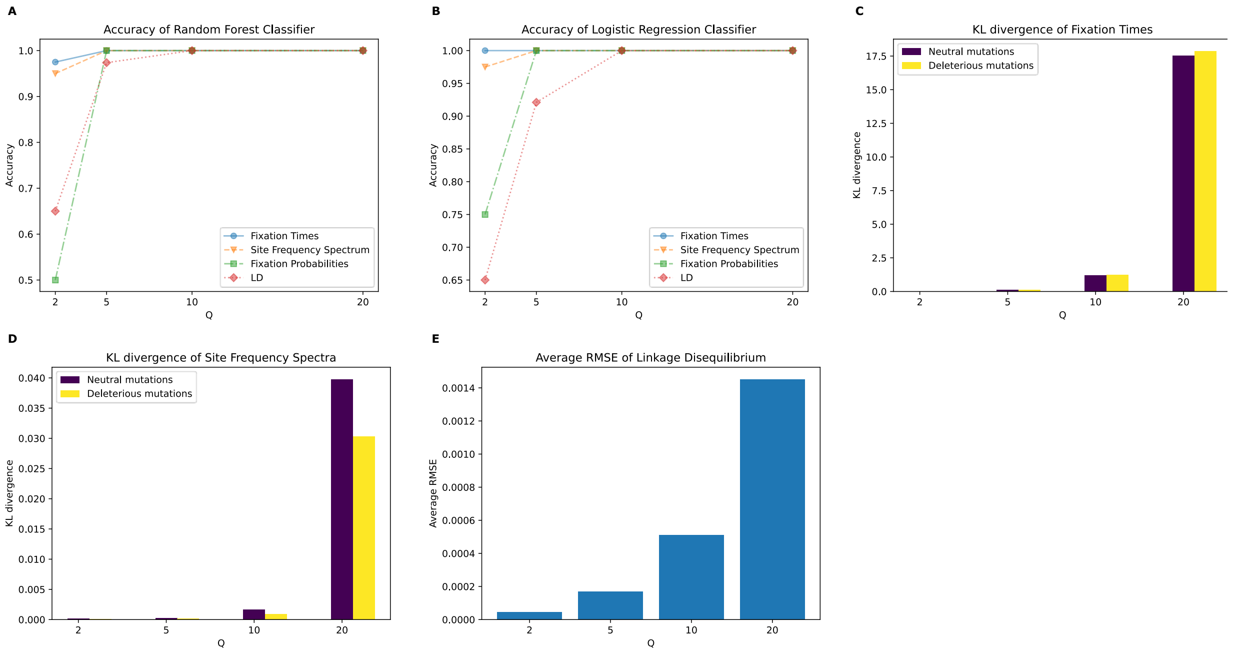


**Figure S6. Metrics of the effect of scaling for the no-beneficials larger population model.**
**(A)** Accuracy of the logistic regression classification using features subsets for each scaling factor. **(B)** Accuracy of the random forest classification using feature subsets for each scaling factor. **(C)** KL divergence of fixation time distributions for all mutation types. **(D)** KL divergence for the SFS for all mutation types. **(E)** Average RMSE of LD. Values and errors bars for panels **(C)**, **(D)**, and **(E)**, represent mean and standard deviation of 1000 bootstrapped samples.

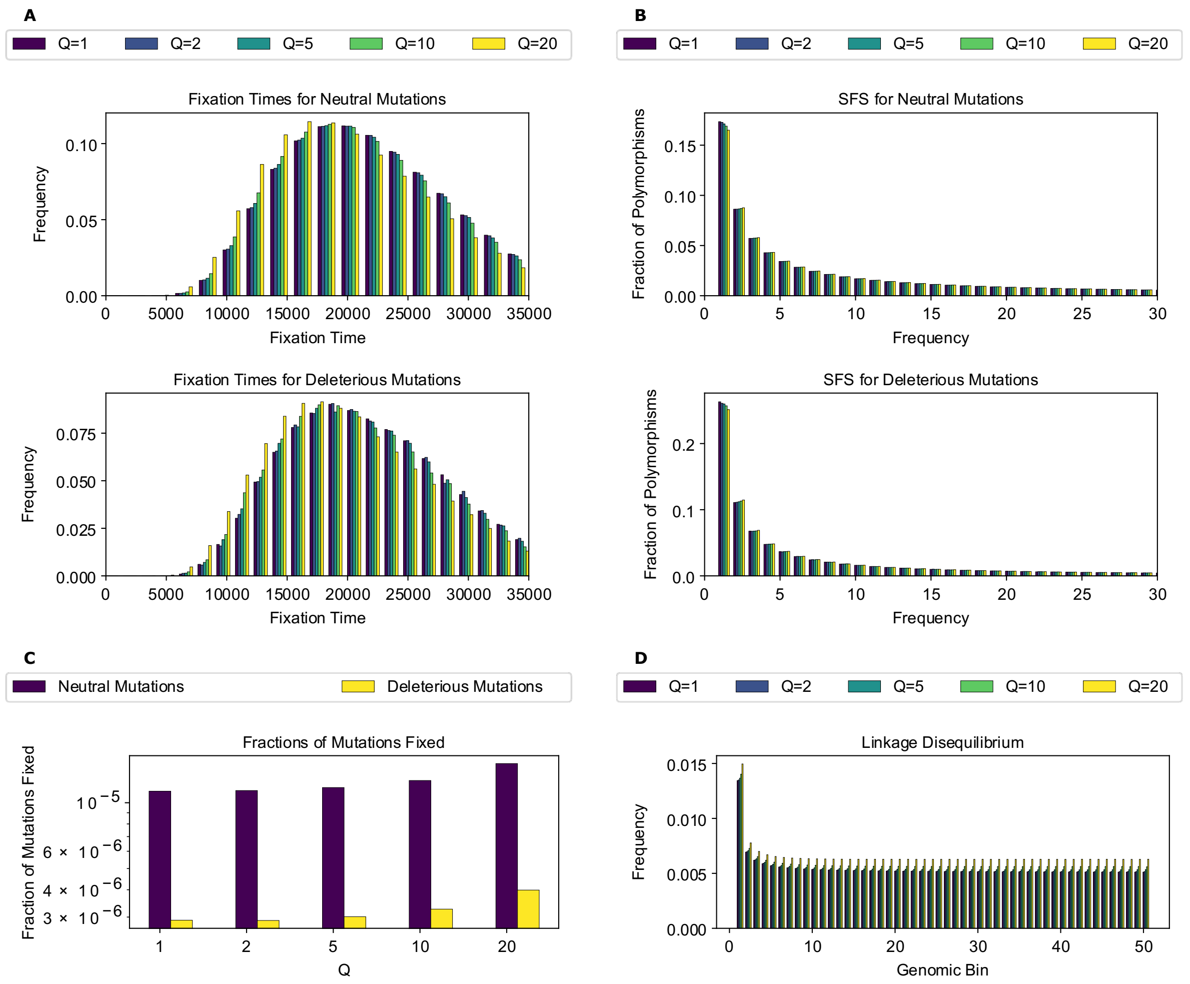


**Figure S7. Simulation outcomes for the no-beneficials sparse model.**
**(A)** Fixation time distributions for all mutation types. **(B)** Site frequency spectra for all mutation types. **(C)** Fractinos of mutations fixed for all mutation types. **(D)** Linkage disequilibrium across the chromosome for 50 genomic bins, as measured by $r^2$.

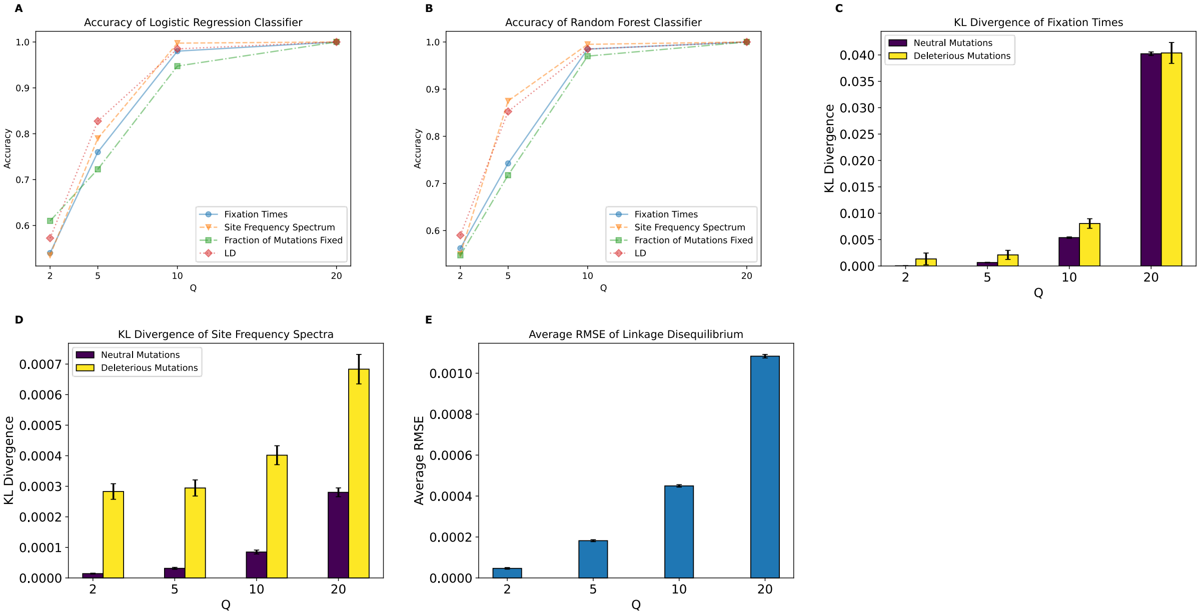


**Figure S8. Metrics of the effect of scaling for the no-beneficials sparse model.**
**(A)** Accuracy of the logistic regression classification using features subsets for each scaling factor. **(B)** Accuracy of the random forest classification using feature subsets for each scaling factor. **(C)** KL divergence of fixation time distributions for all mutation types. **(D)** KL divergence for the SFS for all mutation types. **(E)** Average RMSE of LD. Values and errors bars for panels **(C)**, **(D)**, and **(E)**, represent mean and standard deviation of 1000 bootstrapped samples.

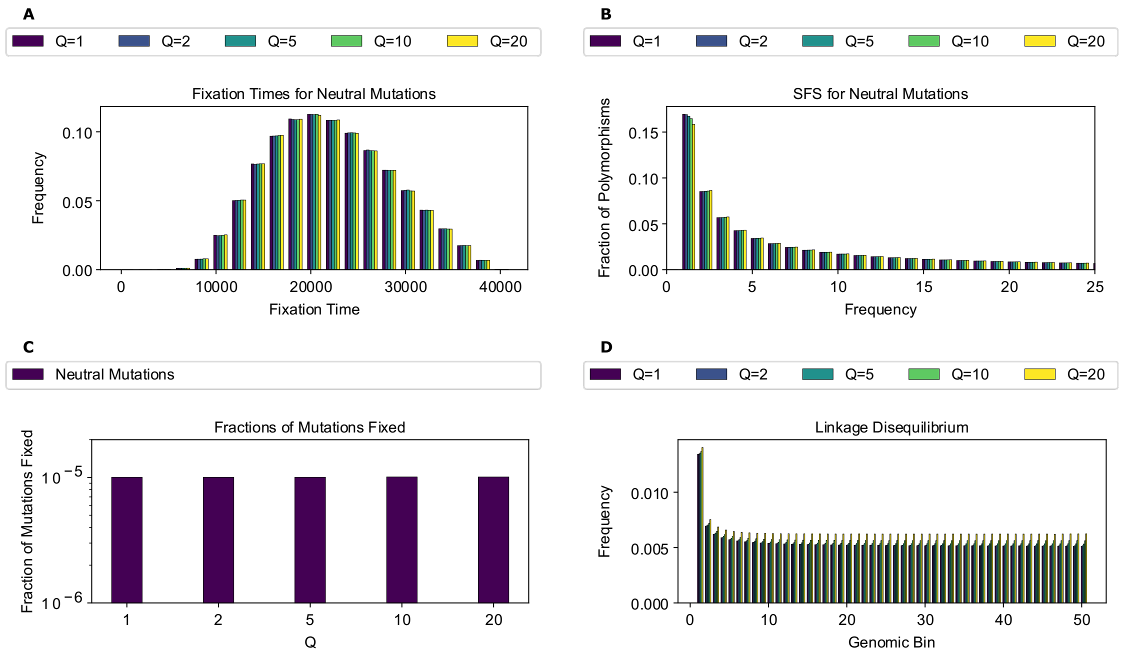


**Figure S9. Simulation outcomes for the strictly neutral model.**
**(A)** Fixation time distributions for neutral mutations. **(B)** Site frequency spectra for neutral mutations. **(C)** Fraction of neutral mutations fixed. **(D)** Linkage disequilibrium across the chromosome for 50 genomic bins, as measured by $r^2$.

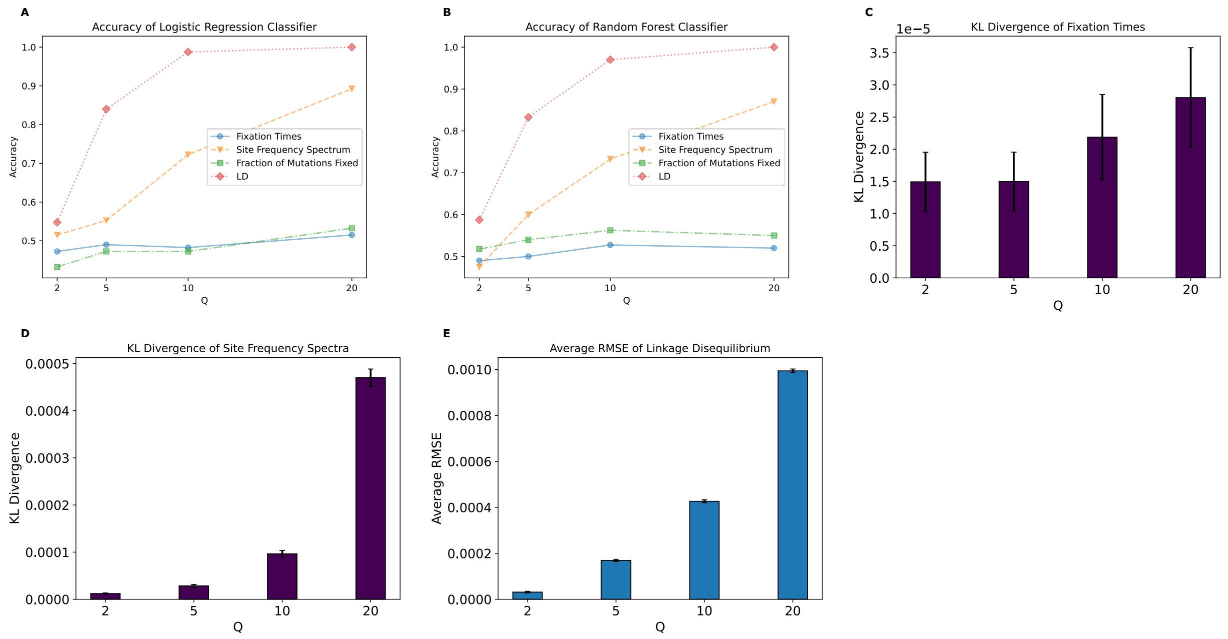


**Figure S10. Metrics of the effect of scaling for the strictly neutral model.**
**(A)** Accuracy of the logistic regression classification using features subsets for each scaling factor. **(B)** Accuracy of the random forest classification using feature subsets for each scaling factor. **(C)** KL divergence of fixation time distributions for neutral mutations. **(D)** KL divergence for the SFS for neutral mutations. **(E)** Average RMSE of LD for neutral mutations. Values and errors bars for panels **(C)**, **(D)**, and **(E)**, represent mean and standard deviation of 1000 bootstrapped samples.

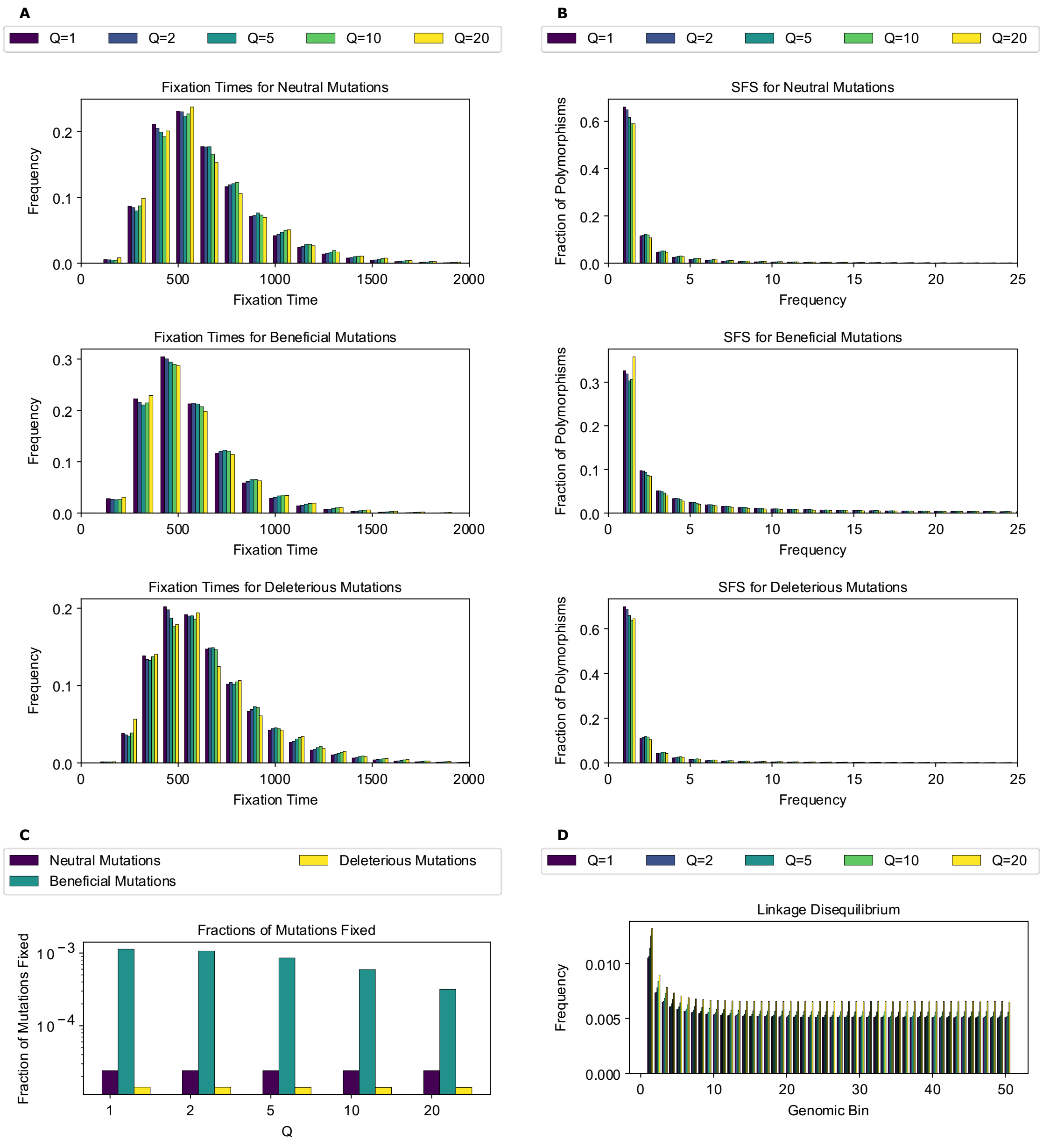


**Figure S11. Simulation outcomes for the population expansion model.**
**(A)** Fixation time distributions for all mutation types. **(B)** Site frequency spectra for all mutation types. **(C)** Fractions of mutations fixed for all mutation types. **(D)** Linkage disequilibrium across the chromosome for 50 genomic bins, as measured by $r^2$.

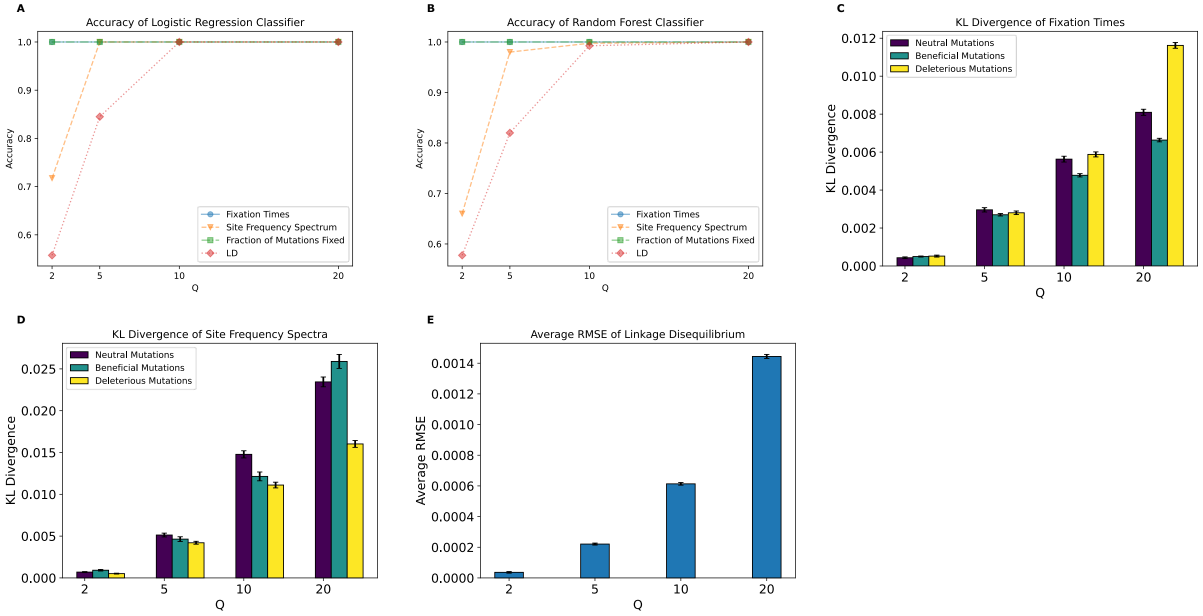


**Figure S12. Metrics of the effect of scaling for the population expansion model.**
**(A)** Accuracy of the logistic regression classification using features subsets for each scaling factor. **(B)** Accuracy of the random forest classification using feature subsets for each scaling factor. **(C)** KL divergence of fixation time distributions for all mutation types. **(D)** KL divergence for the SFS for all mutation types. **(E)** Average RMSE of LD. Values and errors bars for panels **(C)**, **(D)**, and **(E)**, represent mean and standard deviation of 1000 bootstrapped samples.

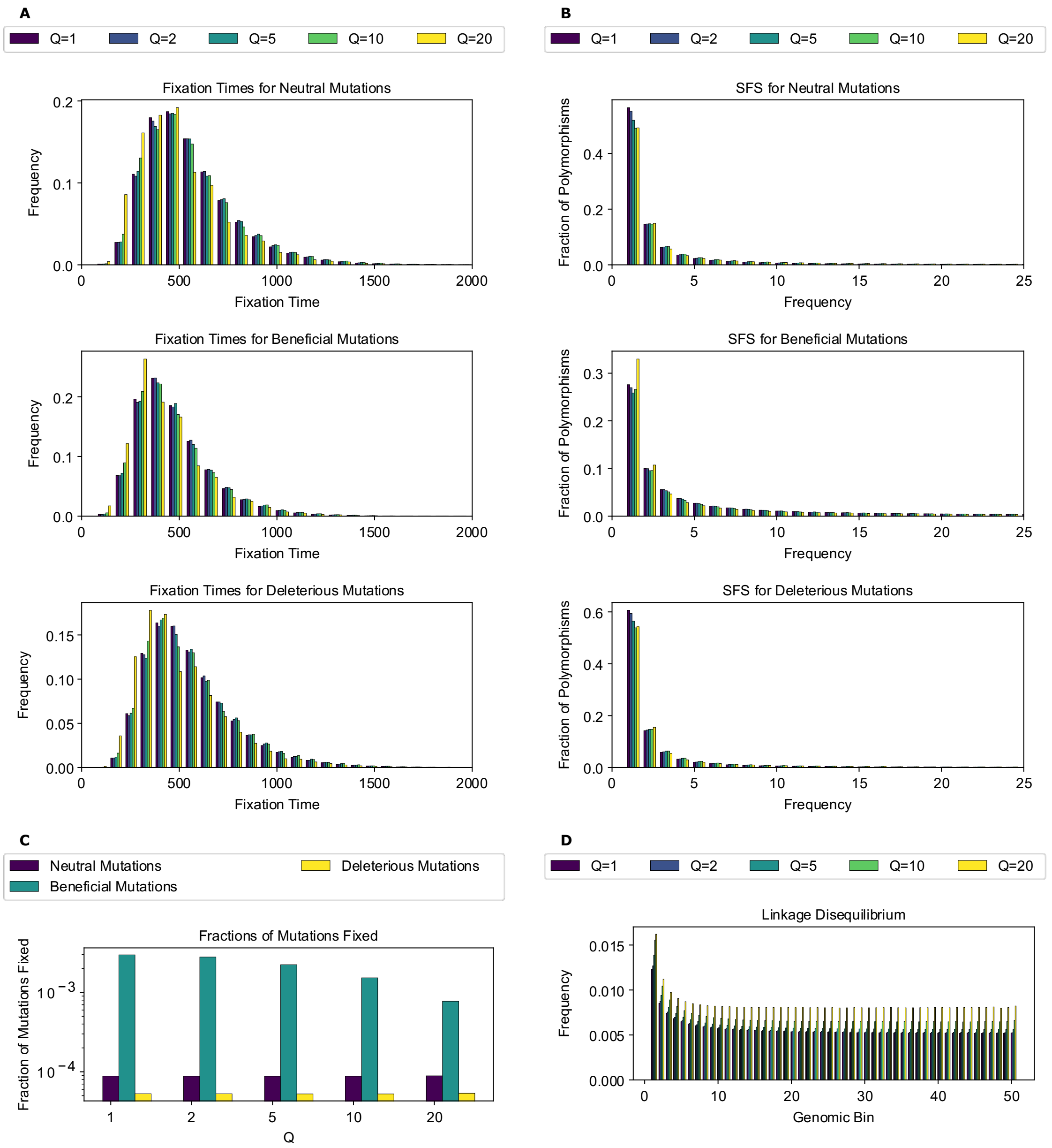


**Figure S13. Simulation outcomes for the population contraction model.**
**(A)** Fixation time distributions for all mutation types. **(B)** Site frequency spectra for all mutation types. **(C)** Fractions of mutations fixed for all mutation types. **(D)** Linkage disequilibrium across the chromosome for 50 genomic bins, as measured by $r^2$.

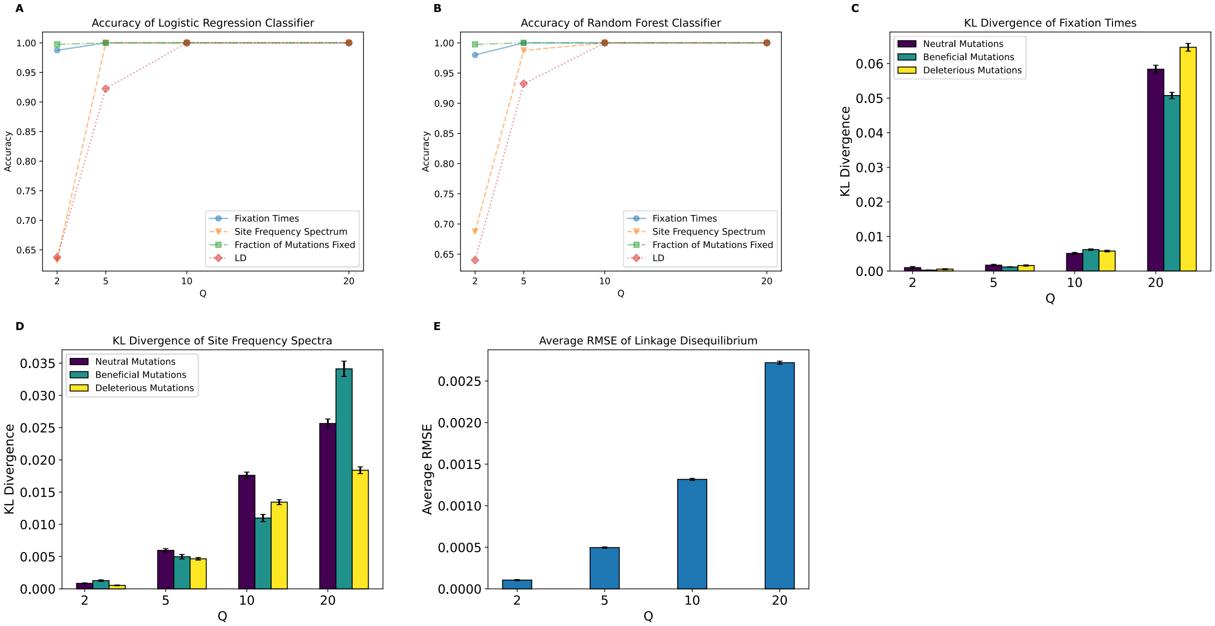


**Figure S14. Metrics of the effect of scaling for the population contraction model.**
**(A)** Accuracy of the logistic regression classification using features subsets for each scaling factor. **(B)** Accuracy of the random forest classification using feature subsets for each scaling factor. **(C)** KL divergence of fixation time distributions for all mutation types. **(D)** KL divergence for the SFS for all mutation types. **(E)** Average RMSE of LD. Values and errors bars for panels **(C)**, **(D)**, and **(E)**, represent mean and standard deviation of 1000 bootstrapped samples.

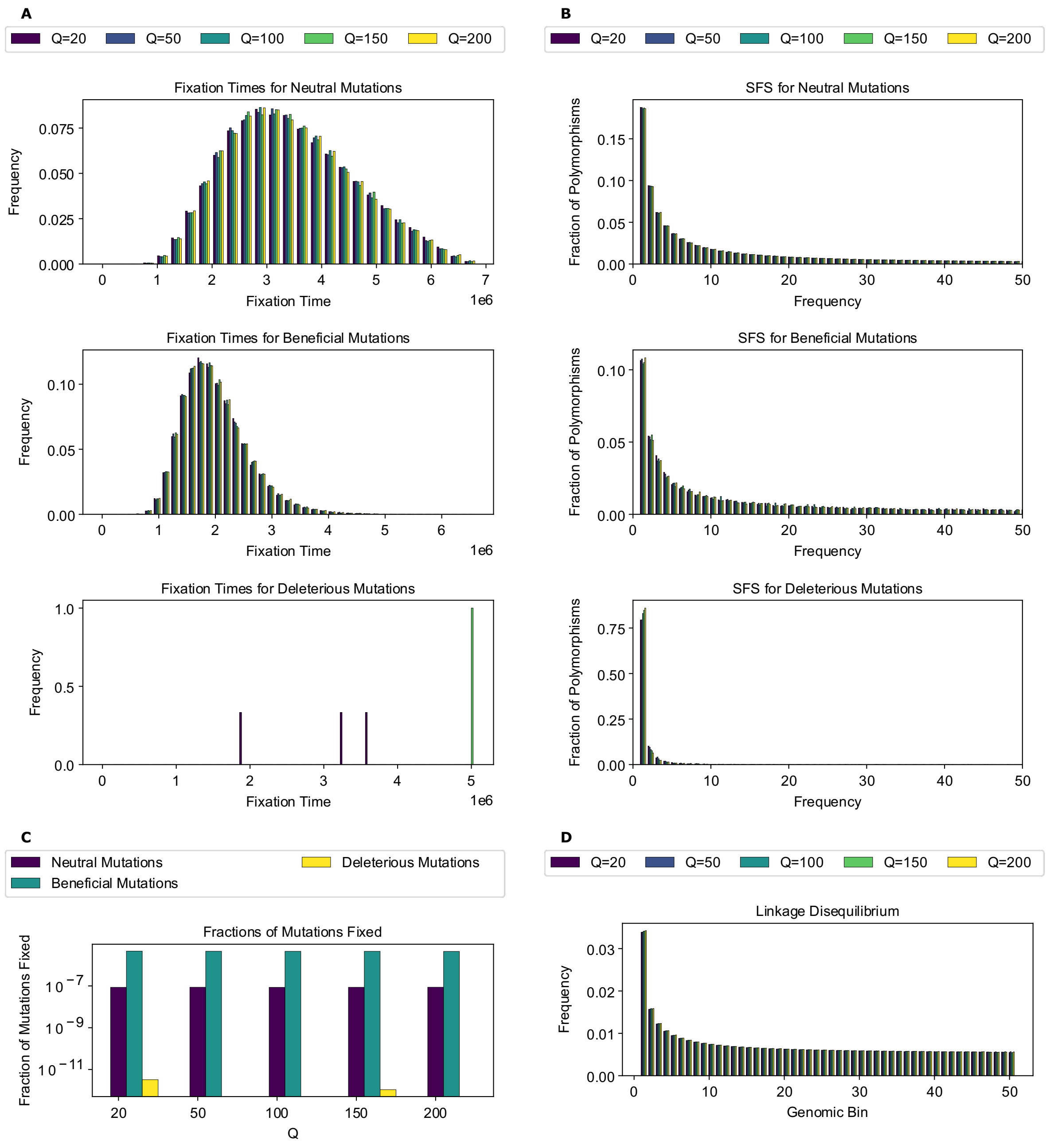


**Figure S15. Simulation outcomes for the *Drosophila* model.**
**(A)** Fixation time distributions for all mutation types. **(B)** Site frequency spectra for all mutation types. **(C)** Fractions of mutations fixed. for all mutation types. **(D)** Linkage disequilibrium across the chromosome for 50 genomic bins, as measured by $r^2$.

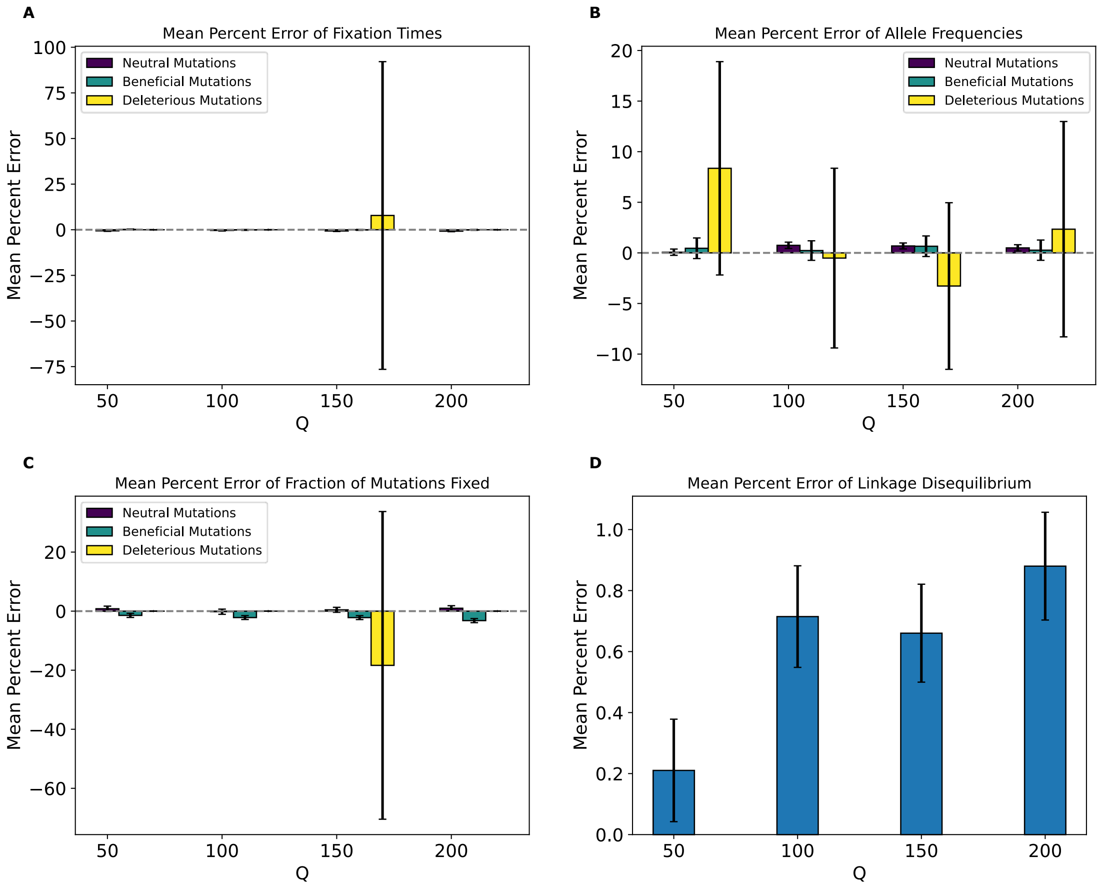


**Figure S16. Mean percent error of various statistics for the *Drosophila* model by values of *Q*.**
**(A)** Mean percent error for average mutation fixation times. **(B)** Mean percent error for average allele frequencies. **(C)** Mean percent error for average fractions of fixed mutations. **(D)** Mean percent error for average linkage disequilibrium as measured by the values of $r^2$. Values and errors bars represent mean and standard deviation of 1000 bootstrapped samples.

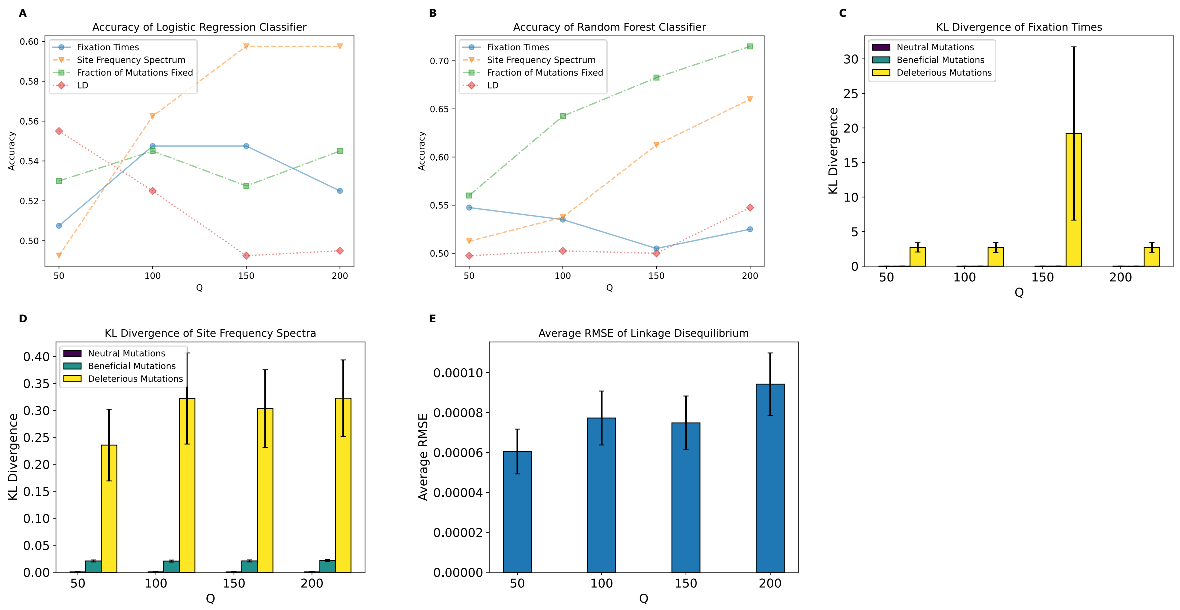


**Figure S17. Metrics of the effect of scaling for the *Drosophila* model.**
**(A)** Accuracy of the logistic regression classification using features subsets for each scaling factor. **(B)** Accuracy of the random forest classification using feature subsets for each scaling factor. **(C)** KL divergence of fixation time distributions for all mutation types. **(D)** KL divergence for the SFS for all mutation types. **(E)** Average RMSE of LD. Values and errors bars for panels **(C)**, **(D)**, and **(E)**, represent mean and standard deviation of 1000 bootstrapped samples.

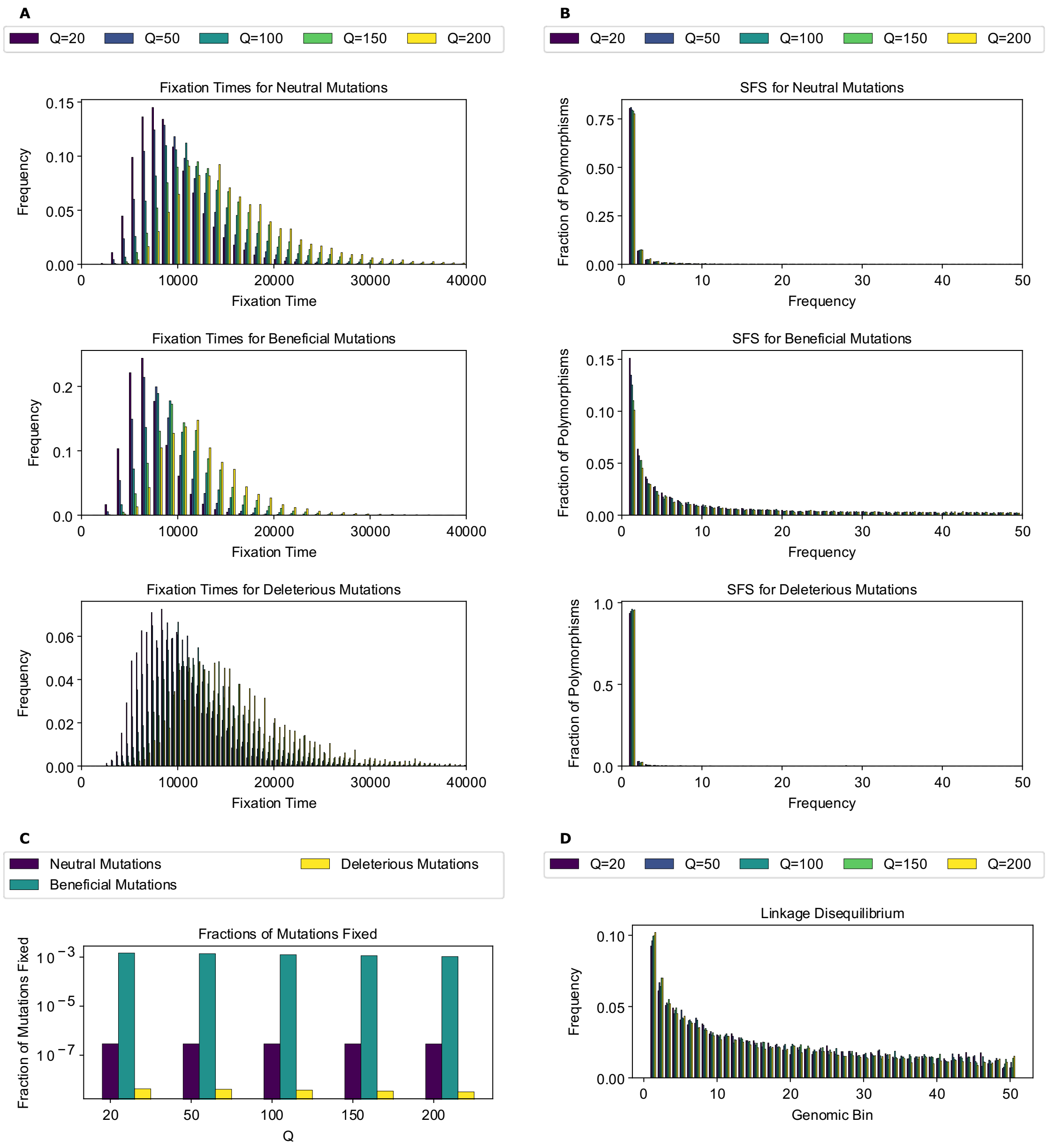


**Figure S18. Simulation outcomes for the *Drosophila* stronger selection model.**
**(A)** Fixation time distributions for all mutation types. **(B)** Site frequency spectra for all mutation types. **(C)** Fractions of mutations fixed for all mutation types. **(D)** Linkage disequilibrium across the chromosome for 50 genomic bins, as measured by $r^2$.

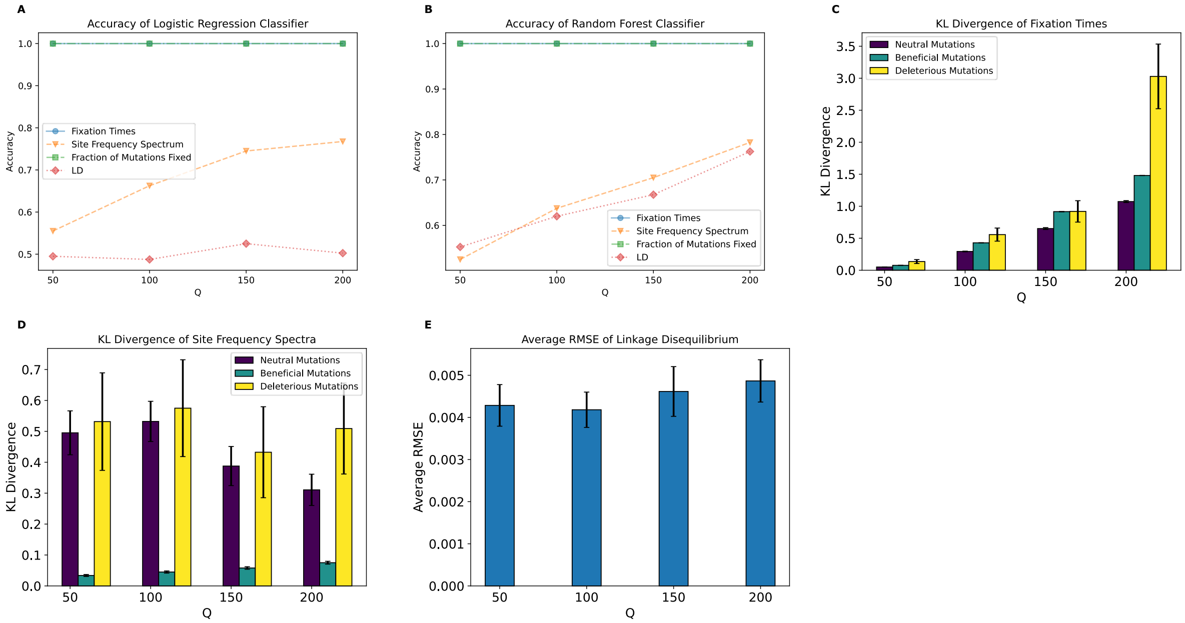


**Figure S19. Metrics of the effect of scaling for the *Drosophila* stronger selection model.**
**(A)** Accuracy of the logistic regression classification using features subsets for each scaling factor. **(B)** Accuracy of the random forest classification using feature subsets for each scaling factor. **(C)** KL divergence of fixation time distributions for all mutation types. **(D)** KL divergence for the SFS for all mutation types. **(E)** Average RMSE of LD. Values and errors bars for panels **(C)**, **(D)**, and **(E)**, represent mean and standard deviation of 1000 bootstrapped samples.

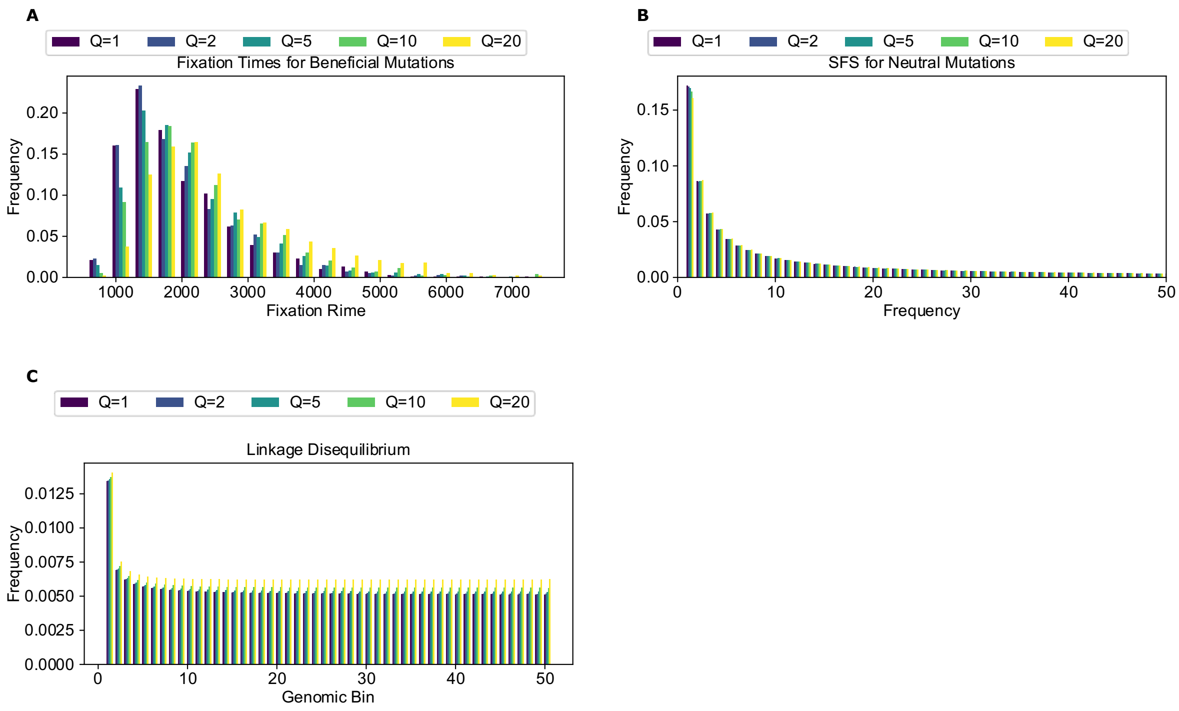


**Figure S20. Simulation outcomes for the sweep model.**
**(A)** Fixation time distributions for beneficial mutations. **(B)** Site frequency spectrum for neutral mutations. **(C)** Linkage disequilibrium for neutral mutations across the chromosome for 50 genomic bins, as measured by $r^2$.

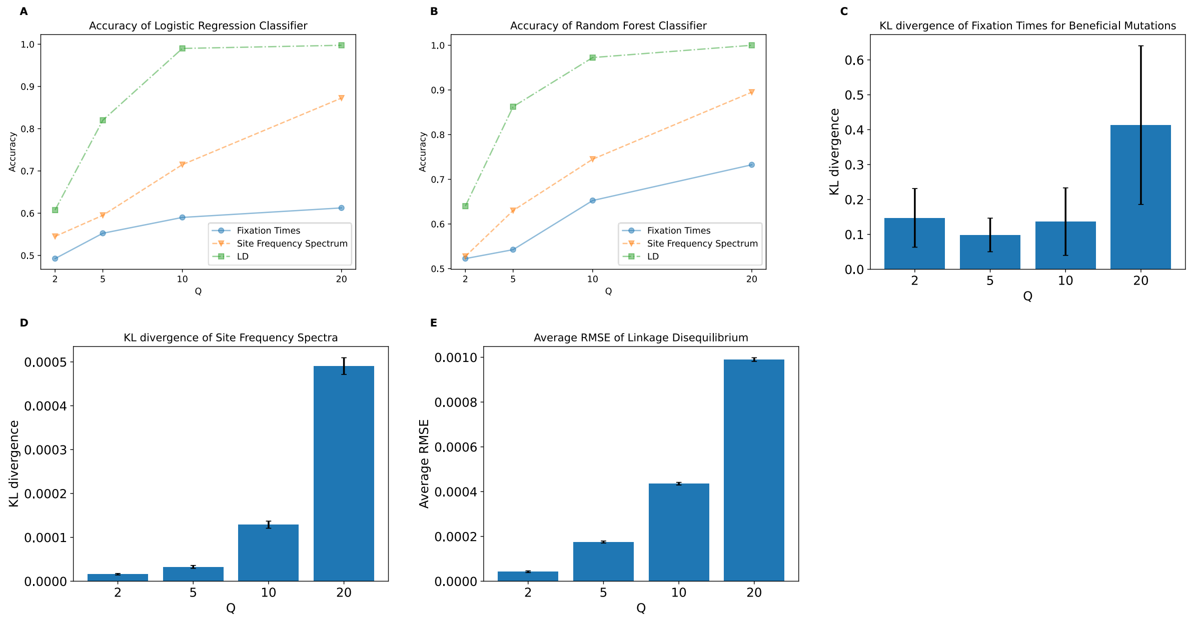


**Figure S21. Metrics of the effect of scaling for the sweep model.**
**(A)** Accuracy of the logistic regression classification using features subsets for each scaling factor. **(B)** Accuracy of the random forest classification using feature subsets for each scaling factor. **(C)** KL divergence of fixation time distributions for beneficial mutations. **(D)** KL divergence for the SFS for neutral mutations. **(E)** Average RMSE of LD for neutral mutations. Values and errors bars for panels **(C)**, **(D)**, and **(E)**, represent mean and standard deviation of 1000 bootstrapped samples.

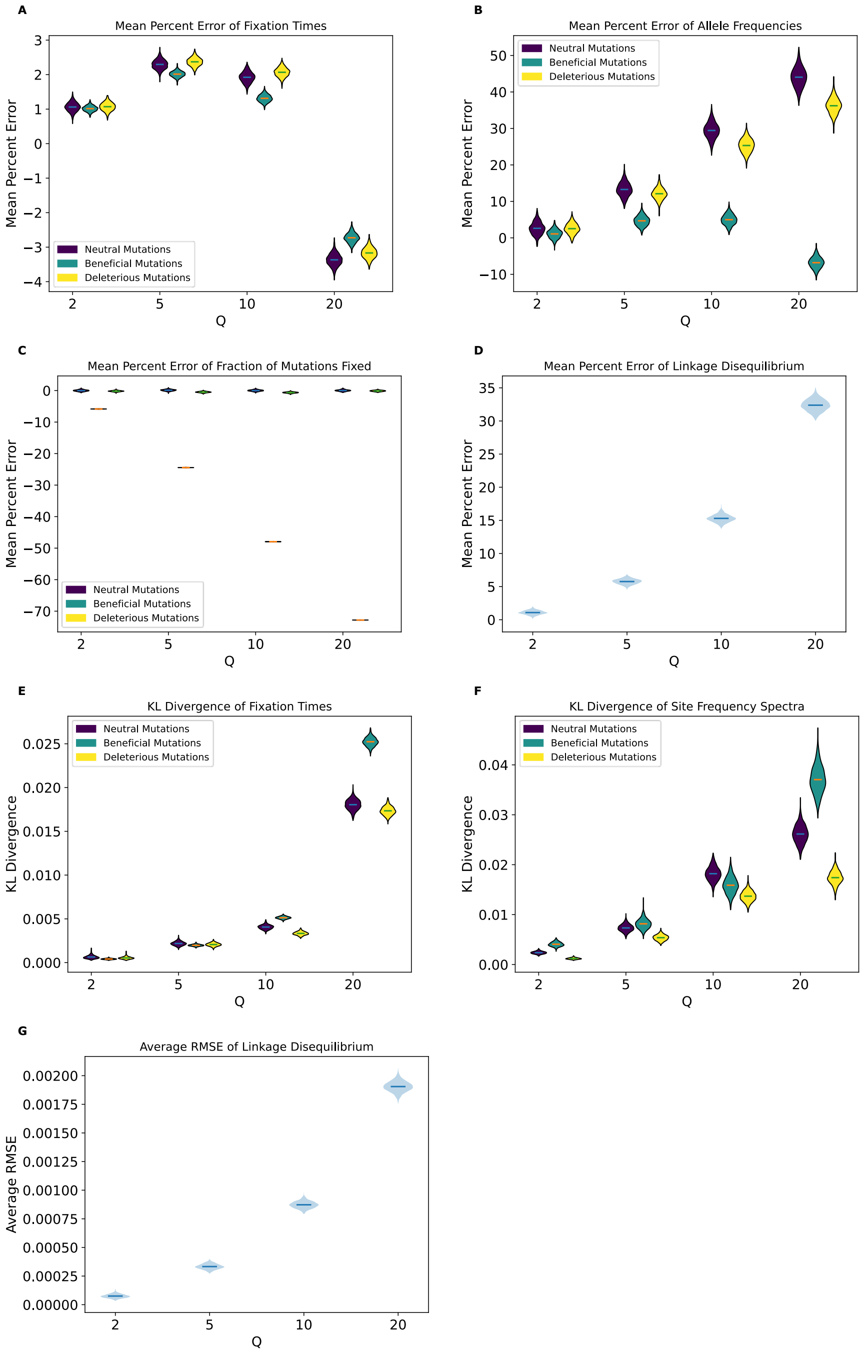


**Figure S22. Measures of deviation for the full model using only 100 randomly selected replicates.** **(A)** Mean percent error of average fixation times for all mutation types. **(B)** Mean percent error of average allele frequencies for all mutation types. **(C)** Mean percent error of average fractions of fixed mutations for all mutation types. **(D)** Mean percent error of average LD as measured by $r^2$. **(E)** KL Divergence of fixation times for all mutation types. **(F)** KL divergence of the SFS for all mutation types. **(G)** Average RMSE of LD as measured by $r^2$ across 50 genomic bins. Distributions are the result of repeating the sampling process 1000 times, with horizontal lines indicating the means of these distributions.

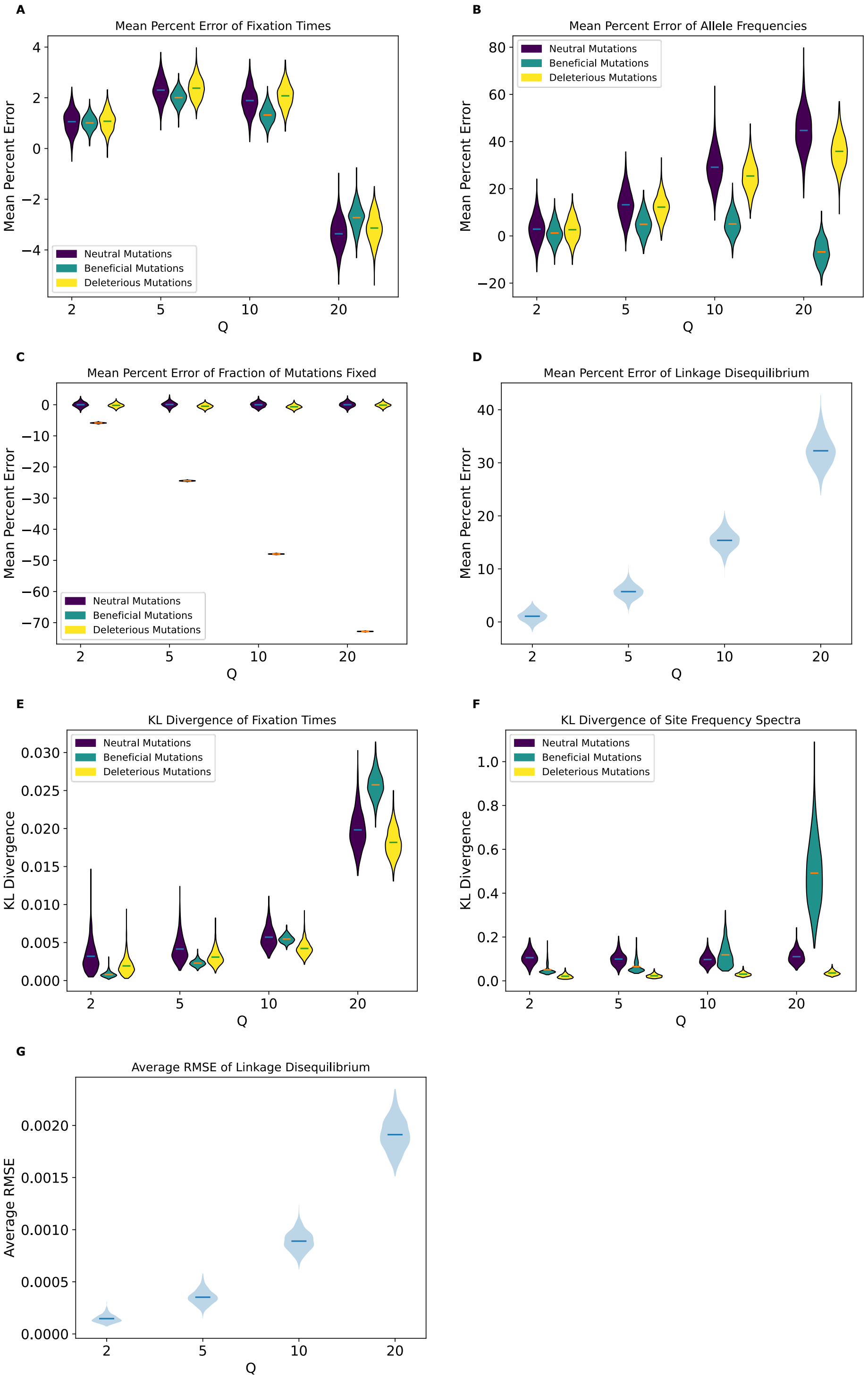


**Figure S23. Measures of deviation for the full model using only 10 randomly selected replicates.** **(A)** Mean percent error of average fixation times for all mutation types. **(B)** Mean percent error of average allele frequencies for all mutation types. **(C)** Mean percent error of average fractions of fixed mutations for all mutation types. **(D)** Mean percent error of average LD as measured by $r^2$. **(E)** KL Divergence of fixation times for all mutation types. **(F)** KL divergence of the SFS for all mutation types. **(G)** Average RMSE of LD as measured by $r^2$ across 50 genomic bins. Distributions are the result of repeating the sampling process 1000 times, with horizontal lines indicating the means of these distributions.

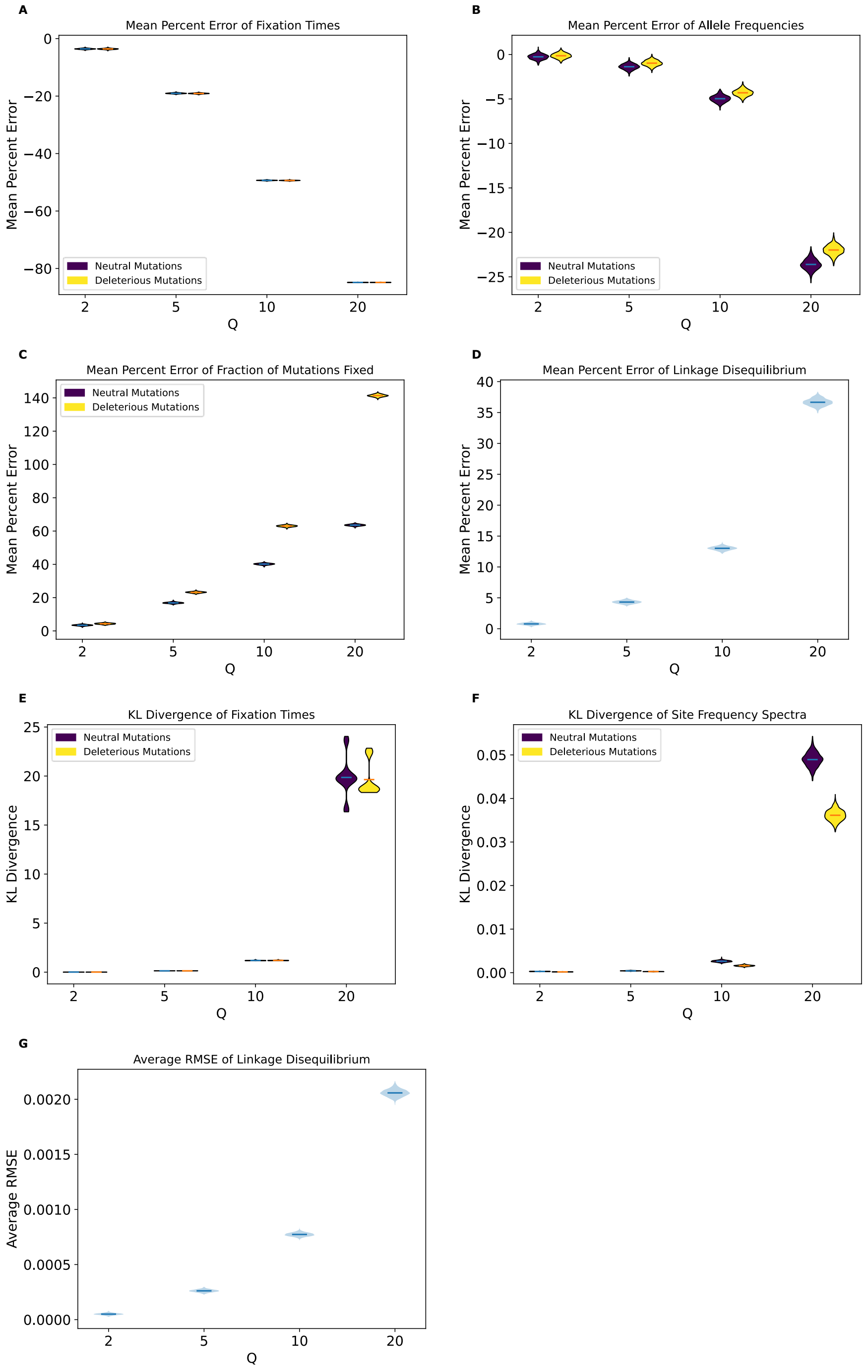


**Figure S24. Measures of deviation for the no-beneficials model using only 100 randomly selected replicates.** **(A)** Mean percent error of average fixation times for all mutation types. **(B)** Mean percent error of average allele frequencies for all mutation types. **(C)** Mean percent error of average fractions of fixed mutations for all mutation types. **(D)** Mean percent error of average LD as measured by $r^2$. **(E)** KL Divergence of fixation times for all mutation types. **(F)** KL divergence of the SFS for all mutation types. **(G)** Average RMSE of LD as measured by $r^2$ across 50 genomic bins. Distributions are the result of repeating the sampling process 1000 times, with horizontal lines indicating the means of these distributions.

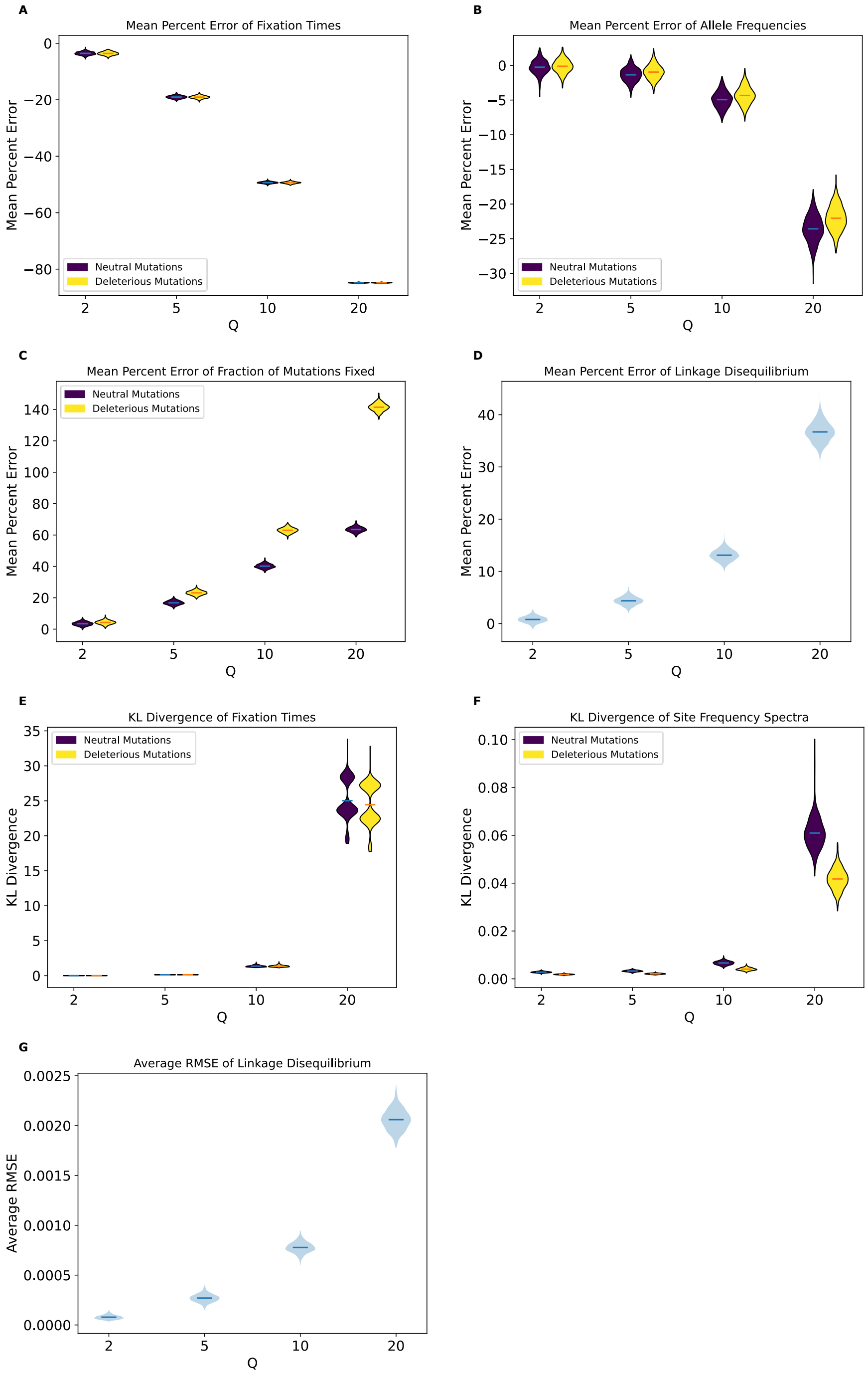


**Figure S25. Measures of deviation for the no-beneficials model using only 10 randomly selected replicates.** **(A)** Mean percent error of average fixation times for all mutation types. **(B)** Mean percent error of average allele frequencies for all mutation types. **(C)** Mean percent error of average fractions of fixed mutations for all mutation types. **(D)** Mean percent error of average LD as measured by $r^2$. **(E)** KL Divergence of fixation times for all mutation types. **(F)** KL divergence of the SFS for all mutation types. **(G)** Average RMSE of LD as measured by $r^2$ across 50 genomic bins. Distributions are the result of repeating the sampling process 1000 times, with horizontal lines indicating the means of these distributions.

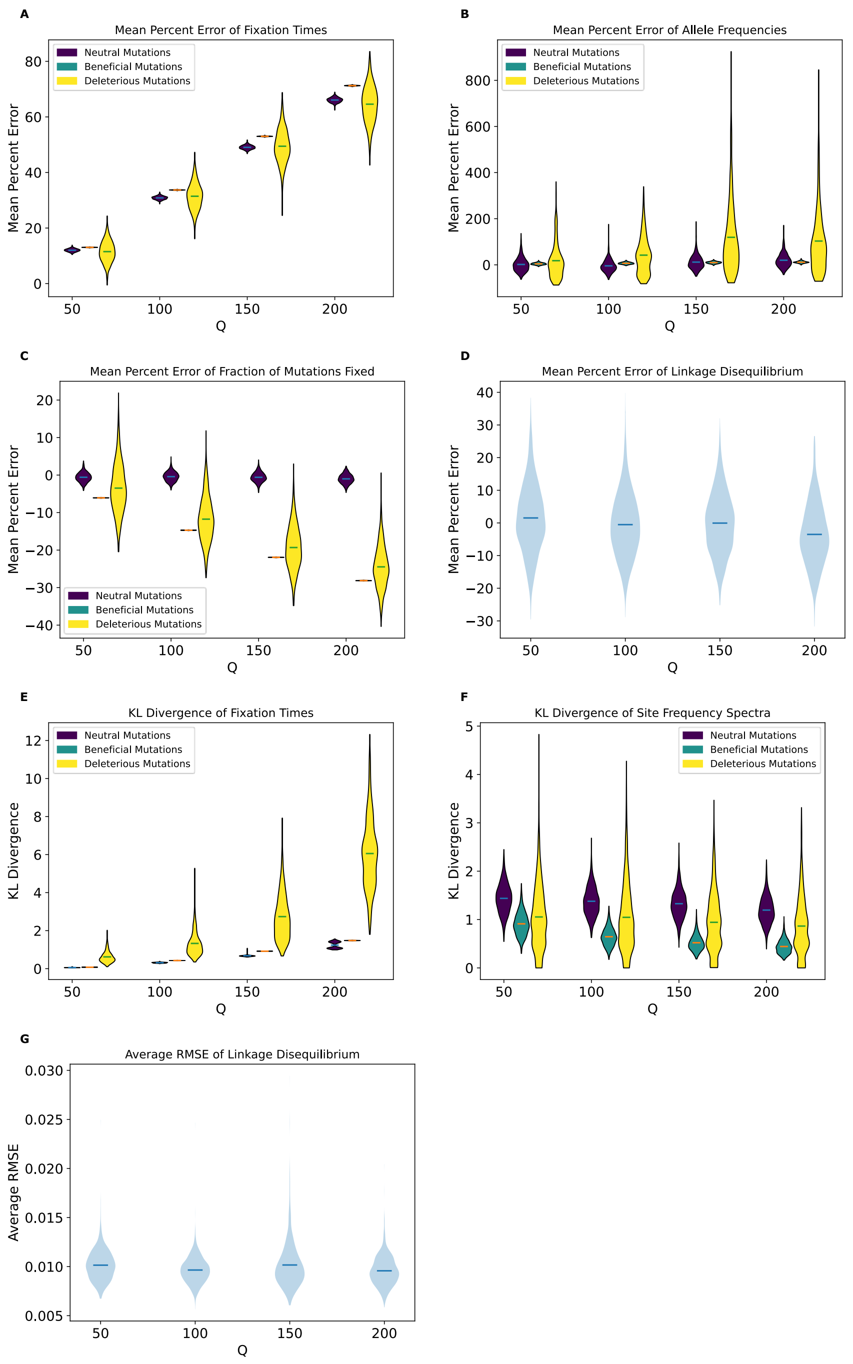


**Figure S26. Measures of deviation for the *Drosophila* stronger selection model using only 100 randomly selected replicates.**

**(A)** Mean percent error of average fixation times for all mutation types. **(B)** Mean percent error of average allele frequencies for all mutation types. **(C)** Mean percent error of average fractions of fixed mutations for all mutation types. **(D)** Mean percent error of average LD as measured by $r^2$. **(E)** KL Divergence of fixation times for all mutation types. **(F)** KL divergence of the SFS for all mutation types. **(G)** Average RMSE of LD as measured by $r^2$ across 50 genomic bins. Distributions are the result of repeating the sampling process 1000 times, with horizontal lines indicating the means of these distributions.

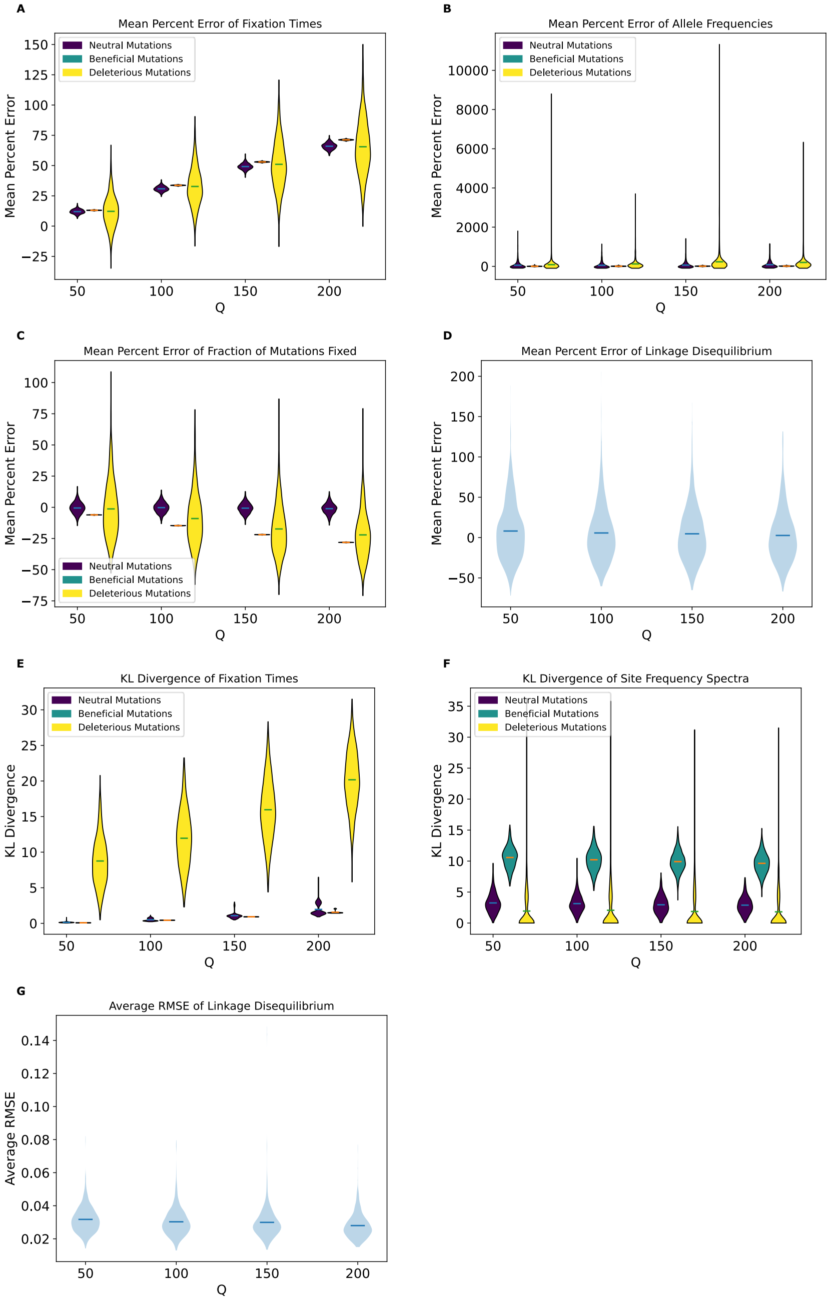


**Figure S27. Measures of deviation for the *Drosophila* stronger selection model using only 10 randomly selected replicates.**
**(A)** Mean percent error of average fixation times for all mutation types. **(B)** Mean percent error of average allele frequencies for all mutation types. **(C)** Mean percent error of average fractions of fixed mutations for all mutation types. **(D)** Mean percent error of average LD as measured by $r^2$. **(E)** KL Divergence of fixation times for all mutation types. **(F)** KL divergence of the SFS for all mutation types. **(G)** Average RMSE of LD as measured by $r^2$ across 50 genomic bins. Distributions are the result of repeating the sampling process 1000 times, with horizontal lines indicating the means of these distributions.

**Table S1. Values of KL Divergence (for Fixation Times and SFS) and Average RMSE (for LD) for All Simulation Models.**

| Model | Mutation Type | Outcome | Q=2 | Q=5 | Q=10 | Q=20 |
|---|---|---|---|---|---|---|
| **Full Model** | Neutral | Fixation Times | 0.00041 | 0.002 | 0.0039 | 0.018 |
| | | SFS | 0.00074 | 0.0057 | 0.017 | 0.025 |
| | Beneficial | Fixation Times | 0.00037 | 0.0019 | 0.0051 | 0.025 |
| | | SFS | 0.00099 | 0.0048 | 0.012 | 0.031 |
| | Deleterious | Fixation Times | 0.00039 | 0.002 | 0.0032 | 0.017 |
| | | SFS | 0.00044 | 0.0047 | 0.013 | 0.017 |
| | All | Linkage Disequilibrium | 0.0000663 | 0.00033 | 0.00087 | 0.0019 |
| **No-Beneficials** | Neutral | Fixation Times | 0.0045 | 0.13 | 1.17 | 17.2 |
| | | SFS | 0.0000636 | 0.00017 | 0.0023 | 0.048 |
| | Deleterious | Fixation Times | 0.0044 | 0.13 | 1.18 | 18.7 |
| | | SFS | 0.0000388 | 0.000089 | 0.0014 | 0.036 |
| | All | Linkage Disequilibrium | 0.0000478 | 0.00026 | 0.00077 | 0.0021 |
| **No-Beneficials Sparse** | Neutral | Fixation Times | 0.0000418 | 0.00066 | 0.0054 | 0.04 |
| | | SFS | 0.0000141 | 3.14E-05 | 8.45E-05 | 0.00028 |
| | Deleterious | Fixation Times | 0.0013 | 0.0021 | 0.008 | 0.04 |
| | | SFS | 0.00028 | 0.00029 | 0.0004 | 0.00068 |
| | All | Linkage Disequilibrium | 0.0000465 | 0.00018 | 0.00045 | 0.0011 |
| **No-Beneficials Larger Population** | Neutral | Fixation Times | 0.0045 | 0.14 | 1.22 | 25.3 |
| | | SFS | 0.00033 | 0.00037 | 0.0017 | 0.04 |
| | Deleterious | Fixation Times | 0.0045 | 0.14 | 1.23 | 22.5 |
| | | SFS | 0.0002 | 0.00023 | 0.0012 | 0.03 |
| | All | Linkage Disequilibrium | 0.0000521 | 0.00019 | 0.00053 | 0.0015 |
| **Strictly Neutral** | Neutral | Fixation Times | 0.000015 | 1.49E-05 | 2.23E-05 | 2.77E-05 |
| | | SFS | 0.000012 | 2.86E-05 | 9.62E-05 | 0.00047 |
| | Neutral | Linkage Disequilibrium | 0.0000314 | 0.00017 | 0.00043 | 0.00099 |
| **Population Expansion** | Neutral | Fixation Times | 0.00043 | 0.003 | 0.0056 | 0.0081 |
| | | SFS | 0.0007 | 0.0051 | 0.015 | 0.023 |
| | Beneficial | Fixation Times | 0.0005 | 0.0027 | 0.0048 | 0.0066 |
| | | SFS | 0.00092 | 0.0047 | 0.012 | 0.026 |
| | Deleterious | Fixation Times | 0.00052 | 0.0028 | 0.0059 | 0.012 |
| | | SFS | 0.00051 | 0.0042 | 0.011 | 0.016 |
| | All | Linkage Disequilibrium | 0.0000364 | 0.00022 | 0.00061 | 0.0014 |
| **Population Contraction** | Neutral | Fixation Times | 0.00098 | 0.0017 | 0.0051 | 0.058 |
| | | SFS | 0.00085 | 0.006 | 0.018 | 0.026 |
| | Beneficial | Fixation Times | 0.00029 | 0.0012 | 0.0062 | 0.051 |
| | | SFS | 0.0013 | 0.005 | 0.011 | 0.034 |
| | Deleterious | Fixation Times | 0.00056 | 0.0016 | 0.0058 | 0.065 |
| | | SFS | 0.00054 | 0.0047 | 0.013 | 0.018 |
| | All | Linkage Disequilibrium | 0.00011 | 0.0005 | 0.0013 | 0.0027 |
| **Sweep** | Neutral | SFS | 1.60E-05 | 3.26E-05 | 1.29E-04 | 4.92E-04 |

| | | | | | | |
|---|---|---|---|---|---|---|
| | | Linkage Disequilibrium | 4.23E-05 | 1.75E-04 | 4.36E-04 | 9.89E-04 |
| | Beneficial | Fixation Times | 0.14 | 0.096 | 0.14 | 0.41 |
| | | | **Q=50** | **Q=100** | **Q=150** | **Q=200** |
| ***Drosophila*** | Neutral | Fixation Times | 0.0025 | 0.0026 | 0.0044 | 0.0045 |
| | | SFS | 0.00077 | 0.00081 | 0.00074 | 0.00076 |
| | Beneficial | Fixation Times | 0.0054 | 0.0052 | 0.0075 | 0.0036 |
| | | SFS | 0.021 | 0.021 | 0.021 | 0.021 |
| | Deleterious | Fixation Times | 2.66 | 2.68 | 18.8 | 2.69 |
| | | SFS | 0.24 | 0.33 | 0.3 | 0.32 |
| | All | Linkage Disequilibrium | 0.0000605 | 7.77E-05 | 7.49E-05 | 9.47E-05 |
| ***Drosophila* Stronger Selection** | Neutral | Fixation Times | 0.051 | 0.29 | 0.65 | 1.07 |
| | | SFS | 0.5 | 0.54 | 0.39 | 0.31 |
| | Beneficial | Fixation Times | 0.077 | 0.43 | 0.92 | 1.48 |
| | | SFS | 0.034 | 0.045 | 0.058 | 0.076 |
| | Deleterious | Fixation Times | 0.14 | 0.57 | 0.92 | 3.02 |
| | | SFS | 0.53 | 0.57 | 0.43 | 0.51 |
| | All | Linkage Disequilibrium | 0.0043 | 0.0042 | 0.0046 | 0.0049 |

**Table S2. Classifier Accuracy for All Simulation Models.**

| Model | Outcome | Classification Method | Q=2 | Q=5 | Q=10 | Q=20 |
|---|---|---|---|---|---|---|
| **Full Model** | Fixation Times | Logistic Regression | 100% | 100% | 100% | 100% |
| | | Random Forest | 100% | 100% | 100% | 100% |
| | SFS | Logistic Regression | 72% | 100% | 100% | 100% |
| | | Random Forest | 73% | 97% | 100% | 100% |
| | Fraction of Mutations Fixed | Logistic Regression | 100% | 100% | 100% | 100% |
| | | Random Forest | 100% | 100% | 100% | 100% |
| | Linkage Disequilibrium | Logistic Regression | 60% | 90% | 100% | 100% |
| | | Random Forest | 60% | 91% | 100% | 100% |
| **No-beneficials** | Fixation Times | Logistic Regression | 96% | 100% | 100% | 100% |
| | | Random Forest | 97% | 100% | 100% | 100% |
| | SFS | Logistic Regression | 91% | 100% | 100% | 100% |
| | | Random Forest | 91% | 100% | 100% | 100% |
| | Fraction of Mutations Fixed | Logistic Regression | 79% | 100% | 100% | 100% |
| | | Random Forest | 81% | 100% | 100% | 100% |
| | Linkage Disequilibrium | Logistic Regression | 66% | 95% | 100% | 100% |
| | | Random Forest | 67% | 96% | 100% | 100% |
| **No-beneficials Larger Population** | Fixation Times | Logistic Regression | 100% | 100% | 100% | 100% |
| | | Random Forest | 100% | 100% | 100% | 100% |
| | SFS | Logistic Regression | 100% | 100% | 100% | 100% |
| | | Random Forest | 100% | 100% | 100% | 100% |
| | Fraction of Mutations Fixed | Logistic Regression | 88% | 100% | 100% | 100% |
| | | Random Forest | 85% | 100% | 100% | 100% |
| | Linkage Disequilibrium | Logistic Regression | 63% | 98% | 100% | 100% |
| | | Random Forest | 58% | 100% | 100% | 100% |
| **No-beneficials Sparse** | Fixation Times | Logistic Regression | 54% | 76% | 98% | 100% |
| | | Random Forest | 56% | 74% | 99% | 100% |
| | SFS | Logistic Regression | 54% | 79% | 100% | 100% |
| | | Random Forest | 55% | 88% | 100% | 100% |
| | Fraction of Mutations Fixed | Logistic Regression | 61% | 72% | 95% | 100% |
| | | Random Forest | 55% | 72% | 97% | 100% |
| | Linkage Disequilibrium | Logistic Regression | 57% | 83% | 99% | 100% |
| | | Random Forest | 59% | 85% | 99% | 100% |
| **Strictly Neutral** | Fixation Times | Logistic Regression | 47% | 49% | 48% | 52% |
| | | Random Forest | 49% | 50% | 53% | 52% |
| | SFS | Logistic Regression | 52% | 55% | 72% | 89% |
| | | Random Forest | 48% | 60% | 73% | 87% |
| | Fraction of Mutations Fixed | Logistic Regression | 43% | 47% | 47% | 53% |
| | | Random Forest | 52% | 54% | 56% | 55% |
| | Linkage Disequilibrium | Logistic Regression | 55% | 84% | 99% | 100% |
| | | Random Forest | 59% | 83% | 97% | 100% |
| **Population Expansion** | Fixation Times | Logistic Regression | 100% | 100% | 100% | 100% |
| | | Random Forest | 100% | 100% | 100% | 100% |
| | SFS | Logistic Regression | 72% | 100% | 100% | 100% |
| | | Random Forest | 66% | 98% | 100% | 100% |
| | Fraction of Mutations Fixed | Logistic Regression | 100% | 100% | 100% | 100% |

| | | | | | | |
|---|---|---|---|---|---|---|
| | | Random Forest | 100% | 100% | 100% | 100% |
| | Linkage Disequilibrium | Logistic Regression | 56% | 85% | 100% | 100% |
| | | Random Forest | 58% | 82% | 99% | 100% |
| **Population Contraction** | Fixation Times | Logistic Regression | 99% | 100% | 100% | 100% |
| | | Random Forest | 98% | 100% | 100% | 100% |
| | SFS | Logistic Regression | 63% | 100% | 100% | 100% |
| | | Random Forest | 69% | 99% | 100% | 100% |
| | Fraction of Mutations Fixed | Logistic Regression | 100% | 100% | 100% | 100% |
| | | Random Forest | 100% | 100% | 100% | 100% |
| | Linkage Disequilibrium | Logistic Regression | 64% | 92% | 100% | 100% |
| | | Random Forest | 64% | 93% | 100% | 100% |
| **Sweep** | Fixation Times | Logistic Regression | 49% | 55% | 59% | 61% |
| | | Random Forest | 52% | 54% | 65% | 73% |
| | SFS | Logistic Regression | 55% | 60% | 72% | 87% |
| | | Random Forest | 53% | 63% | 75% | 90% |
| | Linkage Disequilibrium | Logistic Regression | 61% | 82% | 99% | 100% |
| | | Random Forest | 64% | 86% | 97% | 100% |
| | | | **Q=50** | **Q=100** | **Q=150** | **Q=200** |
| ***Drosophila*** | Fixation Times | Logistic Regression | 51% | 55% | 55% | 53% |
| | | Random Forest | 55% | 54% | 51% | 53% |
| | SFS | Logistic Regression | 49% | 56% | 60% | 60% |
| | | Random Forest | 51% | 54% | 61% | 66% |
| | Fraction of Mutations Fixed | Logistic Regression | 53% | 55% | 53% | 55% |
| | | Random Forest | 56% | 64% | 68% | 72% |
| | Linkage Disequilibrium | Logistic Regression | 56% | 53% | 49% | 50% |
| | | Random Forest | 50% | 50% | 50% | 55% |
| ***Drosophila* Stronger Selection** | Fixation Times | Logistic Regression | 100% | 100% | 100% | 100% |
| | | Random Forest | 100% | 100% | 100% | 100% |
| | SFS | Logistic Regression | 56% | 66% | 75% | 77% |
| | | Random Forest | 53% | 64% | 71% | 78% |
| | Fraction of Mutations Fixed | Logistic Regression | 100% | 100% | 100% | 100% |
| | | Random Forest | 100% | 100% | 100% | 100% |
| | Linkage Disequilibrium | Logistic Regression | 50% | 49% | 53% | 50% |
| | | Random Forest | 55% | 62% | 67% | 76% |

**Table S3. Simulation Runtimes (in Minutes) for Select Simulation Models.**

| Model | Q=1 | Q=2 | Q=5 | Q=10 | Q=20 |
|---|---|---|---|---|---|
| **Full Model** | 308 | 100 | 22 | 6 | 1.4 |
| **No-beneficials Larger Population** | 5645 | 1929 | 433 | 109 | 20.5 |
| **Strictly Neutral** | 493 | 173 | 48 | 18 | 7.7 |
| **Population Expansion** | 408 | 122 | 24 | 6 | 1.6 |
| **Population Contraction** | 208 | 57 | 12 | 3 | 0.7 |
| | **Q=20** | **Q=50** | **Q=100** | **Q=150** | **Q=200** |
| ***Drosophila*** | 1458 | 311 | 78 | 32 | 23 |
| ***Drosophila* Stronger Selection** | 1383 | 201 | 43 | 26 | 12 |
